# Cell-resolved high-dimensional imaging analysis, therapeutic modeling, and a Phase Ib clinical study establish BCL-2 as a target across heterogeneous CRPC subtypes

**DOI:** 10.1101/2025.07.08.663739

**Authors:** Anmbreen Jamroze, Xiaozhuo Liu, Surui Hou, Wen (Jess) Li, Han Yu, Amanda Tracz, Qiuhui Li, Kent Nastiuk, Xin Chen, Jiaoti Huang, Kevin Lin, Yue Lu, Igor Puzanov, Jason S. Kirk, Gurkamal Chatta, Dean G. Tang

## Abstract

BCL-2 has been implicated in prostate cancer (PCa) progression and development of castration-resistant disease (CRPC). However, it remains unclear how the BCL-2- and AR-expressing PCa cell populations evolve across the PCa continuum, how AR molecularly regulates BCL-2 and whether BCL-2 represents a common therapeutic target in heterogeneous CRPC. Importantly, BCL-2 inhibitors have yet to be approved for treating PCa patients. Here we first show the selective induction of BCL-2 by AR pathway inhibitors (ARPIs) in both patient specimens and xenograft models. Vectra-based quantitative multiplex immunofluorescence (qmIF) and image mass cytometry (IMC) analyses with single-cell resolution reveal markedly expanded BCL-2^+^ (AR^+^ or AR^-^) PCa cell populations in CRPC. Mechanistically, AR represses *BCL-2* transcription through genomic binding via several AR binding sites and ARPIs relieve this repression, leading to BCL-2 upregulation. Comprehensive therapeutic studies in cells, organoids and xenografts establish that castration-induced BCL-2 is not merely associated with resistance but represents a shared and actionable vulnerability as the BCL-2 inhibitor ABT-199 potently suppressed the growth of multiple subtypes of CRPC. A Phase Ib clinical trial (NCT03751436) combining enzalutamide and BCL-2 inhibitor venetoclax demonstrated reduced circulating tumor cells in responding patients. Together, our findings elucidate the AR^+/-^BCL-2^+/-^ PCa cell subpopulation dynamics during PCa progression, reveal a direct mechanistic link between AR inhibition and BCL-2–mediated resistance, and provide a strong rationale for targeting BCL-2 from the outset to eliminate emerging resistant subpopulations, inhibit treatment-induced cellular heterogeneity and plasticity, and improve therapeutic outcomes in CRPC.

## INTRODUCTION

Prostate cancer (PCa) continues to claim a high mortality with >35,000 American men estimated to die from metastatic castration-resistant PCa (mCRPC) in 2025.^1^ The incidence rates of advanced PCa have increased significantly and the proportion of men diagnosed with distant metastases has doubled from 2011 to 2019 when the second-line androgen receptor (AR) pathway inhibitors (ARPIs) such as abiraterone and enzalutamide (Enza) were introduced to the clinic. Recent studies have linked both the increased diagnosis of advanced and metastatic PCa and persistently high PCa mortality to therapy resistance driven by PCa cell heterogeneity and plasticity.^2–8^

Advanced PCa is treated with androgen deprivation therapy (ADT) with or without ARPIs. ADT/Enza are effective in targeting AR-expressing (AR^+^) PCa cells; however, PCa at all stages, from newly diagnosed primary tumor to mCRPC, harbors AR^-/lo^ PCa cells expressing little or no AR,^4,5,9–23^ which we have shown to be *de novo* refractory to Enza.^20^ Moreover, the AR^-/lo^(PSA^-/lo^) PCa cells pre-exist in treatment-naïve tumors and exhibit gene expression profiles and functional properties of PCa stem cells (PCSCs).^5,14–18,22,24–26^ CRPC is often characterized with increased AR^-/lo^ PCa cells, which may stem from an expansion of pre-existing AR^-/lo^ population and/or from ARPI-induced reprogramming (plasticity or de-differentiation) of AR^+^ to AR^-/lo^ cells.^5,8,14,15,17,20,21,23^ ARPIs induce PCa cell plasticity via several inter-woven mechanisms involving genetic mutations, epigenetic plasticity, and transcriptional and metabolic re-wiring.^8^ In principle, to achieve enduring clinical efficacy, both cancer cell heterogeneity and therapy-induced plasticity and both AR^+^ and AR^-/lo^ PCa cell populations should be therapeutically targeted.^5^

We have recently reported, in experimental and patient CRPC, three CRPC subtypes with distinct AR expression levels and patterns, i.e., AR^+/hi^ (high nuclear AR), AR^-/lo^ (minimal or absent AR) and AR^cyto^ (predominantly cytoplasmic AR).^20^ These three CRPC subtypes manifest distinct responses to Enza: AR^+/hi^ LNCaP-CRPC exhibits an acute and transient sensitivity and AR^cyto^ LAPC4-CRPC and VCaP-CRPC a more drawn-out sensitivity whereas AR^-/lo^ LAPC9-CRPC shows no response to Enza.^20^ Importantly, this study nominated BCL-2 as a potential molecular driver and therapeutic target for CRPC.^20^

BCL-2 is a well-studied prosurvival and stemness factor and has been implicated in progression and therapy resistance of many blood and solid cancers including PCa.^27–50^ Early studies from our lab have shown the preferential expression of BCL-2 in AR^-/lo^ PCSCs,^14,17^ critical prosurvival functions of BCL-2 in PCa cells under nutrient deprivation and oxidative and genotoxic stresses,^29,32^ and induction of BCL-2 in PCa models by chemotherapeutic drugs, castration and castration/Enza.^20,29,32,37,43^ Consistently, castration, ARPIs, chemotherapeutic drugs, prooxidants and radiation have all been reported to induce BCL-2, which in turn promotes PCa cell survival and resistance to such agents.^20,27–29,32,35,37,40,42,43,45–50^ Therefore, BCL-2 has long been considered a therapeutic target in PCa, and several types of BCL-2-targeting therapeutics have been developed for the clinic, including antisense oligonucleotides (ASO; oblimersen sodium),^51–55^ siRNA,^56^ Gossypol (a polyphenol extract from cottonseed) and its derivative AT-101 (R-(-)-gossypol acetic acid),^57–58^ and BH3 mimetics such as HA14-1,^59^ WL-276,^60^ Venetoclax (ABT-199),^61–64^ and recently, Lisaftoclax (APG-2575)^65–67^ and Sonrotocalx (BGB-11417)^68–69^.

Despite these developments, BCL-2 inhibitors have not yet reached the clinic for PCa, and significant knowledge gaps remain. For example, we lack a holistic understanding of the dynamic changes in, and interrelationship between, BCL-2- and AR-expressing PCa cells across the PCa evolutionary continuum; it is incompletely understood how AR regulates BCL-2 under different androgen conditions and how ARPIs trigger the upregulation of BCL-2; and it remains unclear whether BCL-2 represents a common therapeutic target across diverse CRPC subtypes such as AR^+/hi^, AR^cyto^ and AR^-/lo^ CRPC.^20^ Herein, we address these knowledge gaps by employing human PCa specimens and *in vivo* and *in vitro* models of CRPC subtypes, combined with high-content quantitative imaging analyses, mechanistic studies, and therapeutic evaluations using organoid and xenograft systems. These investigations validate BCL-2 as a critical therapeutic target in heterogeneous CRPC. In addition, we present correlative data from a Phase Ib clinical trial^70^ combining Enza with Venetoclax in mCRPC patients (NCT03751436), showing that this combination reduced circulating tumor cells (CTCs) in several ‘responsive’ patients, underscoring the translational relevance of this therapeutic strategy.

## RESULTS

### BCL-2 is selectively and consistently induced by castration in patient PCa and xenograft models

We first examined a transcriptomic dataset^71^ for *BCL-2* mRNA levels in 3 epithelial cell (CD45^-^EpCAM^+^) populations of the normal human prostate, i.e., basal (B; CD49f^hi^CD38^lo^), luminal (L; CD49f^lo^CD26^+^CD38^hi^) and luminal progenitor (LP; CD49f^lo^CD26^+^CD38^lo^) cells. We found that *BCL-2* mRNA was expressed at high levels in LP and basal cells but had lower expression in mature luminal cells (Fig. 1a). We then examined *BCL-2* levels in 422 untreated primary tumors in TCGA-PRAD, which displayed an increasing trend that correlated with tumor grade (i.e., combined Gleason Score, GS) as supported by Jonckhere-Terpstra (J-T) trend test (Fig. 1b). We subsequently investigated the mRNA levels of all 5 prosurvival BCL-2 family members, i.e., BCL-2, BCL-xL, MCL-1, BCL-W and A1/BFL-1, in 4 treated patient datasets including 3 datasets of PCa patients subjected to neoadjuvant ADT (nADT; short-term (∼2-6 months) ADT prior to prostatectomy)^72,73,unpublished^ and 1 dataset of mCRPC patients who failed long-term ADT/Enza^23^. As shown in Fig. 1c, *BCL-2* was the only family member that was commonly and consistently induced in the 3 nADT datasets and showed a trend of upregulation in Enza-resistant PCa. Finally, we determined the mRNA and protein levels of BCL-2 family members in our 4 castration-resistant (i.e., androgen-independent; AI) xenograft models (i.e., LNCaP, LAPC9, VCaP, and LAPC4) developed by serially passaging the parental androgen-dependent (AD) tumors in castrated mice^14,18,20,24–26^ (Fig. 1d-e; also see Supplementary Fig. S6, below). We observed that *BCL-2* mRNA was commonly and exclusively upregulated in castration-resistant (primary CRPC), castration/Enza-resistant (secondary CRPC) and AR-knockout (ARKO) LNCaP models as well as in AR^-/lo^ LAPC9 CRPC^20^ (Fig. 1e) whereas BCL-2 protein was upregulated in 3 of the 4 CRPC (i.e., AI) models except VCaP (Fig. 1d). In contrast, the other 4 prosurvival BCL-2 family proteins showed model-dependent changes. For example, BCL-xL was slightly upregulated in VCaP CRPC where BCL-2, A1/BF-L1 and MCL-1 were all downregulated (Fig. 1d). BCL-W was reduced in 3 (LAPC9, LNCaP and LAPC4) xenograft CRPC whereas A1/BF-L1 and MCL-1 were both decreased in LAPC9, LNCaP and VCaP CRPC (Fig 1d).

**Figure 1.**
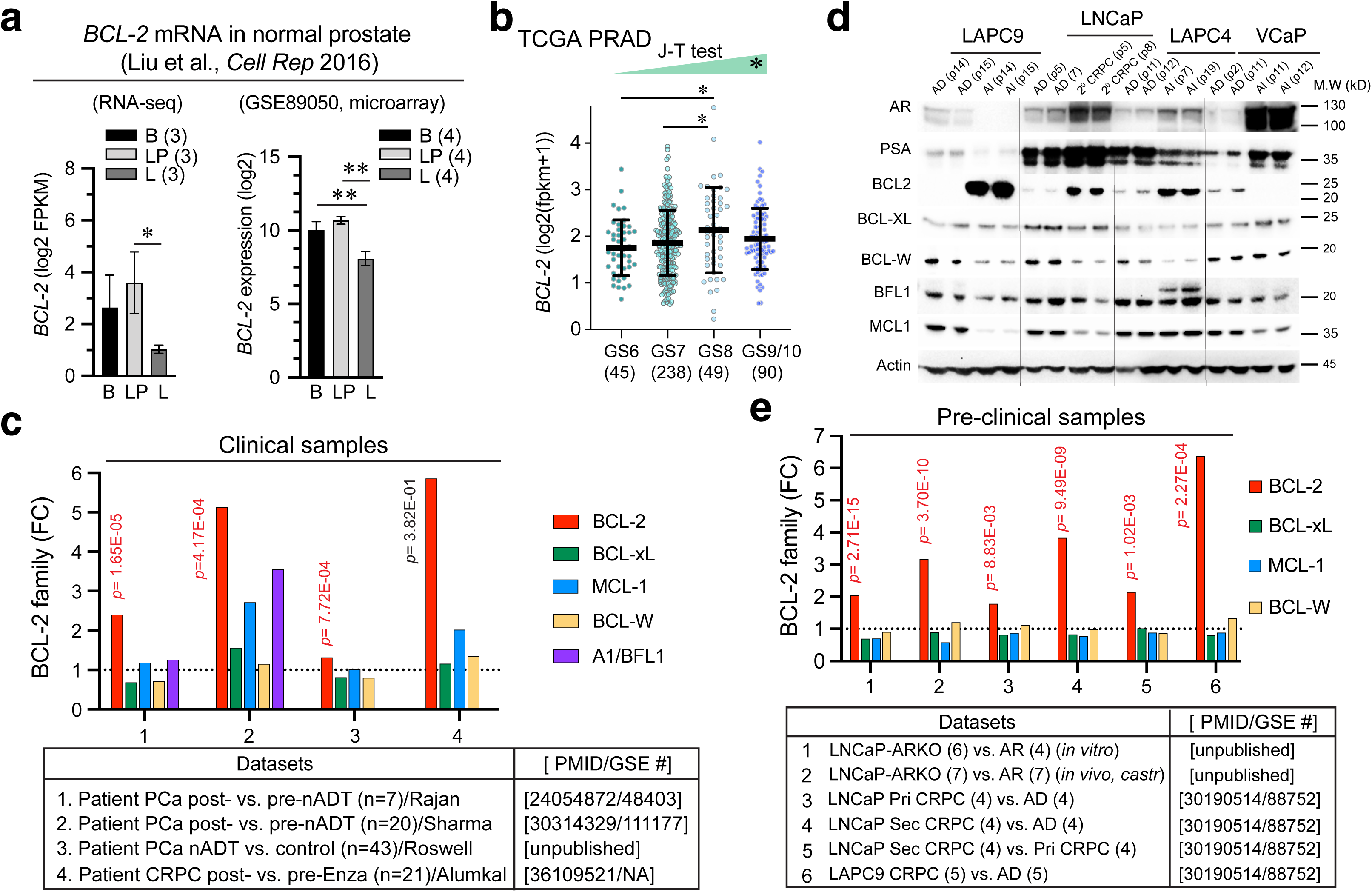
Selective upregulation of BCL-2 by castration (ADT) and antiandrogens. **a.** *BCL-2* mRNA is expressed at higher levels in normal human prostate basal (B) and CD38^lo^ luminal progenitor (LP) cells than mature differentiated luminal (L) cells. The original publication (dataset) is indicated on top (ref. 71), and the two bar graphs were derived from RNA-seq (left) and microarray (right), respectively. n indicated in parentheses. \**p*<0.05; \*\**p*<0.01 (paired Student’s *t*-test). **b.** *BCL-2* mRNA levels are increased in high-grade PCa and correlate with tumor grade. Data were extracted from TCGA_PRAD consisting of PCa with increasing grade (GS, Gleason Score; n indicated below). \**p*<0.05 (Student’s *t*-test). J-T trend test showed that *BCL-2* mRNA levels correlated with increasing tumor grade (\**p*<0.05). **c.** *BCL-2* mRNA (among 5 members) was upregulated in patients’ PCa treated with nADT (the first 3 datasets) and showed a trend of upregulation in the Alumkal pre-/post-Enza cohort (i.e., dataset 4). Note that *A1/BFL-1* was not detected in datasets 3 and 4. **d-e**. BCL-2 family protein (d) and mRNA (e) levels in AD/AI (CRPC) xenograft models. d, AD and AI tumors serially passaged (p; indicated in parentheses) were used in WB analysis of the proteins indicated. 20 μg protein was loaded for each lane and actin was used as loading control. e, *BCL-2* mRNA is commonly increased in 6 CRPC datasets (n indicated in parentheses). Datasets 1 and 2 were comparisons of LNCaP clones with intact AR vs. AR knockout (ARKO) either cultured in vitro (dataset 1) or propagated in castrated NSG mice (dataset 2). Datasets 3-6 were from our earlier publication (PMID and GSE# indicated).

In toto, these data from both PCa models and patient specimens indicate that *castration (ADT and/or ADT/Enza) commonly and preferentially upregulates BCL-2*.

### Treatment-naive primary prostate tumors are populated mostly by AR^+^BCL-2^-^ PCa cells

The above observations (Fig. 1) suggest a reciprocal relationship between AR signaling and BCL-2 expression. To confirm and extend these findings, we employed Vectra-based quantitative multiplex immunofluorescence (qmIF) to assess PCa cells expressing BCL-2 and/or AR in regular FFPE (formalin-fixed and paraffin-embedded) sections as well as TMA (tissue microarray) and whole-mount (WM) sections from benign prostate (n=123), treatment-naïve primary PCa (n=125) and treatment-failed CRPC (n=25) (Fig. 2; Supplementary Fig. S1–S5). We utilized cytokeratin (CK) staining to demarcate the epithelial compartment. Consistent with earlier reports^27,38,41,44^ and mRNA data (Fig. 1a), BCL-2 protein was expressed in the basal cell layer while AR was detected in luminal layer in benign glands (Fig. 2a; Supplementary Fig. S1c, Fig. S2a), and BCL-2^+^ cells and AR^+^ cells displayed a strong inverse correlation in benign prostate (R=-0.74, *p*=0.00159; Fig. 2e, top). In untreated primary PCa, there was a striking expansion of AR^+^ and AR^+^BCL-2^-^ cell populations compared to benign tissues (Fig. 2b; Supplementary Fig. S1d, Fig. S2b, Fig. S5c) and the AR^+^ and BCL-2^+^ PCa cells still exhibited an inverse correlation (R= −0.626, *p*=0.0014; Fig. 2e, middle). Consistent with decreases in BCL-2^+^ PCa cells, the *BCL-2* mRNA levels were reduced in primary tumors compared to benign tissues (Supplementary Fig. S5a-b).

**Figure 2.**
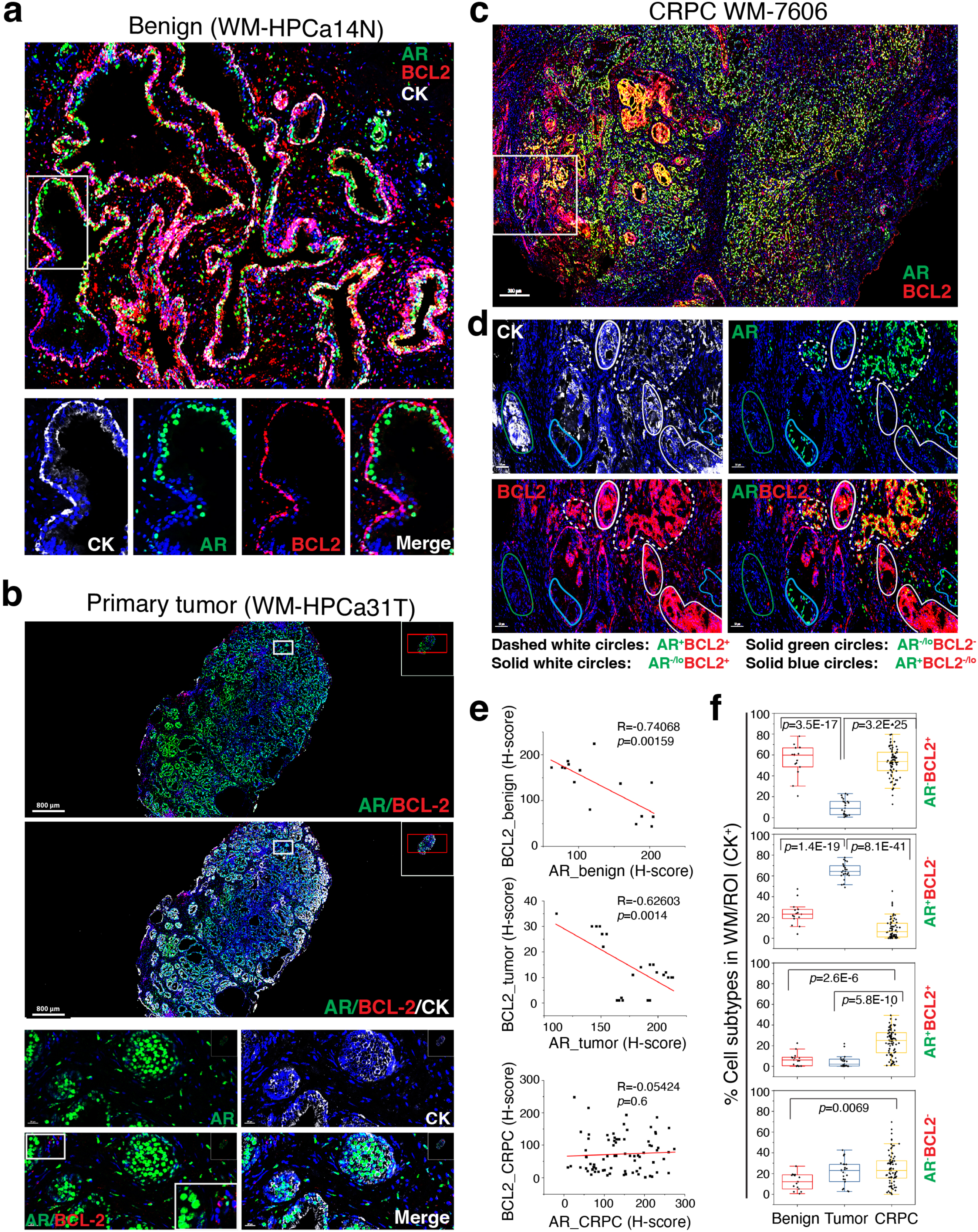
qmIF analysis reveals predominant AR^+^BCL-2^-^ cells in untreated primary PCa and markedly increased (AR^+^ or AR^-^) BCL-2^+^ cells in CRPC. **a.** In benign prostatic glands (HPCa14N), BCL-2^+^ cells are mainly in the basal cell layer whereas AR^+^ cells in luminal layer. Cytokeratin (CK) staining was used to mark epithelial compartment. Note both AR and BCL-2 were also expressed in stromal cells. Magnified images of individual stains from the boxed region in the whole-mount (WM) image (top) were presented below. **b.** Primary PCa is characterized by significantly increased AR^+^BCL-2^-^ cells. Shown above are two WM images of HPCa31T and down below zoom-in images of individual or merged markers. **c-d**. Increased cellular heterogeneity and markedly expanded (AR^+^ or AR^-^) BCL-2^+^ cell population in CRPC. c, WM low-magnification image showing AR/BCL-2 expression. d, Zoom-in images of the boxed area in c showing individual markers. **e.** Relationship between (CK^+^) AR-expressing and/or BCL-2-expressing cells in benign tissue (top), primary tumor (middle) and CRPC (bottom). Regression line, and Pearson R and *P* values are indicated. **f.** Box plots summarizing the relative % of 4 PCa cell subtypes in WM images analyzed in benign tissues, primary tumors and CRPC. Each dot in the box plots represents a CK^+^ ROI. *P* values were determined by repeated measures two-way ANOVA with Bonferroni multiple comparison test.

Cell subtype analysis of AR^+/-^ and/or BCL-2^+/-^ cells revealed that among all the CK^+^ ROIs (Region of Interest, 1 mm^2^) analyzed by Vectra in benign prostate, the AR^-^BCL-2^+^ cell population represented the majority, followed by AR^+^BCL-2^-^ cells, with very low representations of AR^+^BCL-2^+^ (double positive) and AR^-^BCL-2^-^ (double-negative) cells (Fig. 2f; Supplementary Fig. S5d-f). In primary PCa, the AR^+^BCL-2^-^ PCa cells became the predominant cell population followed by AR^-^BCL-2^-^ cells with low representations of the double-positive and AR^-^BCL-2^+^ cell subtypes (Fig. 2f; Supplementary Fig. S5d-f).

### Castration drives PCa cell heterogeneity and upregulates BCL-2^+^ PCa cells in patient CRPC

We analyzed a total of 25 CRPC including 20 in TMA and 5 WM sections (Supplementary Fig. S1b) and found that the reciprocal relationship between BCL-2^+^ and AR^+^ PCa cells observed in primary PCa was lost in CRPC (Fig. 2e). Relative to primary tumors, CRPC showed reduced fraction of AR^+^ but significantly increased fraction of BCL-2^+^ PCa cells (Supplementary Fig. S5c). Notably, we observed increased PCa cell heterogeneity: AR^-^BCL-2^+^ cells were nearly 50% of the 4 cell-type subpopulations (AR^-^ BCL-2^+^, AR^+^BCL-2^-^, AR^-^BCL-2^-^ and AR^+^BCL-2^+^ cells) amongst (CK^+^) CRPC cells (Fig. 2d, f; Supplementary Fig. S2c-d, Fig. S3c, Fig. S4 and Fig. S5d-f). We also observed significant increases in double-positive (AR^+^BCL-2^+^) PCa cells in CRPC (Fig. 2d, f; Supplementary Fig. S3b and Fig. S5d-f). Aggregating cell density (cell count/mm^2^) in WM sections (Supplementary Fig. S5e) or in all specimens (Supplementary Fig. S5f) and compared to primary PCa, we observed that the AR^+^BCL-2^-^ PCa cells represented the majority in primary tumors whereas in CRPC the AR^-^BCL2^+^ cells became the predominant cell population accompanied by increased AR^+^BCL-2^+^ and AR^-^BCL-2^-^ cells (Fig. 2f; Supplementary Fig. S5d). Of interest, in CRPC, while the majority of AR^+^ PCa cells showed nuclear AR (e.g., Fig. 2d; Supplementary Fig. S2d and Fig. S3b, c), we also observed increased AR^+^ CRPC cells with prominent cytoplasmic AR (AR^cyto^; Supplementary Fig. S4b, d), as we previously noted in patient CRPC^20^.

### Castration-induced dynamic changes in AR^+/-^BCL-2^+/-^ PCa cell subpopulations in xenograft tumors recapitulate the changes observed in patient CRPC

Next, we employed both Vectra-based qmIF and Imaging Mass Cytometry (IMC) platforms to quantitatively assess the dynamics of AR^+/-^BCL-2^+/-^ cells in the 4 well-annotated PCa xenograft models developed in our lab (Fig. 3; Supplementary Fig. S6 – S9), which were matched AD and AI (or primary (1°) CRPC) pairs of LNCaP, LAPC9, LAPC4 and VCaP tumors serially propagated in immunodeficient mice (Supplementary Fig. S6a).^14,17,18,20,24–26^ In accord with our previous findings,^20^ IHC characterization of the AR expression patterns and levels in 1° castration-resistant (AI) tumors revealed the LNCaP-AI tumors to be (nuclear) AR^+/hi^ and LAPC9-AI tumors AR^-/lo^ whereas LAPC4-AI and VCaP-AI tumors largely AR^cyto^ (Supplementary Fig. S6b). Re-analysis of individual AI tumors under therapy^20^ revealed distinct Enza responses associated with the AR heterogeneity in these CRPC models: the AR^+/hi^ LNCaP-AI responded to Enza for ∼4 weeks (Supplementary Fig. S6c) and AR^-/lo^ LAPC9-AI were *de novo* refractory to Enza (Supplementary Fig. S6d) whereas AR^cyto^ LAPC4-AI and VCaP-AI responded to Enza with distinct latencies prior to the emergence of castration/Enza-resistant 2° CRPC (Supplementary Fig. S6e-f).

**Figure 3.**
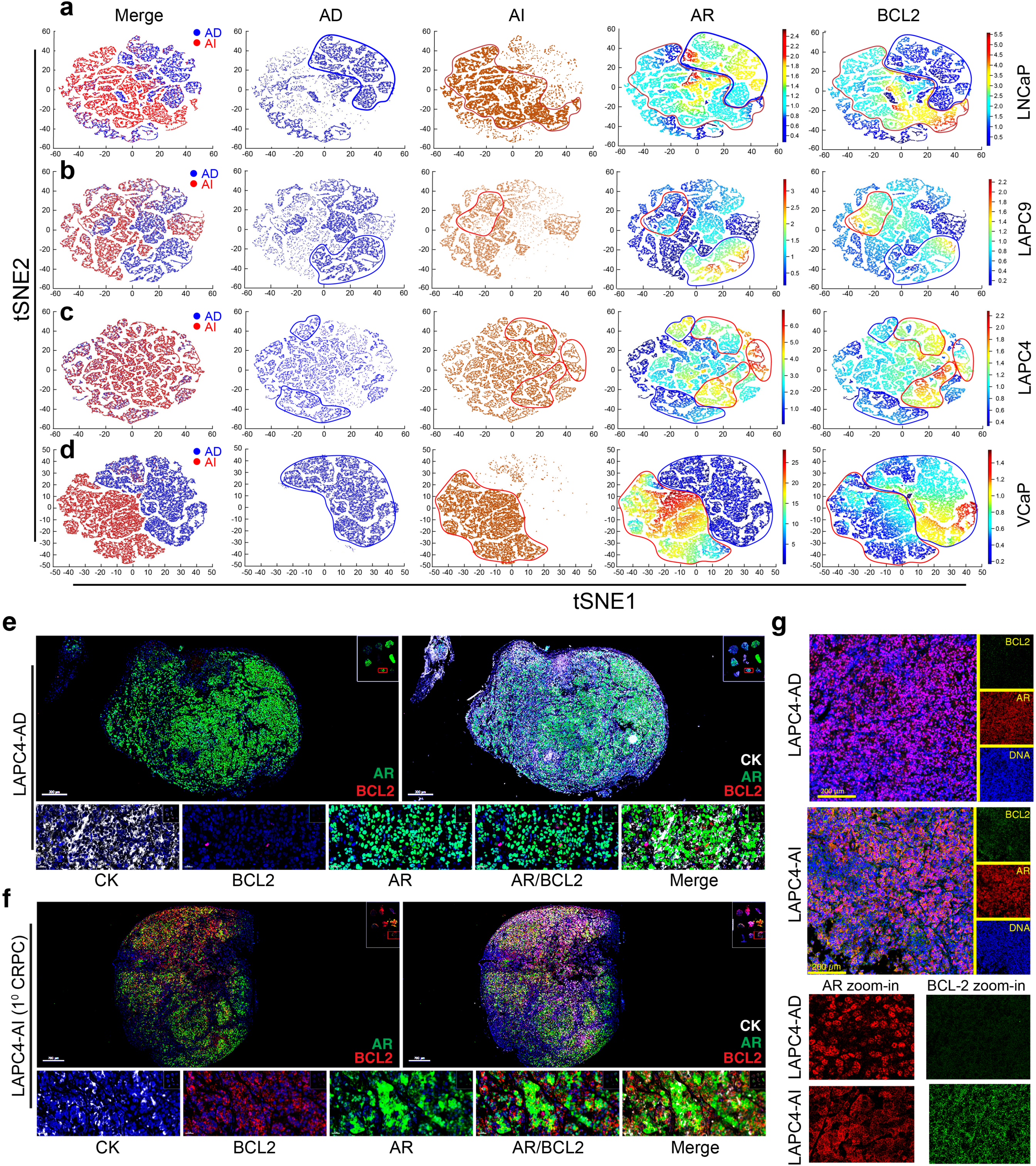
Vectra and IMC analysis reveals castration-induced dynamic changes in AR^+/-^BCL-2^+/-^ cell types in 4 xenograft CRPC models. **a-d**. tSNE plots of IMC-derived single-cell imaging data. Shown are the individual and merged tSNE plots from LNCaP-AD/AI (a), LAPC9-AD/AI (b), LAPC4-AD/AI (c), and VCaP-AD/AI (d) xenografts. Columns represent merged AD and AI cell clusters (first column), individual AD and AI cell populations and contours (second and third columns), and AR and BCL-2 expression overlaid on tSNE maps (fourth and fifth columns). Blue and red contours indicate AD and AI cell subpopulations, respectively. e. LAPC4-AD tumors are populated mostly by AR^+^BCL-2^-^ PCa cells. Shown on top are qmIF WM images and at the bottom zoom-in images. f. LAPC4-AI (1° CRPC) tumors are populated by AR^cyto^BCL-2^+^ PCa cells. Shown on top are qmIF WM images and at the bottom zoom-in images illustrating cytoplasmic AR^+^ LAPC4-AI cells with upregulated BCL-2. Representative zoom-in images of AR and BCL-2 are presented below.

In the LNCaP-AD/AI system (Fig. 3a; Supplementary Fig. S7b-e), qmIF analysis of WM LNCaP-AD tumors revealed that both AR^-^ and AR^+^ cells expressed little BCL-2 (Supplementary Fig. S7b). Most cells in LNCaP-AI (1° CRPC) tumors turned AR^+/hi^ with slightly increased BCL-2 (Supplementary Fig. S7c). Sensitive IMC-generated single-cell profiling with tSNE-based cell clustering corroborated AR and BCL-2 upregulation in LNCaP-AI cells (Fig. 3a; Supplementary Fig. S7e). Strikingly, most PCa cells in castration/Enza-resistant 2° LNCaP-CRPC were AR^+/hi^BCL-2^+/hi^ (Supplementary Fig. S7d). The LAPC9-AD tumors resembled LNCaP-AD tumors in containing both AR^-^BCL-2^-^ and AR^+^BCL-2^-^ cells (Supplementary Fig. S8a). In contrast, LAPC9-AI tumors, unlike LNCaP-AI tumors, apparently evolved into AR^-^BCL-2^+/hi^ (Supplementary Fig. S8b). IMC and tSNE-anchored cell subpopulation analyses also revealed reduced AR expression concomitant with elevated BCL-2, resulting in substantially increased AR^-^ BCL-2^+^ cells in LAPC9-AI tumors (Fig. 3b; Supplementary Fig. S8c). Most LAPC4-AD tumor cells exhibited an AR^+^BCL-2^-^ phenotype on qmIF analysis (Fig. 3e) and castration reprogrammed LAPC4-AI cells to the AR^cyto^BCL-2^+^ phenotype (Fig. 3f). Single-cell analysis by IMC confirmed the qmIF-described AR^cyto^BCL-2^+^ phenotype in LAPC4-AI tumors (Fig. 3c, g). Finally, the VCaP AD/AI model was an exception, where most VCaP-AD tumor cells were AR^+^BCL-2^+^ while VCaP-AI tumor cells became AR^cyto^BCL-2^-/lo^ (Supplementary Fig. S9a-b). IMC analysis validated the AR^+^BCL-2^+^ phenotype of VCaP-AD cells and demonstrated cytoplasmic AR redistribution and reduced BCL-2 in VCaP-AI tumors (Fig. 3d; Supplementary Fig. S9c).

In summary, 3 of the 4 AD xenograft models (LNCaP, LAPC9 and LAPC4) recapitulated the major AR^+^BCL-2^-^ cellular phenotype observed in human primary PCa whereas VCaP-AD tumors, intriguingly, displayed the AR^+^BCL-2^+^ phenotype. Consistent with the diverse AR^+/-^BCL-2^+/-^ cell subtypes observed in patient CRPC, the 4 xenograft CRPC models manifested variegated cellular subpopulations: the LNCaP (1° and 2°) CRPC predominantly AR^+^BCL-2^+^, LAPC9-CRPC predominantly AR^-^BCL-2^+^, LAPC4-CRPC predominantly AR^cyto^BCL-2^+^ and VCaP-CRPC predominantly AR^cyto^BCL-2^-^ phenotype, respectively.

### Developing castration/Enza-resistant LAPC4 culture models to recapitulate castration-induced AR^cyto^BCL-2^+^ phenotype

Our previous analysis of 195 CRPC specimens revealed that as much as 40% of the CRPC had the mixed cytoplasmic/nuclear AR^20^. qmIF studies herein also identified prominent AR^cyto^ CRPC cells, which often expressed high levels of BCL-2 and presented an AR^cyto^BCL-2^+^ phenotype (e.g., Supplementary Fig. S4b, d), much like the AR^cyto^BCL-2^+^ LAPC4-CRPC *in vivo* (Fig. 3f, g). To further study the unique phenotype of AR^cyto^BCL-2^+^ CRPC, we established the castration-resistant and dual castration- and Enza-resistant LAPC4 cell models (Fig. 4; Supplementary Fig. S10). To this end, we cultured regular LAPC4-AD cells long-term (4 months) in charcoal dextran stripped serum (CDSS) medium to develop castration-resistant LAPC4 (LAPC4-CR or LAPC4-AI) cells, which were subsequently exposed to either 20 μM or 100 μM Enza-containing CDSS medium for 1 month resulting in the LAPC4-Enza(20)-R or LAPC4-Enza(100)-R models (Supplementary Fig. S10a). These CDSS and CDSS/Enza selected LAPC4 sublines displayed more elongated and mesenchymal morphology (Supplementary Fig. S10b) and proliferated more slowly (Supplementary Fig. S10c) than LAPC4-AD cells. Moreover, although LAPC4-AD cells were inhibited by both 20 μM and 100 μM of Enza (Fig. 4a), the LAPC4-CR and LAPC4-Enza(20)-R cells were sensitive only to 100 μM of Enza (Fig. 4b-c). LAPC4-Enza(100)-R cells were resistant to both 20 μM and 100 μM of Enza (Fig. 4d). Western blotting (WB) analysis revealed a progressive re-distribution of AR from nucleus to cytoplasm and increase in BCL-2 expression (Fig. 4e–f). Multiplex IF imaging analysis also showed a shift from nuclear AR^+^ and low BCL-2 in LAPC4-AD to an AR^cyto^BCL-2^+/hi^ phenotype in resistant sublines, especially in LAPC4-Enza(100)-R cells in which BCL-2 co-localized with the Mito-tracker (Fig. 4g; Supplementary Fig. S10d). The LAPC4-Enza(20)-R cells displayed an intermediate phenotype (Supplementary Fig. S10d), suggesting an ongoing transition. Quantification of cell subtypes demonstrated that in LAPC4-AD cells, the AR^+^BCL-2^-^ phenotype predominated while the resistant sublines showed expansion in AR^cyto^(AR^-^)BCL-2^+^ and AR^+^BCL-2^+^ subpopulations (Fig. 4h).

**Figure 4.**
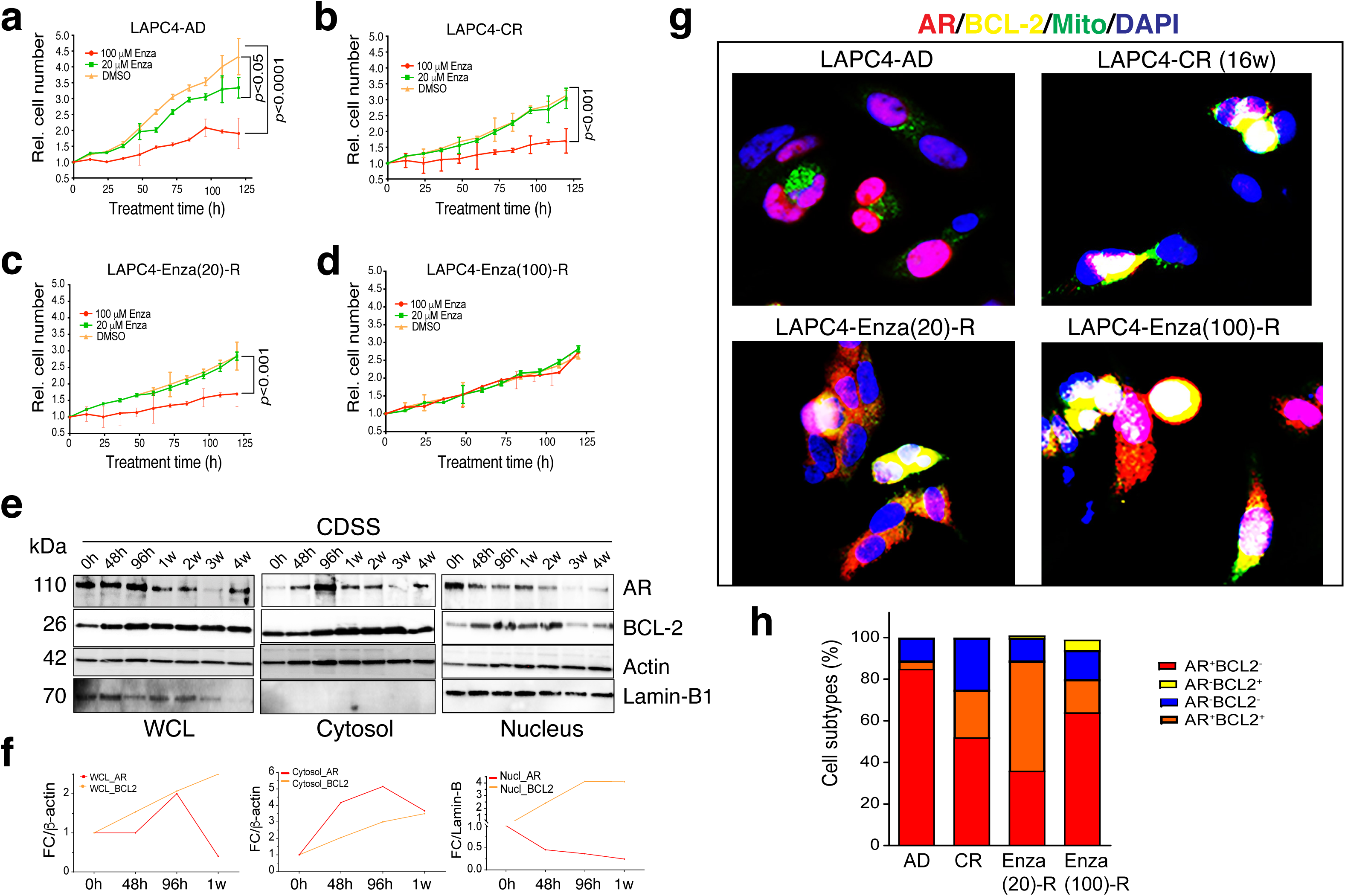
Enzalutamide resistance is associated with AR cytoplasmic relocalization and BCL-2 upregulation in LAPC4 models. **a–d.** Cell proliferation assays showing relative cell numbers over time in LAPC4-AD (a), LAPC4-CR (b), and two Enza-resistant sublines—LAPC4-Enza(20)-R (c) and LAPC4-Enza(100)-R (d). Shown are the mean ± SEM (*p<* 0.001, two-way ANOVA). **e.** Immunoblot analysis of LAPC4-AD cells cultured in CDSS over time shows dynamic changes in AR and BCL-2 expression using whole-cell lysates (WCL), and cytosolic and nuclear fractions. Lamin-B1 and actin serve as loading controls. **f.** Densitometric quantification of immunoblots in (e) showing fold change (FC) of AR and BCL-2 relative to loading controls over the time course in WCL (left) and cytosolic (middle) and nuclear fractions (right). **g.** IF staining of LAPC4-AD, LAPC4-CR, and Enza-resistant sublines (20 μM and 100 μM) showing AR (red), BCL-2 (yellow), mitochondria (green), and nuclei (DAPI, blue). Images reveal increased cytoplasmic AR and mitochondrial BCL-2 in resistant sublines. **h.** Quantification of AR and BCL-2 expression patterns from the immunofluorescence data in (g), categorized into four groups: AR⁺BCL-2⁻, AR⁺BCL-2⁺ (or AR^cyto^BCL-2⁺), AR⁻BCL-2⁺, and AR⁻BCL-2⁻.

### AR directly binds to the BCL-2 genomic locus to repress BCL-2 transcription

These qmIF and IMC studies of BCL-2 and AR protein expression at the single-cell levels in patient CRPC and xenograft models (Figs. 1-4) revealed a reciprocal relationship between the two, suggesting that AR may transcriptionally represses *BCL-2*. In support of this relationship, in 3 cohorts of PCa patients who underwent nADT,^72–74^ *BCL-2* mRNA levels exhibited a perfect inverse correlation with the AR activity^75^: AR activity was high and *BCL-2* was low in pre-nADT PCa whereas AR activity was reduced but *BCL-2* mRNA levels were upregulated in patient-matched post-nADT samples (Fig. 5a). We also observed an inverse correlation between *BCL-2* mRNA levels and AR activity in an in-house RNA-seq dataset of 25 PCa patients treated for greater than 2 months with nADT compared with 25 stage/grade matched control cases (i.e., no nADT) (Fig. 5b). Finally, in a dataset of 40 patient and organoid CRPC samples, which were classified by the ATAC-seq (Assay for Transposase-Accessible Chromatin using sequencing) and RNA-seq analyses as CRPC_AR (*AR* high), CRPC_SCL (stem-cell like; *AR* low), CRPC_Wnt (*AR* negative), and CRPC_NE (*AR*-negative) subtypes^76^, we similarly observed inversely correlated *BCL-2* and *AR* mRNA levels - CRPC_AR subtype had the highest *AR* and lowest *BCL-2* while CRPC_Wnt and CRPC_NE had the lowest *AR* but highest BCL-2 (Fig. 5c). Interestingly, among the 13 CRPC_SCL samples, 7 co-segregated with the CRPC_AR subtype and 6 with the CRPC_Wnt/CRPC_NE (Fig. 5c), suggesting that this chromatin accessibility-stratified CRPC_SCL subtype is heterogeneous with ‘bifurcated’ *AR*^hi^*BCL*-2^lo^ and *AR*^lo^*BCL-2*^hi^ profiles.

**Figure 5:**
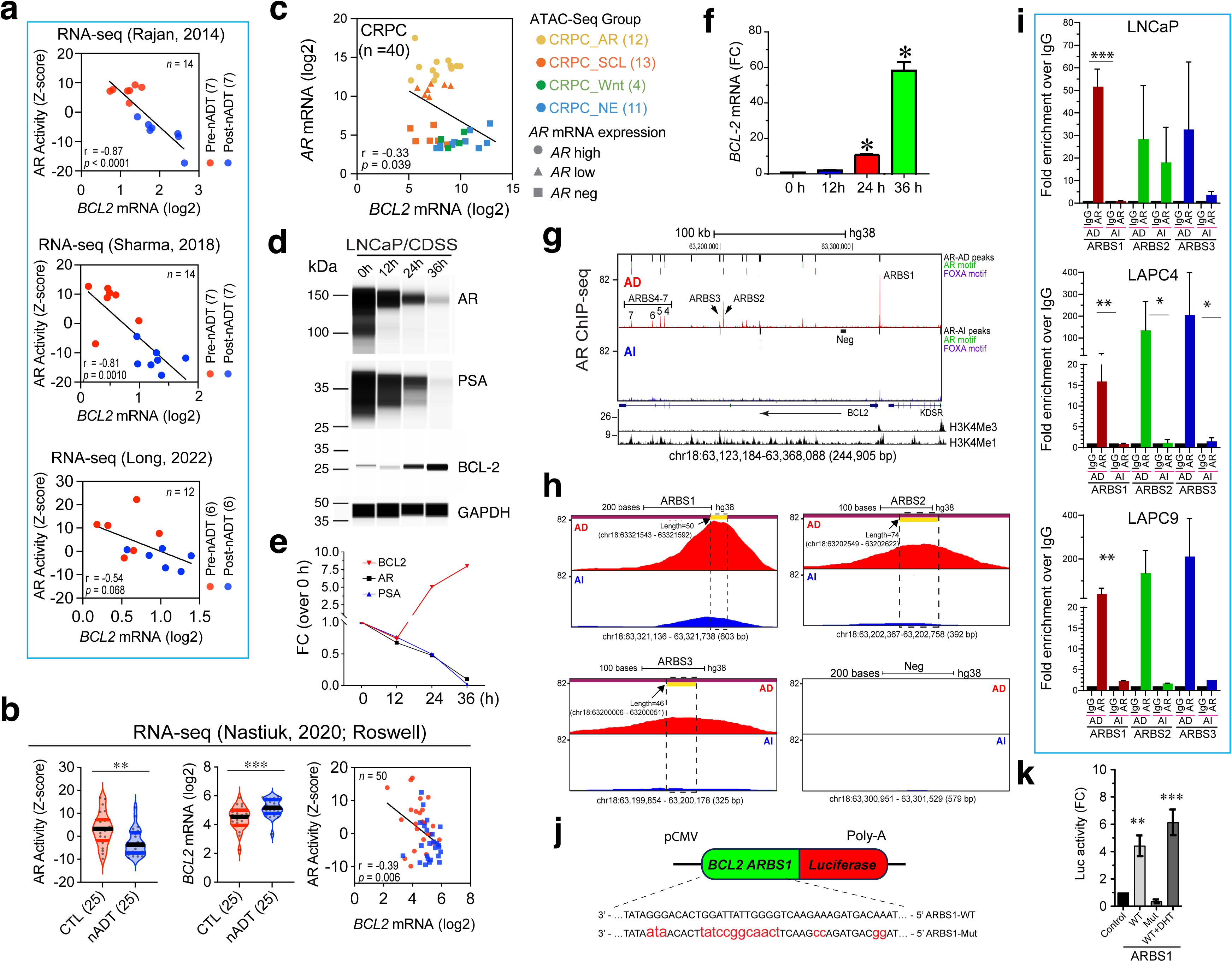
AR directly represses *BCL-2* transcription in PCa. **a.** Inverse correlation between BCL-2 mRNA levels and AR activity in 3 nADT cohorts. **b.** Upregulation of *BCL-2* mRNA and inverse correlation between *BCL-2* mRNA levels and AR activity in the matched Roswell Park nADT cohort. **_c._** Inverse correlation between *BCL-2* and *AR* mRNA levels in the 40 CRPC specimens in the Tang F et al dataset.^76^ **d-f**. LNCaP cells castrated in vitro time-dependently downregulate AR and AR signaling but upregulate BCL-2. Shown are WB of AR, PSA and BCL-2 protein levels in LNCaP cells cultured in CDSS-containing medium for the time intervals indicated (d), densitometric quantification normalized to GAPDH (e), and changes in *BCL-2* mRNA levels measured by RT-qPCR (f; n=3, mean ± SD; \**p*<0.05). g. AR binds to multiple ARBS in *BCL-2* genomic locus. Shown are the 7 potential ARBS in LNCaP-AD (top) that are lost in LNCaP-AI (bottom) cells (Neg: genomic region used as negative control in ChIP-qPCR). Shown on right are AR peaks and canonical AR and FOXA motifs and below H3K4Me3 and H3K4Me1 peaks. h. Zoom-in representation of ARBS1-3 in LNCaP-AD (top) vs. LNCaP-AI (bottom) cells. i. ChIP–qPCR analysis of AR binding at the ARBS1–3 in the indicated 3 AD/AI xenograft models. The bar graphs represent the mean ± S.D of 3 independent experiments (\**p*<0.05; \*\**p*<0.01; \*\*\**p*<0.001; Student’s *t*-test). j. Schematic of the luciferase reporter construct containing wild-type (WT) or mutated (Mut) ARBS1 from the *BCL2* locus. k. Luciferase (Luc) assays in LNCaP cells showing strong induction of Luc activity by ARBS1-WT but not the ARBS1-Mut (n=2; \*\**p*<0.01; \*\*\**p*<0.001; Student’s *t*-test).

We observed a similar inverse relationship between *BCL-2* mRNA levels and AR expression/activity in LNCaP-AD/AI xenografts^20^ (GSE88752) (Supplementary Fig. S11a). Experimental castration of LNCaP cells time-dependently downregulated AR protein and inhibited AR activity over time (Fig. 5d-e) but induced BCL-2 mRNA and protein (Fig. 5e, f). We analyzed the AR chromatin immunoprecipitation sequencing (ChIP-seq) data from LNCaP cells under regular (AD; GSM699631) and castrated (AI; GSM699630) conditions (Fig. 5g). We observed, in LNCaP-AD cells, multiple AR-binding sites (ARBS) across the *BCL-2* locus (Chr18: 63.12–63.32 Mb), some of which were enriched for AR and/or FOXA1 motifs (Fig. 5g, top). We identified seven prominent ARBS peaks (ARBS1–7), with the ARBS1 representing the main peak located at the *BCL-2* promoter region overlapping with histone mark H3K4Me3 associated with active promoters (Fig. 5g). ARBS1 and most other ARBSs were also enriched in H3K4Me1, a histone mark linked to enhancers and active gene transcriptions (Fig. 5g). Also of interest, ARBS1 (and ARBS3 and ARBS6) showed no AR or FOXA1 motif and ARBS7 harbored both motifs while the remaining peaks (ARBS 2, 4 and 5) were associated with either motif (Fig. 5g; top). Remarkably, in LNCaP-AI cells, AR binding was lost at all sites except for a faint residual peak at ARBS1 (Fig. 5g; bottom). We performed ChIP-qPCR experiments using primers specific for ARBS1 – ARBS3 (Fig. 5h) in our paired LNCaP-AD/AI, LAPC4-AD/AI and LAPC9-AD/AI xenograft models^20^ in which all 3 AI models upregulated BCL-2 (Fig. 1d). We found that the AR binding to ARBS1 in the *BCL-2* locus was lost in the 3 AI models whereas AR binding to ARBS2 and ARBS3 was either lost or greatly reduced (Fig. 5i). These results suggest that the ARBS1 represents the major *cis*-regulatory region of BCL-2 by AR. We cloned wildtype (WT) and a mutated (Mut) version of *BCL-2* ARBS1 into a luciferase construct (Fig. 5j) and luciferase reporter assays in LNCaP cells demonstrated that, although the WT ARBS1 drove robust luciferase activity that was further enhanced by DHT, the Mut ARBS1 completely lost luciferase activity (Fig. 5k).

Together, these data suggest that AR directly binds and represses the *BCL-2* locus in a ligand-dependent manner and inhibition of AR signaling would block *AR* occupancy at key regulatory regions such as ARBS1 allowing for transcriptional activation of *BCL-2*. In support, most ARBS, especially the ARBS1, were reduced or lost in multiple castration-resistant LNCaP sublines (Supplementary Fig. S11b). We interrogated an integrated dataset (GSE130408) of AR ChIP-seq in patient primary PCa and CRPC^77^ and observed globally reduced AR enrichment across *BCL-2* genomic region in CRPC compared to primary PCa (Supplementary Fig. S11c-h). We identified a total of 15 AR-binding peaks (except for ARBS6; Supplementary Fig. S11c, d, f) in patient PCa and CRPC samples all of which (except ARBS11 and ARBS12) were reduced in CRPC (Supplementary Fig. S11g). Side-by-side comparisons in primary PCa and CRPC showed that all 15 ARBS occupancy was decreased in CRPC (Supplementary Fig. S11g-h).

### Increased chromatin accessibility in the BCL-2 genomic region in AR^-/lo^BCL-2^+^ CRPC

We leveraged the aforementioned (Fig. 5c) ATAC-/RNA-seq dataset of 40 patient and organoid CRPC samples^76^ to determine the potential differences in chromatin accessibility in the *BCL-2* locus in AR^+^ vs. AR^-/lo^ CRPC (Supplementary Fig. S12). We found that in CRPC_AR tumors, chromatin accessibility across the *BCL-2* genomic region remained low (Supplementary Fig. S12a), which coincided with low *BCL-2* transcript levels (Supplementary Fig. S12b). In contrast, the AR^-/lo^ CRPC_Wnt and CRPC_NE exhibited increased chromatin accessibility (Supplementary Fig. S12a), accompanied by elevated *BCL-2* mRNA expression (Supplementary Figure S12b). Quantitative analysis of ATAC-seq peak intensities across CRPC subtypes revealed a strong positive correlation between *BCL-2* mRNA levels and chromatin accessibility, with the AR^-/lo^ CRPC_NE and CRPC_Wnt exhibiting the highest values for both (Supplementary Fig. S12c). Furthermore, *BCL-2* mRNA levels were found to inversely correlate with AR activity (Supplementary Fig. S12d).

These findings reveal increased chromatin accessibility surrounding the *BCL-2* genomic locus in AR^-/lo^BCL-2^+^ subtype of CRPC and suggest that loss of AR expression and signaling might also promote BCL-2 expression via modulating the chromatin landscape.

### The AR^cyto^ subtype of CRPC is susceptible to BCL-2 inhibition

Hereafter, we extended our study to specifically address whether BCL-2 plays a functional role in CRPC progression and may represent a therapeutic vulnerability in diverse subtypes of CRPC (Fig. 6, 7; Supplementary Fig. S13, S14). We started with the LAPC4-AI model, representing the AR^cyto^ CRPC subtype, as shown by castration-induced AR^cyto^ phenotype *in vivo* (Fig. 3f, g; Supplementary Fig. S6b) and *in vitro* (Fig. 4; Supplementary Fig. S10) and whose clinical relevance is supported by presence of AR^cyto20^ and AR^cyto^BCL-2^+^ cells (Supplementary Fig. S4b, d) in patient CRPC. WB revealed that the LAPC4 1° CRPC and, particularly, the 2° CRPC significantly upregulated AR and GR (glucocorticoid receptor) as well as BCL-2 (Fig. 6a,b). We conducted drug sensitivity assays in organoids derived from LAPC4-AD and LAPC4-AI tumors (Fig. 6c-g; Supplementary Fig. S13), which manifested the AR^+^BCL-2^-^ and AR^cyto^BCL-2^+^ phenotypes, respectively (Fig. 3e-g). Under optimized assay conditions (Supplementary Fig. S13a-c), the LAPC4-AD organoids were more sensitive to Enza than LAPC4-AI organoids (Fig. 6c,d; IC_50_ 43 μM vs. 95 μM, respectively; p=0.0286, Mann-Whitney U test). In contrast, the BCL-2 inhibitor (BCL-2i) ABT-199 elicited selective toxicity to LAPC4-AI organoids (Fig. 6e-f; IC_50_ ∼10 μM) but barely showed any inhibitory effect on organoids of LAPC4-AD tumors (Fig. 6e-f; IC_50_ ∼10 μM) that lacked BCL-2 expression. Importantly, combination of Enza and ABT-199 synergistically inhibited the LAPC4-AI organoids (Fig. 6g; Supplementary Fig. S13d-f) but not LAPC4-AD organoids (Supplementary Fig. S13g). Interestingly, RU486, a GR antagonist, although showing overall similar toxicities against LAPC4-AD and LAPC4-AI organoids when used alone (Supplementary Fig. S13h, i), exhibited synergistic inhibitory effects on LAPC4-AI organoids when combined with Enza (Supplementary Fig. S13j).

**Figure 6:**
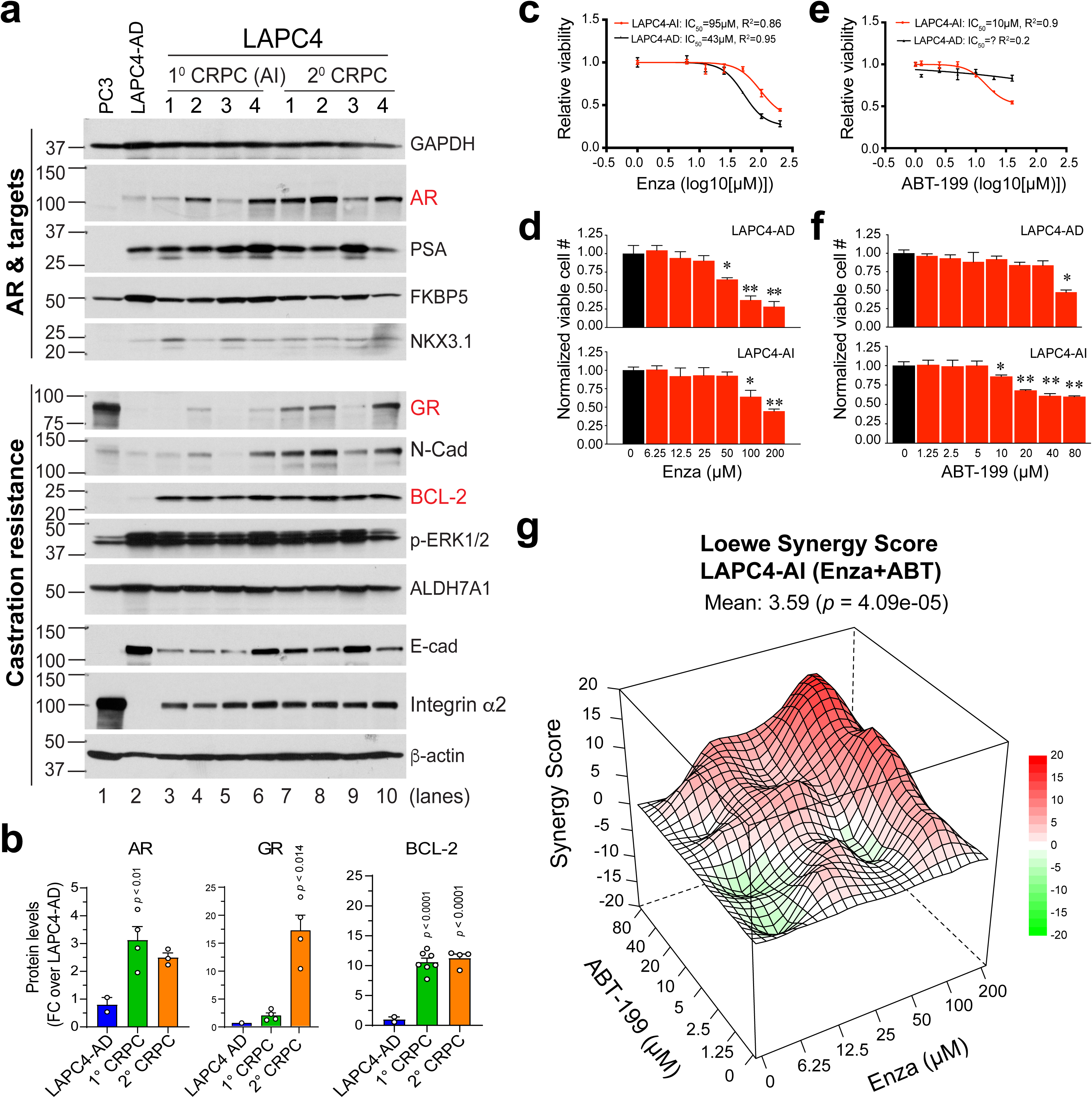
BCL-2 induced in the AR^cyto^ LAPC4-CRPC represents a therapeutic vulnerability. **a.** WB analysis of representative regulators in AR signaling and castration resistance in LAPC4-AD and its derived CRPC xenograft tumors, including first-generation (1° CRPC; lanes 3–6) and second-generation (2° CRPC; lanes 7–10) tumors. PC3 and parental LAPC4-AD cells (lanes 1–2) serve as controls. AR, GR and BCL-2 are highlighted in red. b. Quantification of AR, GR, and BCL-2 protein levels from panel a, normalized to β-actin and shown as fold-change relative to LAPC4-AD (mean ± SD, n = 4 tumors/group; *p*-values were calculated using a two-tailed test). **c–f.** Dose-response curves showing reduced sensitivity to Enza but increased response to ABT-199 in LAPC4-AI compared to LAPC4-AD in organoids assays, with corresponding IC₅₀ values indicated (panel c and e). Cell viability was measured by Resazurin. Data represent the mean ± SD (n=3; \**p<* 0.05, \*\**p<* 0.01; Student’s *t-*test). **g.** Loewe synergy analysis evaluating the combined effects of Enza and ABT-199 on LAPC4-AI cells. A 3D synergy surface plot depicts the interaction across increasing concentrations of both agents.

We then performed *in vivo* therapeutic studies in castrated male NOD/SCID mice bearing LAPC4-AI tumors (Fig. 7a; Supplementary Fig. S14a, b). The results revealed that although Enza modestly inhibited LAPC4-AI, ABT-199 exhibited strong tumor-inhibitory effects on AR^cyto^BCL-2^+^ LAPC4-AI tumors whereas the ABT-199/Enza combination demonstrated only slightly better tumor-controlling effects than ABT-199 alone (Fig. 7b; Supplementary Fig. S14c, d).

**Figure 7:**
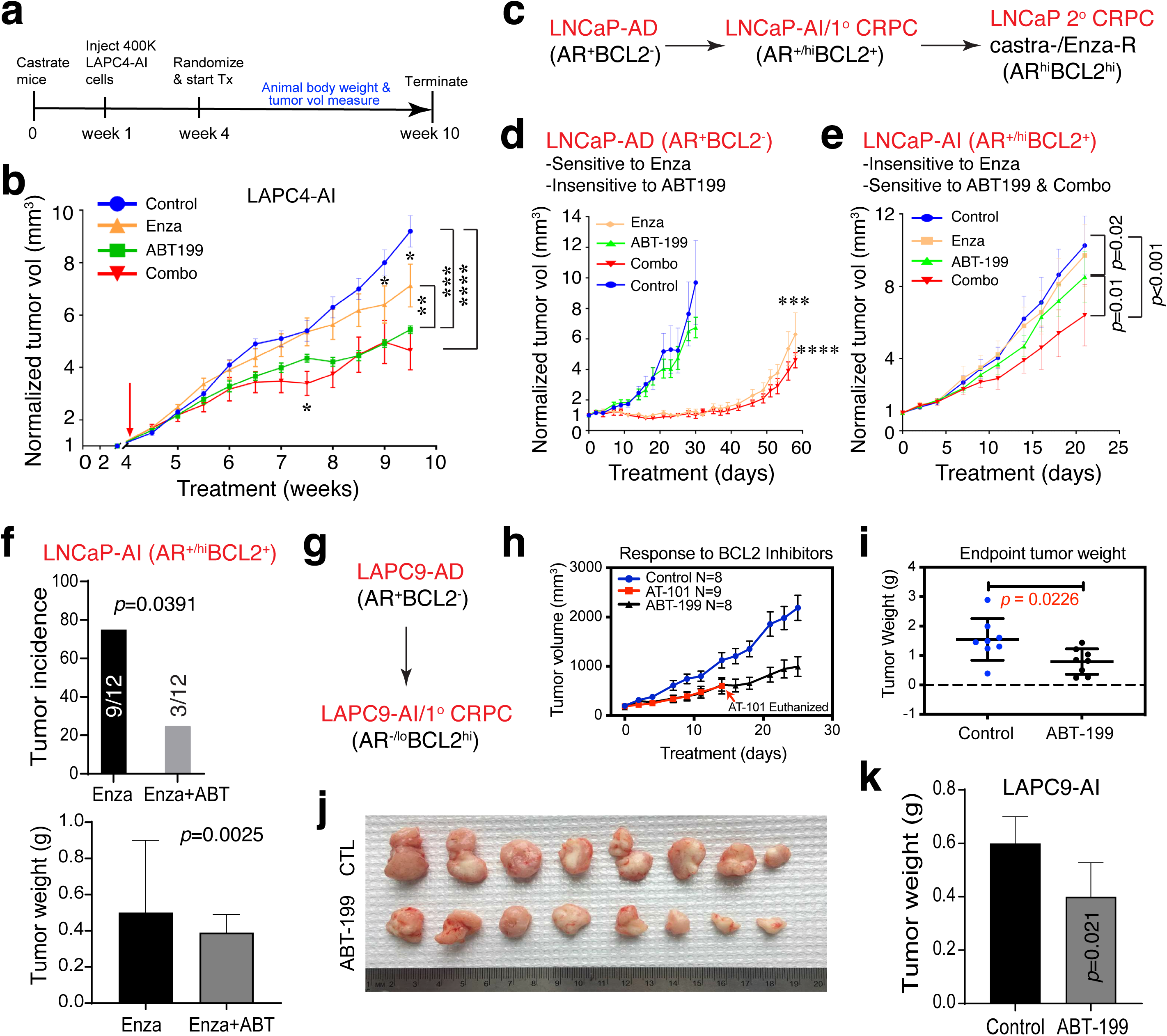
BCL-2 represents a functional therapeutic target across distinct CRPC subtypes. **a-b.** Experimental schema (a) and tumor growth curves of LAPC4-AI xenografts under indicated treatments (n=8 mice/group) (b). Presented are the tumor volumes normalized to the mean tumor volumes at the beginning of treatment, i.e., week 4 (mean ± SEM; \*\**p<* 0.001, \*\*\**p<* 0.0001, and \*\*\*\**p*<0.00001; two-way ANOVA). **c-f**. Therapeutic studies in the progressive LNCaP-AD/LNCaP-AI models (c). In vivo tumor growth of LNCaP-AD (d) and LNCaP-AI (e) xenografts treated with vehicle (control), Enza, ABT-199, or Enza + ABT-199 (Combo). The results showed that LNCaP-AD tumors were sensitive to Enza but not ABT-199 (d) while LNCaP-AI tumors are resistant to Enza but sensitive to ABT-199 and the combination (e). Tumor volume was normalized to baseline (n = 10 per group; mean ± SEM; *p*-values determined by two-way ANOVA). Shown in f is an independent therapeutic study showing significant inhibition of tumor incidence (top) and weight (bottom) in LNCaP-AI tumors treated with Enza and ABT-199 (ABT) combination compared to Enza monotherapy (tumor incidence and endpoint weight were determined by Fisher’s exact test and unpaired Student’s *t*-test, respectively). **g-k.** Therapeutic studies in the LAPC9-AI model. Shown are tumor growth curves of LAPC9-AI xenografts treated with ABT-199 (h; note the systemic toxicities of AT-101), endpoint tumor weight (i) and images (j), and an independent ABT-199 monotherapy study in LAPC9-AI xenografts (n=8 mice/group; p-value determined by Student’s *t-*test).

### The AR^+/hi^ subtype of CRPC is similarly sensitive to BCL-2 inhibition

Next, we examined the sensitivity of AR^+/hi^ subtype of CRPC to ABT-199 using the progressive LNCaP-AD ® 1° CRPC (AI) → 2° CRPC (castration/Enza-resistant) models (Fig. 7c; Supplementary Fig. S7). We first established castration- and castration/Enza-resistant LNCaP cell models that showed similar dynamic changes in AR and BCL-2 to the *in vivo* models (Supplementary Fig. S14e-g). Briefly, cells were cultured in CDSS-containing media for 8 weeks to generate castration-resistant LNCaP-CR cells, which were then cultured in CDSS medium containing Enza (50 µM) for 2 weeks to derive LNCaP-C/Enza-R cells (Supplementary Fig. S14e). Immunoblot analysis showed a reduction in AR protein levels in LNCaP-CR cells, followed by AR re-expression in LNCaP-C/Enza-R cells; in contrast, BCL-2 expression progressively increased from the LNCaP-CR to LNCaP-C/Enza-R states (Supplementary Fig. S14f, g). LNCaP-C/Enza-R cells were treated with increasing doses of ABT-199 (Supplementary Fig. S14h) and viability assays demonstrated significantly higher sensitivity in LNCaP-C/Enz-R cells to BCL-2 inhibition across all doses compared to LNCaP-AD cells (Supplementary Figure S14i).

*In vivo* therapeutic studies also revealed strikingly different responses based on androgen dependency: the AR^+^BCL-2^-^ LNCaP-AD tumors responded to Enza but not ABT-199 and the combination treatment resulted in similar tumor-inhibitory effects to Enza alone (Fig. 7d) while the AR^+/hi^BCL-2^+^ LNCaP-AI tumors responded well to ABT-199 and the combination treatment resulted in more pronounced tumor-inhibitory effects than ABT-199 alone (Fig. 7e). In an independent therapeutic study, the ABT-199/Enza combination more prominently inhibited both the incidence and endpoint weights of the LNCaP-AI tumors compared to Enza alone (Fig. 7f).

### The AR^-/lo^ subtype of CRPC is also vulnerable to BCL-2 inhibition

Finally, we asked whether the AR^-/lo^BCL-2^+^ LAPC9-AI (Fig. 7g; Supplementary Fig. S8), which was refractory to Enza^20^ (Supplementary Fig. S6d), may also be sensitive to BCL-2 inhibition. To this end, we treated castrated male NOD/SCID mice bearing LAPC9-AI tumors (Supplementary Fig. S14j, k) with ABT-199 or as a control, AT-101, an early-generation BCL-2/BCL-xL inhibitor derived from gossypol.^57,58^ We observed that ABT-199 significantly inhibited the growth of LAPC9-AI tumors (Fig. 7h-j) without apparent toxicity (Supplementary Fig. S14l). An independent therapeutic study verified the inhibitory effect of ABT-199 on LAPC9-AI (Fig. 7k). AT-101 also inhibited LAPC9-AI growth (Fig. 7h) but it manifested significant systemic toxicity in mice such that the treatment had to be terminated early (Supplementary Fig. S14l).

### Correlative studies in a phase Ib clinical trial link BCL-2 inhibition to treatment response

These preclinical studies revealed the efficacy of BCL-2i ABT-199, either alone or together with Enza, in inhibiting the 3 subtypes (i.e., AR^+/hi^, AR^-/lo^, and AR^cyto^) of BCL-2^+^ CRPC. These results, coupled with our early observations^20^, led us to conduct a Phase Ib clinical trial^70^ evaluating Enza plus venetoclax (ABT-199) in 10 patients with mCRPC (NCT03751436), and our correlative studies suggested potential therapeutic benefit of this combination in a subset of mCRPC patients (Fig. 8; Supplementary Fig. S15).

**Figure 8:**
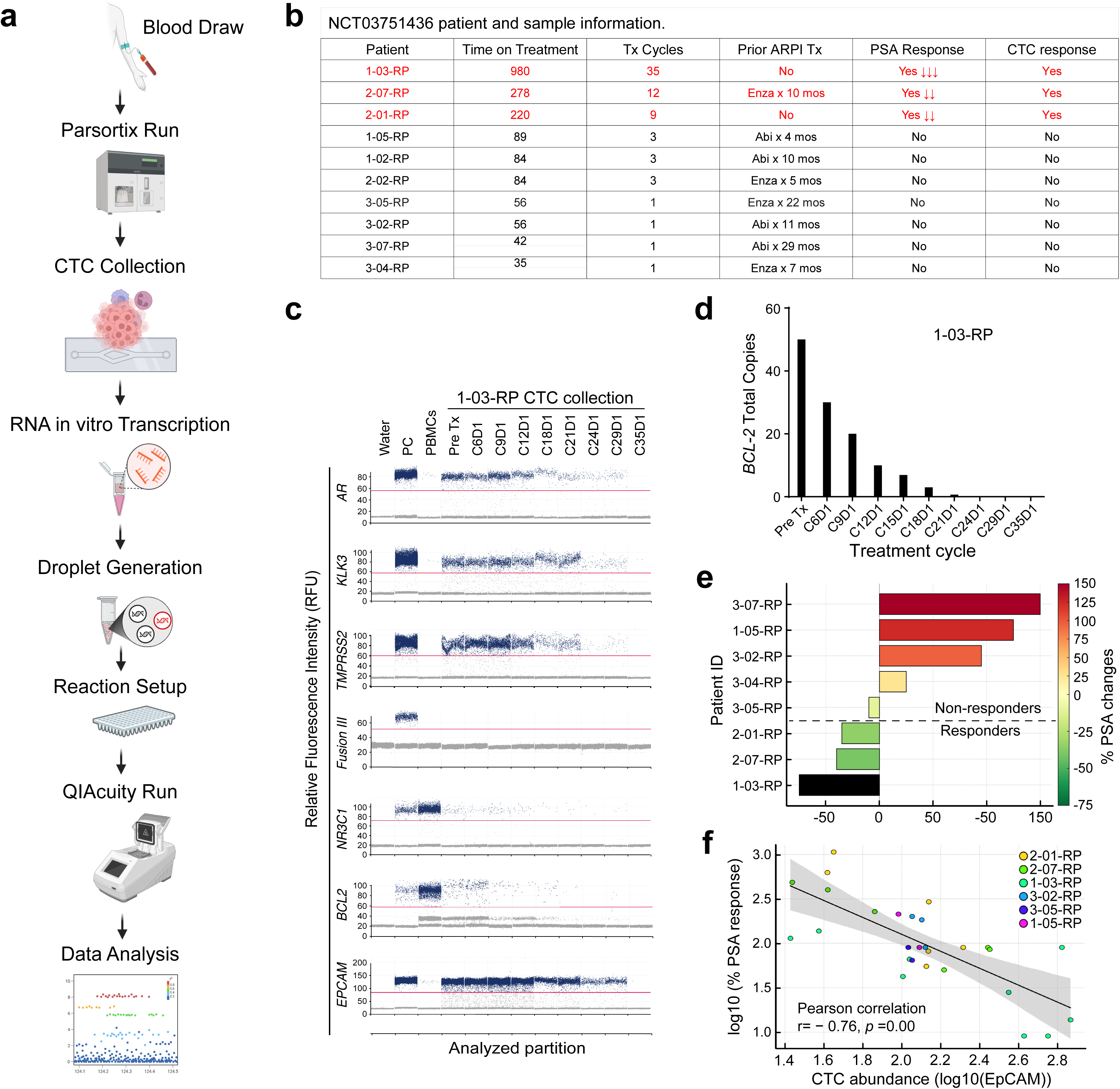
Correlative studies of a Phase Ib trial validates BCL-2 as a target in PCa patients. **a.** Schematic workflow. **b.** Tabulated summary of the trial patients’ information. **c.** One-dimensional ddPCR fluorescence plots demonstrating expression profiles of the indicated target genes across treatment timepoints in the responder 1-03. **d.** Quantitative longitudinal analysis of *BCL-2* transcript levels (Total copies) in patient 01-03. **e.** PSA response stratified by molecular responder vs. non-responder groups. **f.** Log-transformed correlation of baseline CTC burden (Log10 EpCAM expression) with PSA response.

Longitudinal peripheral blood samples were collected from patients for isolation and transcriptomic profiling of circulating tumor cells (CTCs) using a standardized workflow incorporating microfluidic (Parsortix) enrichment, RNA amplification, and digital droplet PCR (ddPCR) (Fig. 8a, b). This approach enabled high-sensitivity quantification of dynamic changes in *BCL-2* levels, *AR* expression and AR pathway activity (*KLK3, TMPRSS2*), and potential compensatory or ‘bypass’ survival signaling (e.g., *NR3C1*, i.e., GR) (Fig. 8c; Supplementary Fig. S15a, b). CTC collections were also characterized by ddPCR for *TMPRSS2*-ERG fusion (type III) and for relative CTC abundance using the epithelial marker *EPCAM* (Fig. 8c; Supplementary Fig. S15a, b). These CTC molecular profiles were then integrated with serum PSA changes, CTC burden, and prior ARPI exposure to identify potential biomarkers of clinical benefit and resistance (Fig. 8b-f; Supplementary Fig. S15).

Amongst the 10 trial patients, we observed 3 potential “responders” ^70^ (Fig. 8b). For example, CTC collections from patient 1-03, who did not receive any prior ARPI treatment and was on treatment the longest (35 cycles and 980 days; Fig. 8b), were characterized as *TMPRSS2*-ERG fusion negative and showed treatment-related reductions in *BCL-2* and *AR*, AR activity (*KLK3* and *TMPRASS2*) and overall CTC burden (*EPCAM*) (Fig. 8c). Correspondingly, this patient displayed treatment-related decreases in *BCL-2* copy numbers in CTCs (Fig. 8d) and the most significant drop in serum PSA levels (Fig. 8e; Supplementary Fig. S15c). Two additional patients, 02-01 and 02-07, whose CTCs were characterized as *TMPRSS2*-ERG positive (Supplementary Fig. S15b), were treated for 9 and 12 cycles, respectively (Fig. 8b) and exhibited some molecular and clinical responses. Late in treatment cycles and towards end of treatment (EOT), the CTCs in both patients showed decreases in *BCL-2* and *AR* expression, AR activity (*KLK3* and *TMPRSS2*), and overall abundance as evidenced by reduced *EPCAM* and the *TMPRSS2*-ERG fusion (Supplementary Fig. S15b). Correspondingly, these two patients also had PSA responses (Fig. 8e). Nevertheless, earlier treatment discontinuation in both patients was followed by faster PSA rebound (Supplementary Figure S15c) compared to patient 1-03.

In contrast to the above 3 responders, PSA dynamics categorized most other patients as non-responders (Figure 8b, e; Supplementary Fig. S15c). These non-responders (e.g., 1-05, 3-07, 3-02, 3-04) were generally treated with the Enza/venetoclax combination for only 1-3 cycles (Fig. 8b) and exhibited persistent PSA rise (Fig. 8e, Supplementary Fig. S15c). At the molecular levels, although the CTCs from the non-responders did exhibit reduced *AR*, *KLK3* and *TMPRSS2-ERG* mRNA levels at EOT, their CTCs showed minimal baseline (pre-Tx) expression of *BCL-2* that did not appreciably change on therapy (Supplementary Fig. S15a). Intriguingly, unlike in the 3 responders, the *NR3C1* expression levels in the CTCs of non-responders increased at EOD (compare Supplementary Fig. S15a vs. the results in Supplementary Fig. S15b and Fig. 8c).

Finally, to assess how CTC levels might be related to treatment response, we performed a correlation analysis between *EPCAM*-based CTC levels and PSA values across all available time points from all patients. This longitudinal patient-integrated analysis allowed us to track how CTC abundance evolved in relation to PSA dynamics (i.e., treatment response) over time. The results revealed a strong negative correlation (r = –0.76, *P* = 0.001) between the two (Fig. 8f), suggesting that decreased CTC abundance is closely associated with PSA response during treatment.

## DISCUSSION

PCa is inherently heterogeneous, with treatment-naïve tumors already harboring both ARPI-sensitive AR^+^ cells and ARPI-refractory AR^-/lo^ cell populations.^5,8–18,20^ The widespread clinical use of potent second-generation ARPIs has further expanded AR^-/lo^ foci, which now comprise up to 20–30% of metastatic CRPC cases.^5–8,19,21^ Yet, how these phenotypically distinct cell and mCRPC subtypes adapt to survive AR-targeted therapies—and whether they share common, targetable resistance programs—remains poorly understood. Building on our prior classification of CRPC into three phenotypically distinct subtypes—AR^+/hi^, AR^-/lo^, and AR^cyto^ (20) —and the emerging recognition of BCL-2 as a survival factor driving castration resistance, we sought to define how AR^+/-^BCL-2^+/-^ PCa cell subtypes dynamically evolve in response to castration and during progression to CRPC, understand how AR signaling controls BCL-2 expression in different androgen signaling environments, and, critically, determine whether BCL-2 represents a shared, therapeutically actionable resistance mechanism across distinct CRPC subtypes.

Our high-dimensional single-cell imaging analysis across benign prostate (n=123), primary tumors (n=125), CRPC specimens (n=25), and matched xenograft models reveals that the BCL-2⁺ cell subpopulations dynamically evolve and are substantially induced by castration and ARPIs across the PCa continuum. In benign prostate, AR and BCL-2 occupy distinct epithelial compartments—luminal and basal, respectively, reinforcing lineage segregation. Primary tumors remain largely dominated by AR⁺BCL-2⁻ luminal cells, with minimal detection of BCL-2^+^ PCa cells. This is consistent with the known loss of basal cells during tumorigenesis and the concept that the cell(s) of origin for prostatic adenocarcinomas are likely (AR^+^) luminal progenitor cells rather than (AR^-^) basal cells.^5^ However, this balance is disrupted by castration, which induces the evolution and selective expansion of BCL-2⁺ clones across AR^-/lo^, AR^cyto^, and even AR^+/hi^ PCa cell populations. To our knowledge, this report may represent the first detailed mapping of AR^+/-^BCL-2^+/-^ cell subpopulation dynamics during CRPC progression. It should be noted that although some other prosurvival BCL-2 family members such as BCL-xL may be upregulated in castration-resistant PCa cell lines^78^ or involved in promoting PCa cell survival in different contexts^79^, our data suggest that of the 5 prosurvival BCL-2 family members, only BCL-2 is consistently and preferentially induced by castration (Fig. 1). The preferential induction of BCL-2 after castration suggests that BCL-2, unlike other family members, plays a pivotal role in promoting the survival of castrated PCa cells.

Regardless, the castration-induced expansion in BCL-2^+^ PCa cells would inevitably result in dramatic increases of BCL-2 expression in CRPC, as supported by early and recent reports (see Introduction). How does loss of AR signaling lead to BCL-2 upregulation? Upon establishing a consistent inverse relationship between AR activity and *BCL-2* mRNA levels in clinical cohorts, we discovered that AR directly represses *BCL-2* transcription via binding to multiple ARBS across the *BCL-2* genomic region, especially the ARBS1 around the promoter, and castration and ARPI treatment block AR engagement to most ARBS leading to de-repression of *BCL-2* transcription. This establishes, for the first time, a direct transcriptional link between AR loss or inhibition of AR signaling and BCL-2 de-repression and upregulation. It is fascinating that compared to CRPC_AR subtype, the AR^-/lo^ subtypes of CRPC, i.e., CRPC_Wnt and CRPC_NE, maintain notably more open chromatin regions (i.e., ATAC peaks) across the *BCL-2* genomic region in association with much more pronounced *BCL-2* transcription (Supplementary Fig. S13). This integrated analysis of the chromatin landscape and *BCL-2* transcription in well-defined subtypes of CRPC with contrasting AR signaling activity^76^ suggests an interesting possibility that the increased BCL-2 expression in CRPC may result not only from loss of AR-mediated *BCL-2* repression, but also from active *BCL-2* transcription mediated by other transcription factors such as NF-κB, STAT3 and E2F1^33,36,42^ bound to these newly available chromatin loci. Our ongoing work is testing this exciting possibility.

Considering the well-reported association of BCL-2 with castration resistance and upregulation of BCL-2 expression in CRPC and NEPC (see Introduction), what are the significance and relevance of our high-content analysis of AR^+/-^BCL-2^+/-^ cell subpopulation dynamics and detailed molecular dissection of AR regulation of BCL-2 expression? These results allowed us to pinpoint several BCL-2-expressing cell populations in patient CRPC, including AR^-^BCL-2^+^, AR^+^BCL-2^+^ and AR^cyto^BCL-2^+^ PCa cells, that manifest no or attenuated sensitivities to ARPI but common sensitivity to BCL-2i. It is remarkable that the AR^-^BCL-2^+^, AR^+^BCL-2^+^ and AR^cyto^BCL-2^+^ PCa cell subpopulations identified in patient CRPC are representatively modeled by our unique LAPC9-AI, LNCaP-AI and LAPC4-AI xenograft system, respectively. Subsequent therapeutic studies in these xenograft (and organoids) models demonstrated that the BCL-2i ABT-199 exhibits single-agent efficacy in all three BCL-2 overexpressing AI models whereas Enza and ABT-199 combination manifests the greatest potency in inhibiting the AR^+^BCL-2^+^ (i.e., double-positive) LNCaP-AI. These results functionally validate BCL-2 as a shared therapeutic target in diverse subtypes of CRPC.

Building on these mechanistic insights and therapeutic outcomes in defined CRPC models, we conducted a Phase Ib trial evaluating Enza combined with venetoclax in mCRPC.^70^ Although the regimen was well tolerated, clinical activity was overall modest, likely due to Enza-induced CYP3A4 that accelerated venetoclax clearance and limited its therapeutic exposure.^70^ Nevertheless, PSA responses together with molecular changes in CTCs identified 3 potential responders (1-03, 2-01 and 2-07), who were all on the combination treatment for multiple cycles and shared several common characteristics. *First*, 2 of the 3 responders were ARPI-naïve, suggesting that the Enza/venetoclax combination might derive better clinical efficacy when given together at the outset of treatment (rather than sequentially after Enza failure). *Second*, all 3 patients appeared to have higher baseline (i.e., pre-treatment) *BCL-2* levels in their CTCs than the non-responders, suggesting that the response to BCL-2i in PCa patients, understandably, is dictated by the tumor expression of BCL-2 and future BCL-2i clinical trials should stratify PCa patients on BCL-2 expression status and levels as has been done in breast cancer patients.^80–82^ *Third*, all 3 responders showed concurrent suppression of *AR* and *BCL-2* expression and AR activity with evidence of CTC clearance. Last, unlike the non-responders, the CTCs in the 3 responders did not show increased expression of *NR3C1*, which encodes the GR protein. GR has been well reported to drive castration resistance^20,83^ and indeed, our studies here indicate a synergistic effect of GR antagonist and Enza in suppressing the AR^cyto^BCL-2^+^ LAPC4-AI organoids.

By integrating high-dimensional imaging analysis at single-cell resolution, in-depth mechanistic dissection, extensive therapeutic studies in defined preclinical models, and CTC-based correlative studies associated with a Phase Ib clinical trial, this investigation highlights therapy-induced cellular diversification, exposes the 3 BCL-2-upreulated CRPC subtypes with varying AR expression and signaling intensity (i.e., AR^+^BCL-2^+^, AR^-/lo^BCL-2^+^, and AR^cyto^BCL-2^+^), advances our mechanistic understanding of AR-mediated repression and ARPI-triggered activation of BCL-2 transcription, and, critically, validates BCL-2 as a therapeutic target across distinct CRPC subtypes. On the other hand, our study has several limitations and raises some important questions. First, lack of serial samples at progressive stages of castration resistance in our imaging analysis precluded us from obtaining a dynamic picture of which subpopulation(s) of AR^+/-^BCL-2^+/-^ PCa cells emerge first and how these subpopulations inter-convert and transition during full CRPC development. Second, although we now know that AR directly represses *BCL-2* transcription under androgen-proficient conditions, there is little understanding on how BCL-2 is expressed and even upregulated in AR^+/hi^ subtype of CRPC such as the LNCaP-AI model. Third, we provided evidence that the AR^-/lo^ subtype of CRPC (such as LAPC9-AI and CRPC_Wnt and CRPC_NE) may upregulate BCL-2 not only through the loss of AR-mediated repression but also through increased chromatin accessibility, but what TFs might be actively transcribing the *BCL-2* via these open chromatin loci in the AR^-/lo^ CRPC cells? Fourth, the AR^cyto^BCL-2^+^ CRPC cells, which can be recapitulated in the LAPC4-AI model both in vivo and in vitro, clearly exist in patient CRPC but are probably among the least understood. It will be interesting to investigate whether persistent BCL-2 expression in AR^cyto^ CRPC cells is caused by the loss of nuclear AR-mediated BCL-2 repression, by signaling functions of cytoplasmic AR, and/or by other nuclear TFs such as NF-κB, STAT33 and E2F1^33,36,42^. Finally, the VCaP-AD model displays an AR^+^BCL-2^+^ phenotype while the VCaP-AI model, intriguingly, presents an AR^cyto^BCL-2^-^ phenotype. VCaP is the only model in our study that has both the TMPRSS2–ERG fusion and endogenous AR-V7, two clinically relevant alterations that frequently co-occur in CRPC patients. While both are known to reshape transcription and chromatin, their impact on BCL-2 remains unclear. Such insights could expand the clinical utility of BCL-2–targeted therapies in genomically defined subsets of PCa patients.

Ultimately, these findings should facilitate the translation of BCL-2i to the clinic to treat PCa patients. Our CTC-based correlative studies demonstrate that effective therapeutic responses to the ARPI/BCL-2i combination in mCRPC patients may require concurrent suppression of AR and BCL-2 signaling and clearance of CTCs whereas treatment resistance is marked by persistent CTC burden and associated with activation of alternative survival pathways such as GR. Looking forward, precision stratification of PCa patients based on AR and BCL-2 status could optimize therapeutic efficacy. Integration of liquid biopsy transcriptomics could identify patients with BCL2-driven disease. Additionally, substituting Enza with agents like abiraterone—an AR inhibitor with strong CYP3A4 inhibition—may enhance BCL-2i exposure and drive more complete clinical benefit in PCa patients receiving the ARPI/BCL-2i treatment.

## Materials &Methods

### Cell lines, cell culture, xenograft lines, animals, and animal protocols

LNCaP, LAPC4 and VCaP cells were purchased from the American Type Culture Collection (ATCC) and cultured in RPMI-1640 medium (for LNCaP) or DMEM (for LAPC4 and VCaP) plus 10% heat-inactivated fetal bovine serum (FBS) plus antibiotics. LAPC4 and VCaP cells were cultivated in poly L-lysine coated plates. LAPC4 and LAPC9 xenograft lines were initially provided by Dr. Robert Reiter (UCLA) and have been used extensively in our previous studies.^e.g.,14,17,18, 20^ These PCa cell and xenograft lines were authenticated regularly in our institutional CCSG Cell Line Characterization Core and examined to be free of mycoplasma contamination. Immunodeficient NOD/SCID (non-obese diabetic severe combined immunodeficiency) and NOD/SCID-IL2Rγ^-/-^ (NSG) mice were obtained from the Jackson Laboratory, and the breeding colonies were maintained in standard conditions in our animal core. All animal-related studies in this study have been approved by our Institutional Animal Care and Use Committee (IACUC) at Roswell Park Comprehensive Cancer Center (animal protocol# 1328M, 1331M).

### Establishment of AD and AI (CRPC) xenografts

Briefly, AD (i.e., androgen-dependent) xenograft tumors, LNCaP, VCaP, LAPC4, and LAPC9, were routinely maintained in intact immunodeficient NOD/SCID or NSG mice. To establish the castration-resistant or androgen-independent (AI) xenograft lines, parental AD tumor cells were purified, mixed with Matrigel, injected subcutaneously and serially passaged in surgically castrated male immunodeficient mice. The LAPC4 and LAPC9 AD/AI xenograft lines were maintained in intact and castrated male NOD/SCID mice whereas LNCaP and VCaP AD/AI xenograft lines were passaged in intact and castrated male NSG mice, respectively. Xenograft tumors that became castration-resistant were termed primary (1°) CRPC (or AI) and the AI tumors that became Enza-resistant were termed secondary (2°) CRPC.^20^

To purify human PCa cells, xenograft tumors were harvested, chopped into small pieces (∼1 mm^3^), and digested with Accutase (Sigma, A6964) for 30 min at room temperature (RT) under constant rotations. Single cells were collected via a pre-wetted 40-μm strainer and further purified on Histopaque-1077 (Sigma) gradient to deplete debris and dead cells.

### Antibodies

Multiple antibodies were used in Vectra-based qmIF (quantitative multiplex immunofluorescence), imaging mass cytometry (IMC), regular Western blotting (WB), quantitative WB (WES), immunofluorescence (IF) and immunohistochemistry (IHC) studies. The basic information for these antibodies is summarized in Supplementary Table S1. Main antibodies used in Vectra qmIF analysis included AR, BCL-2 and pan-Cytokeratin. The antibodies used in IMC studies were conjugated to metals using the MaxPar X8 Multimetal Labeling Kit (Fluidigm) according to the manufacturer’s instructions. Before testing antibodies, manufacturer’s website and antibody databases were used for choosing antibodies for targets of interest. Antibodies were initially tested using unconjugated antibodies in regular IF staining of lymph node, spleen, and breast cancer sections. Antibodies that revealed expression patterns consistent with the literature were chosen for metal conjugation. After the conjugation, another round of testing was undertaken using IMC with breast cancer sections and antibodies that showed staining patterns consistent with the literature and with sufficient signal intensity were utilized.

### Primary human PCa (HPCa) and CRPC whole-mount (WM) tissue sections and CRPC tissue microarrays (TMAs) used in the study

In our imaging analyses, we utilized a total of 123 benign prostate tissues, 125 primary HPCa, and 25 CRPC specimens (Supplementary Fig. S1a, b). The benign tissues comprised 120 specimens in TMA-1 and TMA-2 as well as 3 WM benign tissues adjacent to the corresponding HPCa; the 125 treatment-naïve HPCa specimens consisted of 121 tumor sections in TMA-1 and TMA-2 and 4 WM HPCa sections; and the 25 CRPC specimens included 20 in TMA-1 and 5 WM CRPC sections (Supplementary Fig. S1b). Both TMA-1 and TMA-2 were made in Dr. J. Huang’s laboratory, with TMA-1 consisting of 5 benign prostate tissues, 6 primary tumors, and 20 CRPC (2 cores per sample) and TMA-2 consisting of 115 matched benign and tumor tissues (3 cores/sample). The 3 benign and 4 primary tumor WM sections were made in our lab while the 5 CRPC WM slides were provided by Dr. J. Huang (Supplementary Fig. S1b). Formalin-fixed paraffin-embedded (FFPE) sections were cut from these TMAs and WM samples and used for Vectra and IMC studies, and relevant information on patient samples was summarized in Supplementary Table 2.

### Chemicals, drugs, key assay kits and reagents, primers, and probes

All relevant information is summarized in Supplementary Table S3, along with their sources, catalog numbers, and uses. Compounds were chosen based on their relevance to pathway inhibition, therapeutic targeting, or the experimental design and were reconstituted and stored according to the manufacturers’ instructions.

### Regular western blotting (WB) and automated quantitative WB (WES)

For regular WB, total protein extracts were obtained from cell or tissue samples using Pierce RIPA buffer (ThermoFisher Scientific, USA) according to manufacturer’s protocol. Protein concentrations were determined using Pierce BCA Protein Assay Kit (ThermoFisher Scientific, USA). 40 μg of total protein were separated in Bolt 4-12% Bis-Tris Plus Gels (ThermoFisher Scientific, USA) and transferred to Hybond P 0.45 PVDF membranes. Membranes were blocked with 5% BSA in TBST (FisherSci, USA) and probed overnight at 4°C with primary antibodies (Supplementary Table S1). Secondary antibodies were HRP-conjugated anti-rabbit IgG or anti-mouse IgG (CST 7074, 7076, 1:10000). Protein bands were revealed using Luminata Classico or Luminata Forte HRP substrate (MilliporeSigma, USA) and detected using Chemidoc Imaging system (Bio-Rad, USA)

In some studies, we also employed the automated the Simple Western WES^TM^ immunoassays (https://www.bio-techne.com/brands/proteinsimple), which take place in capillaries and separate proteins by size as they migrate through stacking and separation matrix. The separated proteins are immobilized to the capillary wall via a proprietary, photoactivated capture chemistry. The target protein was identified using a primary antibody and immunoprobed using an HRP-conjugated secondary antibody and chemiluminescent substrate. The resulting chemiluminescent signal is displayed as traditional virtual blot-like image and electropherogram. Quantitative results such as M.W, signal intensity (area), % area, and signal-to-noise for each immunodetected protein are presented in the results table automatically.

### Immunofluorescence (IF) in cultured cells

Cells were seeded at 40,000 cells per well in an ibidi 8-well chamber slide and allowed to adhere overnight. The following day, cells were incubated with MitoTracker™ (e.g., 100 nM in complete medium) at 37°C for 30 minutes to label mitochondria, followed by gentle washing with warm PBS. Cells were then fixed with 4% paraformaldehyde in PBS for 15 minutes at RT, permeabilized with 0.1% Triton X-100 in PBS for 10 minutes and blocked with 5% BSA in PBS for 30 minutes. For single IF staining, cells were incubated either with Alexa Fluor 647-conjugated anti-AR antibody for 1 hour at RT or with anti-BCL-2 antibody (Supplementary Table S1) for 4 hours at RT, followed by appropriate Alexa Fluor-conjugated secondary antibody incubation. For co-staining, cells were sequentially incubated with anti-BCL-2 antibody and secondary antibody, followed by Alexa Fluor 647-conjugated anti-AR antibody. Control wells included a no-antibody condition as well as isotype controls, where rabbit IgG replaced the AR antibody and normal mouse IgG replaced the BCL-2 antibody. After final PBS washes, nuclei were optionally counterstained with DAPI, and slides were mounted using antifade medium and imaged using Keyence BZX-700 microscope.

### IHC in FFPE sections

The slides were deparaffinized and rehydrated according to common protocols. Antigen retrieval was performed by boiling the slides in citrate buffer (pH=6.0) for 10 minutes using microwave. Sections were blocked using solution of 10% FBS (Gibco, USA) and 1% BSA in TBS for 2 hours at RT and probed with AR antibody (CST D6F11, 1:100 in TBS with 1% BSA) overnight at 4°C. Slides were rinsed with 0.0025% Triton-X100 (Fisher Scientific, USA) in TBS, probed with secondary HRP-conjugated anti-rabbit IgG (CST 7074, 1:1000 in TBS with 1% BSA) for 1 hour at RT and developed with 3,3-diaminobenzidine (DAB) kit (Vector Laboratories, USA). Separate slides were counterstained with hematoxylin (Fisher Scientific, USA). After staining, slides were rinsed with DI water, dehydrated, and mounted using toluene (Fisher Scientific, USA). The slides were scanned using Aperio ScanScope imaging system (Aperio Technologies, Vista, CA, USA) and a 40x objective. Images were analyzed using the ScanScope software.

### Vectra-based quantitative multiplex IF (qmIF) staining and data analysis using inForm

qmIF analysis with tyramide signal amplification was performed in FFPE slides of TMAs, WM CRPC patient samples and PCa AD/AI xenografts. The qmIF antibody panel (Supplementary Table S1) consisted of AR, BCL-2 and pancytokeratin AE1/AE3 and the FFPE sections (5 mm) stained with the automated Vectra platform. Antibody staining in the qmIF panel was compared to that in single stained slides on control tonsil and prostate tissue. Staining patterns did not differ between the single- and multiplex scanned slides. Each marker was also validated with conventional IHC. Slides were scanned and analyzed with the Vectra Polaris (Akoya Biosciences).

After staining, slides were imaged using the Vectra 3.0 automated imaging system (Akoya Biosciences). First, whole-slide scans were made at 10x magnification. Subsequently, multispectral image scan was processed to single images using Phenochart slide viewer (Akoya Biosciences). Multispectral library slides were created by staining a representative sample with each of the specific dyes. The multispectral library slides were unmixed into eight channels using inForm software (version 2.4): DAPI, CK OPAL 480, AR OPAL520, BCL2 OPAL570 and Auto Fluorescence and exported to a multilayered qTIFF file. The multilayered TIFFs were fused with Phenoimager software (version 3.0) to create one file for each sample. Cells were phenotyped as pancytokeratin AE1/AE3 positive (CK+) or negative (CK-) cells. The cell density for each cell subset was further assessed for CK+ epithelial compartment and expressed as number of AR^+/-^ BCL-2^+/-^ cells/mm^2^.

For Vectra data analysis, we employed the inForm Cell Analysis program, which enabled us to define the biology of interest within a tissue section. It phenotypes cells based on their biomarker expression within cells, nuclei and membranes. *inForm* Tissue Finder added exceptional functionality to inForm Cell Analysis. It automated the detection and segmentation of specific tissue compartments through powerful algorithms. Automation provided consistent reproducible results and enabled comparative studies of multiple markers and multiple specimens.

### Imaging mass cytometry (IMC)

Data acquisition was performed on a Helios time-of-flight mass cytometer (CyTOF) coupled to a Hyperion Imaging System (Fluidigm). Selected areas for ablation were larger than the actual area of interest to account for loss of overlapping areas among sections due to cumulative rotation. The selected area ablated per section was around 1 mm^2^. Laser ablation was performed at a resolution of approximately 1 µm with a frequency of 200 Hz with estimated acquisition time of 1 mm^2^ h^−1^. To ensure performance stability, the machine was calibrated daily with a tuning slide spiked with five metal elements (Fluidigm). All data were collected using the commercial Fluidigm CyTOF software v.01.

For antibody validation, the AR, CK antibodies were purchased from Fluidigm R (https://www.fluidigm.com). The carrier-free BCL-2 antibody was conjugated Nd146 using the Maxpar X8 metal conjugation kit following manufacturer’s protocol (Fluidigm 201300). Post-conjugation, antibody specificities were again tested using immunofluorescent staining, followed by titration in the IMC platform.

For antibody staining and image acquisition, 2-4 μm FFPE sections were stained with an antibody cocktail (Supplementary Table S1) containing all (AR, BCL-2, Pan CK) antibodies. Briefly, tissue sections were de-paraffinized with xylene and subjected to sequential rehydration from 100% ethanol to 70% ethanol before being transferred to PBS. Heat-induced antigen retrieval was performed at 95°C for 30 min in Tris/EDTA buffer (10 mM Tris, 1 mM EDTA, pH9.2). Slides were cooled to RT and subsequently blocked with PBS+3%BSA for 1h at RT. Meanwhile, the antibody cocktail was prepared in PBS+1%BSA buffer, with appropriate dilutions for each of the antibodies (Supplementary Table S1). Each slide was incubated with 100 μl of the antibody cocktail overnight at 4°C. The next day, slides were washed 3 times with PBS and labeled with 1:400 dilution of Intercalator-Ir (Fluidigm 201192B) in PBS for 30 min at RT. Slides were briefly washed with H2O three times and air dried for at least 30 min before IMC acquisition. All IMC operation was performed following Fluidigm’s operation procedure. Briefly, following daily tuning of IMC, image acquisition was carried out following manufacturer’s instruction at a laser frequency of 200 Hz. 1,000 μm x 1,000 μm regions around islets were selected based on bright field images.

For IMC data analysis, the CellProfiler (version 3.0.0) analysis pipeline was used to run complex image on batches of several hundreds of images. The software requires the use of analysis modules which run image processing algorithms, threshold-based segmentation, calculations, and file processing (https://star-protocols.cell.com/protocols/1361). To quantify each marker, we developed an IMC segmentation and analysis pipeline based on the ‘IMC Segmentation Pipeline’ repository created by the Bodenmiller group.^84^ Cell Profiler was used to prepare the images for segmentation based on an overlay of all markers to identify cytoplasm/membrane, nuclei (iridium), and non-cellular space. This step is independent of signal intensity for markers, as it captures overall cellular morphometry. These data were subsequently loaded into HistoCAT for quantification and downstream analysis of single cell populations. Dimensionality reduction using the tSNE algorithm of HistoCat allowed for visualization of the multiplexed, single cell dataset in two dimensions. tSNE was selected as it has been most widely used for clustering of IMC data. Comparison of AD and AI tumors via tSNE highlighted global differences in the distribution and grouping of single cell data. A heatmap plot visualizing relative signal intensity per cell by individual markers enabled phenotypic classification of the four AR^+/-^BCL-2^+/-^ cell meta-clusters. If a panel of slides is stained using IMC for a discrete marker, the intensity of that marker across cells in an individual section and across different ROIs obtained from the same type of tissue, stained using the same method, can be measured and compared.

### Establishment of castration-resistant and castration/Enza-resistant LAPC4 and LNCaP cells

We established the castration-resistant and dual castration/Enza-resistant LAPC4 cell models (Fig. 4; Supplementary Fig. S10) to further study the AR^cyto^ subtype of CRPC and the therapeutic responses. To this end, we cultured regular LAPC4-AD cells long-term (4 months) in charcoal dextran stripped serum (CDSS) medium to develop castration-resistant LAPC4 (LAPC4-CR or LAPC4-AI) cells, which were subsequently exposed to either 20 mM or 100 mM Enza-containing CDSS medium for 1 month resulting in the LAPC4-Enza(20)-R or LAPC4-Enza(20)-R models (Supplementary Fig. S10a).

Analogously, LNCaP cells were cultured in CDSS-containing medium for 8 weeks to generate castration-resistant LNCaP-CR cells, followed by Enza exposure (50 µM, 2 weeks) to derive LNCaP-C/Enza-R cells (Supplementary Figure S14e).

### Drug treatment of 2D cell cultures and 3D organoids

Drugs such as enzalutamide (Enza; Apex Bio, A3003) and ABT-199 (Apex Bio, A8194) were dissolved in DMSO. Control samples in all experiments were treated with vehicle (DMSO) only and vehicle concentration in growth medium did not exceed 0.2%. For drug treatment in 2D cultures, cells were seeded in 24-well ultra-low attachment plates (Corning, USA) at 2,000 cells/well. After 7 days cell clusters were harvested and disrupted by mild trypsinization, pelleted, and resuspended in PBS. Cell viability was assessed by counting using a Countess automated cell counter (ThermoFisher Scientific, USA) in presence of 0.4% Trypan Blue. For organoids-based drug assays, xenograft tumor cells were isolated and seeded into 384-well microtiter plates at 10,000 cells per well in F12 advanced DMEM media supplemented with 10% Matrigel. Drugs in the same culture medium were added in half-log dilutions starting at 0.1 μM. Cells were incubated at 37°C (5% CO2) for 6 days and the effects of drugs on organoids viability were determined using Resazurin cell viability assays (abcam). Data were normalized to “Control” samples and presented as mean ± SD. Kruskall-Wallis H-test was applied against corresponding concentrations of drugs (e.g., Enza) that yielded the fraction affected (FA) when acted alone and in combination with BCL-2 and AR inhibitors. The drug-drug interaction studies were assessed using Compusyn software https://compusyn.software.informer.com/1.0/ and synergy finder www.synergyfinderplus.org.

### ChIP-qPCR analyses of the AR-binding sites (ARBS) on the BCL-2 genomic locus

ChIP was performed in LNCaP, LAPC4, and LAPC9 AD/AI xenograft tissue lysates using the ChIP-IT High Sensitivity Kit (Active Motif, #53040), following the manufacturer’s protocol. Chromatin fragments containing the *BCL-2* regulatory regions were immunoprecipitated using ChIP-grade anti-AR antibody (EPR179 # ab108341). An anti–RNA Polymerase II monoclonal antibody and normal IgG, both included in the kit, were used as positive and negative ChIP controls, respectively. Input samples—representing 2% of the total chromatin prior to immunoprecipitation—were processed in parallel and used for normalization. After reverse crosslinking, immunoprecipitated DNA and input samples were analyzed by qPCR using ARBS– specific primers targeting the *BCL-2* locus (Supplemental Table 3). IgG pulldown samples served as negative controls to confirm the specificity of enrichment.

To validate AR occupancy at the *BCL-2* locus, we designed primers targeting three high-confidence ARBS (i.e., ARBS1–3) and one negative (Neg) control region based on AR ChIP-seq profiles in androgen-dependent (AD; GSM699631) versus androgen-independent (AI; GSM699630) LNCaP cells (Figure 5g-h). Candidate peaks were selected using MACS (v1.4.2) peak calling with a stringent p-value cutoff (1e-5) and were visualized on the UCSC Genome Browser (hg38) for cross-comparison with histone marks and enhancer annotations. The ‘Neg’ control region lacking AR binding (chr18:63,300,951–63,301,529) was selected from a neighboring genomic region. The following genomic coordinates (hg38) were used as the qPCR amplicon targets: ARBS1: chr18:63,321,543–63,321,592 (50 bp); ARBS2: chr18:63,202,549–63,202,622 (74 bp); ARBS3: chr18:63,200,006–63,200,051 (46 bp). Primers and probes were designed by Bio-Rad (Supplementary Table S3) to specifically amplify ARBS1–3. Probes were labeled with FAM for ARBS1–3 and HEX for Neg control to enable clear discrimination of specific AR-binding events. Primer-probe sets were optimized to amplify peaks corresponding to AR occupancy identified by ChIP-seq.

In brief, 10 × 10⁶ single cells isolated from the AD/AI PCa xenograft tissues were fixed in formaldehyde for 15 minutes, and chromatin was sheared using the Covaris E210 focused-ultrasonicator for a total of 16 minutes. Shearing was performed in 12 × 12 mm, 1 mL glass tubes containing a fiber insert (Covaris, cat. no. 520080) placed in a compatible tube holder (Covaris, cat. no. 500276), using the following instrument settings: 20% duty cycle, intensity 8, and 200 cycles per burst, applied in sixteen 1-minute treatment cycles. After shearing, 10 μL of the 2 mL sonicated chromatin was reserved as input, and the remaining chromatin was incubated overnight at 4°C with 4.0 μg of anti–AR antibody or normal rabbit IgG (Cell Signaling Technology, Cat. #2729). For qPCR analysis, 5 μL of immunoprecipitated DNA was used as a template in a 10 μL total reaction volume using TaqMan™ Advanced PCR Master Mix (Thermo Fisher Scientific) and target-specific TaqMan probes, prepared in 384-well plates. Quantitative PCR was performed on a QuantStudio™ 6 Flex Real-Time PCR System (Applied Biosystems) under standard cycling conditions: initial activation at 95°C for 10 minutes, followed by 40 cycles of 95°C for 15 seconds and 60°C for 1 minute. Data were analyzed by calculating fold enrichment over IgG control.

### Dual luciferase assays to assess ARBS functions on the BCL2 promoter

The ARBS1-WT (wild-type) and ARBS1-Mut (mutant) sequences (Figure 5j) were cloned into the pCMV-Luc vector, and the corresponding luciferase reporter constructs were generated by OriGene (catalog numbers CW310528 and CW310529). LNCaP cells were seeded in 24-well plates (3 × 10⁴ cells/well) and co-transfected with 1 μg of the respective firefly luciferase constructs along with a Renilla luciferase control plasmid for normalization, using Lipofectamine 3000 (Invitrogen; catalog number L3000008) and according to manufacturer’s instructions. As a positive control, one group was treated with 10 nM DHT for 12 hours prior to measurement. Luciferase activity was measured 24–48 hours post-transfection using a dual-luciferase assay. Cells were lysed, and 100 μL of firefly luciferase working solution was added; luminescence was measured immediately followed by the addition of 100 μL of Renilla luciferase working solution. The ratio of firefly to Renilla luminescence was calculated to determine relative promoter activity. Statistical significance was assessed using the Kruskal–Wallis test, with *p<* 0.05 considered significant.

### Therapeutic experiments in LNCaP-AD/AI, LAPC4-AI and LAPC9-AI xenografts

Basic experimental protocols for in vivo therapeutic studies were described previously.^20^ Briefly, for all AI xenograft studies, NOD/SCID or NSG male mice (8–10 weeks old) were castrated on day 0. After 7–10 days, 250,000 CRPC cells from various AI xenograft models^20^ (i.e., LNCaP-AI, LAPC4-AI and LAPC9-AI) were injected subcutaneously in Matrigel. Once tumors became palpable, mice were randomized into treatment groups, and drugs were administered by oral gavage: ABT-199 at 50-100 mg/kg five times per week and Enza at 30 mg/kg three times per week, for a total treatment of ∼3-6 weeks (depending on models). All drugs were dissolved in DMSO and formulated in 10% DMSO, 50% PEG300, and 10% Tween-80. Tumor dimensions were measured weekly using calipers, and volumes were calculated as ½ × (length × width²). Animal body weights and health were monitored throughout the treatment period. At the study endpoint, tumors were harvested for analysis of incidence, weight, and gross morphology.

Specifically, we evaluated the therapeutic efficacy of ABT-199, alone or in combination with Enza in models of distinct CRPC subtypes (Fig. 7, Supplementary Fig. S14). In the AR^Cyto^ LAPC4-type CRPC model (Fig. 7b), mice were assigned to four groups: 1) vehicle control (n = 12), 2) Enza (30 mg/kg; n = 10), 3) ABT-199 (50 mg/kg; n = 12), and 4) combination (Enza 30 mg/kg + ABT-199 50 mg/kg; n = 10). In the AR^-/lo^ LAPC9-type CRPC (Fig. 7g-j; Supplementary Fig. S14j-l), mice were treated with either vehicle control, ABT-199 or AT-101 for up to ∼4 weeks. In an independent LAPC9-AI therapeutic study (Fig. 7k), castrated male NOD/SCID mice bearing LAPC9-AI tumors were treated with vehicle control (n=10) or ABT-199 (100 mg/kg; n=12) for 3 weeks. Finally, we conducted several independent therapeutic studies in AR^+^BCL-2^-^ LNCaP-AD and AR^+/hi^BCL-2^+^ LNCaP-AI tumor models (Fig. 7c-f) as previously described.^20^

### RNA isolation and qRT-PCR analysis

Total RNA was extracted using RNeasy Mini Kit (Qiagen) according to the manufacturer’s instructions. qRT-PCR was performed using a CFX Connect Real-Time PCR Detection System (Bio-Rad). qPCR data were normalized to housekeeping gene TBP and HPRT1 expression levels.

### Integrated analysis of ATAC-seq, AR ChIP-seq, and transcriptomic (RNA-seq and microarray) data

#### Data Acquisition

We sourced ATAC-seq, AR ChIP-seq, RNA-seq and microarray datasets from publicly available repositories, with a focus on benign prostate tissue, primary PCa, CRPC, and related PCa cell lines. Dataset sources, accession identifiers, and metadata such as sample type, sample size, and platform are detailed in Supplementary Table S4.

#### Transcriptomic Data (RNA-seq and Microarray)

Normalized gene expression datasets were obtained from the NCBI Gene Expression Omnibus (GEO; RRID:SCR_005012; https://www.ncbi.nlm.nih.gov/geo/), cBioPortal for Cancer Genomics (RRID:SCR_014555; https://www.cbioportal.org/), and UCSC Xena Functional Genomics Portal (RRID:SCR_018938; https://xenabrowser.net/). Cohorts included benign prostate tissues, untreated and treated primary PCa (e.g., pre-/post-nADT, pre-/post-Enza), CRPC, and PCa cell lines.

#### Microarray Data Processing

Two microarray datasets were included. For GSE89050,^71^ normalized data for FACS-purified basal, luminal, and luminal progenitor cells from benign prostate tissue were directly downloaded from GEO. For GSE21034,^85^ normalized expression values for benign prostate tissues, primary PCa and metastatic PCa tissues were retrieved from cBioPortal. For both datasets, expression values were log₂-transformed for downstream visualization and analysis. No additional normalization steps were applied beyond the original authors’ processing.

#### RNA-seq Data Processing

RNA-seq expression values (FPKM, TPM, or DESeq2-normalized counts) were obtained directly from the original data sources. When applicable, fold-change (FC), 95% confidence intervals (CIs), p-values, and false discovery rates (FDR) were either extracted from the original publications or computed using DESeq2 (version 1.28.1; RRID:SCR_015687) for datasets with raw counts—including the Sharma pre-/post-nADT cohort (GSE111177), LNCaP-ARKO/AR datasets, and LNCaP-derived secondary CRPC/primary CRPC/AD samples (GSE88752). For dataset without raw counts (i.e., the Alumkal pre-/post-Enzalutamide cohort), fold-changes were calculated from TPM values, and paired comparisons were assessed using the one-sided Wilcoxon signed-rank tests. No additional normalization or batch correction was applied beyond the processing inherent to DESeq2 or the source datasets, unless otherwise specified.

#### ChIP-Seq Data Acquisition and Analysis

Reads were mapped to the human genome (hg38) using Bowtie (version 1.1.2)^86^ with the following parameters: “-v 2 -m 1 --best --strata”. To avoid PCR bias, only one copy of multiple reads mapped to the same genomic position was retained for further analysis. ***Peak Calling:*** For each ChIP-Seq sample, peaks were identified using MACS (version 1.4.2)^87^ by comparing against the corresponding input sample. A window size of 300 bp was used, and a p-value cutoff of 1e-5 was applied. The peaks that overlapped with ENCODE blacklisted regions^87^ were removed. ***Differential peak analysis:*** For each differential comparison, peaks from all the involved samples were merged, and the number of reads within these merged peaks was counted for each sample. Merged peaks with less than 10 reads in all samples were removed. The resulting count table was used to identify differential peaks using the R/Bioconductor package edgeR.^89^ The numbers of reads within the common peaks of all samples were used as the library sizes in edgeR. Peaks with a false discovery rate (FDR) ≤ 0.05 and a fold change ≥ 1.5 or 2 were identified as differential peaks and presented in heatmap plots. ***Signal Track:*** Each read was extended by 150 bp to its 3’ end. The count of reads covering each genomic position was multiplied by 1×10^7^ divided by the library size used in edgeR. These values were then averaged over a 10 bp resolution. The resulting averaged values were displayed using the Integrative Genomics Viewer (IGV).^90^

#### ATAC-Seq Data Acquisition and Analysis

Adapter sequences were removed from the 3’ ends of reads using Trim Galore! (version 0.6.5; Babraham Bioinformatics. https://www.bioinformatics.babraham.ac.uk/projects/trim_galore/) and cutadapt (version 2.8).^91^ The reads were then mapped to the human genome (hg38) using Bowtie (version 1.1.2) with the following parameters: “--allow-contain --maxins 2000 -v 2 -m 1 --best --strata”. To avoid PCR bias, only one copy of multiple fragments mapped to the same genomic position was retained for further analysis. After the removal of fragments from chrM, for each fragment, the 5’ end was offset by +4 bp and the 3’ end was offset by −5 bp to adjust both ends to represent the center of a transposon binding event. ***Peak Calling:*** For each sample, peaks were identified using MACS2 (version 2.2.7.1) without using any control. Each binding event (i.e., the 5’ or 3’ end of a fragment) was smoothened by 73 bp (i.e., extended 36 bp upstream and 36 bp downstream from the event center). MACS2 was configured to call peaks from the pile-up of smoothened binding events, and a q-value cutoff of 0.05 was applied. The peaks that overlapped with ENCODE blacklisted regions were removed. ***Signal Track:*** each transposon binding event was smoothened to a length of 73 bp, spanning from −36 bp to +36 bp around the center. The count of binding events covering each genomic position was multiplied by 1×10^7^ divided by the total number of binding events. These values were then averaged over a 10 bp resolution. The resulting averaged values were displayed using the Integrative Genomics Viewer (IGV). Pearson’s correlation was used to assess the relationships between chromatin accessibility, AR binding, and *BCL-2* expression.

### Phase Ib clinical trial of treating mCRPC patients with enzalutamide and venetoclax combination

The NCT03751436 trial was a phase Ib open label single-arm, single-center study of Enza with the BCL-2i venetoclax in patients with mCRPC.^70^ A total of 10 patients were enrolled, starting with a standard dose of Enza (160 mg/d) and 3 dose levels (DL) of venetoclax at 400 mg/d (DL1), 600 mg/d (DL2) and 800 mg/d (DL3). The primary objectives of this phase Ib study were to: 1) characterize the safety and tolerability profile and 2) determine the dose-limiting toxicity (DLT), maximum tolerated dose (MTD) and recommended phase II dose (RP2D) of the Enza and venetoclax combination in patients with mCRPC. We also analyzed the pharmacokinetic (PK) profiles of Enza and venetoclax when given in combination to confirm whether drug levels were in the therapeutic range. For patient accrual, treatment and adverse events, clinical outcomes, and major PK parameters, refer to ref. 70.

### Circulating tumor cell (CTC) enrichment and characterization

CTCs were isolated from patient blood samples using the Parsortix system (ANGLE plc’s Parsortix™). Blood was processed according to the manufacturer’s protocol, enabling size- and deformability-based enrichment of CTCs while minimizing contamination by RBCs and WBCs. Following enrichment, the captured CTCs were harvested directly from the Parsortix cassette using ANGLE’s cell harvest protocol. Cell pellets were immediately stored in the extraction buffer provided in the NanoPure RNA extraction kit or processed directly for downstream analysis.

To molecularly characterize the enriched CTC collections, total RNA was extracted using the NanoPure RNA extraction kit, ensuring efficient recovery from low cell numbers. The Arcturus™ RiboAmp™ PLUS RNA Amplification Kit (Thermo Fisher Scientific) was used for linear amplification of low-input RNA from as little as 1–10 cells, enabling molecular analysis from previously challenging quantities. This system amplified RNA from as little as 1 pg of total RNA, producing antisense RNA (aRNA) suitable for digital droplet PCR. To ensure specificity and accuracy of the amplification process, both positive and negative controls were included: positive controls (PC) consisted of RNA from PCa cell lines (LNCaP, PC3 and VCaP) known to express target genes such as AR and negative controls included RNA from healthy donor PBMCs that do not express tumor-specific genes, along with no-reverse transcription (no RT) and water controls. The amplified RNA samples were processed by iCura Diagnostics using the QIAcuity digital PCR system. This approach enabled precise quantification of target transcripts, ensuring accurate representation of the original low-abundance RNA derived from CTCs. The use of digital PCR enhanced both sensitivity and specificity, validating the amplification process and confirming the presence of tumor-specific targets.

The information for ddPCR primers and probes was summarized in Supplementary Table S3. The type III TMPRSS2-ERG fusion was detected using the published information.^92^

### Statistical analyses

Statistical analyses were conducted using GraphPad Prism v10, R, and OriginPro. All results are based on at least two to three independent experiments with biological replicates. For each replication experiment, a new aliquot of cells from the original stock was thawed and cultured. Data normality was assessed using the Shapiro–Wilk test (α = 0.05). If data were normally distributed, parametric tests were used, including unpaired *t*-tests, one-sample *t*-tests (to compare a sample mean against a defined value), and one-way or repeated-measures ANOVA (for multiple group comparisons depending on experimental design). For non-normally distributed data, non-parametric tests such as the Mann–Whitney U test or Kruskal–Wallis test were applied. Multiple group comparisons were performed using one-way ANOVA followed by Tukey’s or Bonferroni’s post hoc test, as appropriate, to adjust for multiple hypothesis testing. Specific statistical tests and sample sizes (*n*) are detailed in the figures and/or figure legends, where *n* refers to the number of animals or samples for in vivo and in vitro experiments, respectively. Data are presented as mean ± SEM or SD, along with individual data points, as specified. Statistically significant differences (*p<* 0.05) are indicated in the graphs; non-significant results are labeled “ns” or left unmarked. *For correlation analysis*, Pearson’s correlation coefficient (r) was used to assess the linear relationship between AR and BCL-2 expression levels. To evaluate the significance of this correlation, a two-tailed *t-*test was performed using OriginPro software.

*For quantification data plots*, box plots summarizing the relative % of 4 PCa cell subtypes in WM images analyzed in benign tissues, primary tumors and CRPC. Each dot in the boxplots represents a CK+ ROI (Region of Interest). P values were calculated via repeated measure two-way ANOVA with Bonferroni multiple comparison test. *p*<0.1; \**p*<0.05; \*\**p*< 0.01; \*\*\**p*< 0.001; \*\*\*\**p*<0.0001. All data analysis was done using OriginPro software.

*For analysis of in vitro and ex-vivo drug treatment data*, at least 3 independent repeats were performed in each experiment. Differences between sample groups were evaluated using Mann-Whitney U-test or Kruskal-Wallis H-test. P-value of less than 0.05 was considered statistically significant. Significance levels are indicated as follows: \**p*<0.05, \*\**p*<0.01, and \*\*\**p*<0.001, respectively. Half-maximal inhibitory concentration (IC_50_) values were calculated using nonlinear regression algorithm in Prism 7 software (GraphPad Software, USA). IC₅₀ values were estimated from relative viable cell counts following treatment with varying drug concentrations. Dose–response curves were generated in GraphPad Prism using a four-parameter logistic model, with response values normalized to the vehicle-treated control (set to 1). IC₅₀ values were calculated from the fitted curves. To compare IC₅₀ values between single-agent and combination treatments across independent experiments, a two-tailed Mann–Whitney U test was performed.

*To determine potential synergy between two treatment conditions,* experiments were performed using incremental dosing of each drug for four weeks. For combination studies, the tumor volume data was annotated as viability data and were loaded into the free software Compusyn and Synergy Finder Plus. Compusyn software implanted Chou-Talalay method to generate dose-effect curves. Combination index below 1 was considered synergistic effect. The SynergyFinder software generated surface response plots by comparing the tumor volume inhibition data to a drug combination reference model obtained from the effect of each drug alone. SynergyFinder implemented a bootstrapping method to compare viability data. We considered a P-value of less than 0.05 to establish statistical differences between drugs combinations. *For statistical re-analysis of the therapeutic data*^20^ *at the individual tumor levels* (Supplementary Fig. S6c-f), tumor volumes were measured weekly using calipers, and volume was calculated using the formula: (length × width²)/2. Individual tumor growth kinetics were plotted over time, and mean tumor volumes ± SEM were computed at each time point. The magenta arrow denotes treatment start point. To assess differences in tumor growth kinetics between groups, log-transformed tumor volumes (log₁₀ mm³) were plotted. A linear mixed model (LMM) was employed to analyze the log10-transformed growth curves, accounting for within-subject correlations. Model parameters were estimated using restricted maximum likelihood (REML). Group differences were assessed by testing the main effect of treatment, as no significant time-by-group interaction was detected following transformation. Statistical significance was determined using two-sided *t*-tests with Satterthwaite’s approximation for degrees of freedom. A p-value less than 0.05 was considered statistically significant.

*For statistical analyses of Omics data,* gene-level comparisons (e.g., *BCL2, AR*) were performed using two-tailed Student’s *t*-test. When raw counts were available, differential expression analysis was performed using DESeq2, with fold change (FC), p-values, adjusted p-values (FDR), and 95% confidence intervals (CIs) reported (e.g., Fig. 1c, e; Supplementary Table S5). For datasets without raw counts, gene expression values (TPM) were analyzed using paired Wilcoxon signed-rank tests for group paired comparison. Pearson’s correlation was used to assess linear associations between gene expression and chromatin features. The Jonckheere–Terpstra (J-T) trend tests were used to assess monotonic relationships between ordinal variables (e.g., Gleason score) and continuous values (e.g., *BCL-2* mRNA levels), using the DescTools R package (v0.99.60). All statistical analyses were conducted using R (v4.3.3) and GraphPad Prism (v10.4.1). Related figures were generated using GraphPad Prism or R as appropriate.

## DATA AVAILABILITY STATEMENT

All experimental data are available upon request from the (co-)corresponding authors. New experimental models (e.g., castration-resistant PCa cell sublines and xenograft models) will be made available upon the publication of the manuscript. All datasets analyzed in this study, summarized in Supplementary Table S4, are either publicly available from repositories or available, upon request, from the (co-)corresponding authors. The present project does not involve proprietary custom codes.

## Acknowledgements

Work in Tang lab was supported, in part, by grants from the U.S. National Institutes of Health (NIH) National Cancer Institute (NCI) R01CA240290 and 2R01CA240290-06A1, grants from the U.S. Department of Defense (DOD) PC220137 and PC220273, Roswell Park Comprehensive Cancer Center and the NCI Center grant P30CA016056, Roswell Park Alliance Foundation (RPAF) and the George Decker Endowment. Work in the labs of Tang and Chatta was further supported by the Prostate Cancer Foundation (PCF) Challenge Award 2022CHAL3788. We thank A. Witkiewicz and W. Luo (Roswell Park) for expert technical support with the Vectra and IMC experiments, respectively; D. Schapiro (Heidelberg University Hospital) in IMC data analysis using the HistoCAT software; J. Alumkal (University of Michigan) for clarifying details regarding the Alumkal dataset used in this study; J. Wang and E. Cortes (Roswell Park) for their valuable suggestions on microarray and RNA-seq data processing; the Roswell Park Public Safety team for their escort services and dedicated on-site support; and all other Tang lab members for helpful discussions and suggestions. We apologize to the colleagues whose work was not cited due to space constraint.

## Author contributions

**A. Jamroze** led the conceptualization, experimental design and execution, data analysis & interpretation, figures preparation and manuscript writing. **X. Liu** analyzed RNA-seq and public datasets, performed secondary analysis of the Lu lab-generated AR ChIP-seq and ATAC-seq data, prepared related figures, and contributed to data interpretation and manuscript writing. **S. Hou and H. Yu** provided statistical support; **H. Yu** also contributed to manuscript editing and feedback. **W. (Jess)** and **A. Tracz** provided technical support with animal studies execution**. J. Kirk** designed and conducted some therapeutic animal studies and contributed to data discussion, and critical review of the manuscript. **J. Huang** provided tissue microarray (TMA) samples and pathology support**. X. Chen and Q. Li** contributed some Western blot and therapeutic data and participated in manuscript review. **Y. Lu and K. Lin** supported *in silico* AR ChIP-seq and ATAC-seq data analyses. **K. Nastiuk** provided the Roswell Park nADT RNA-seq data and editorial input and critical feedback**. I. Puzanov** and **G. Chatta** provided clinical insight and critical comments on the manuscript. **D.G. Tang** was responsible for the overall development, execution and accomplishment of the project, including project concept, experimental strategies, data acquisition, interpretation and presentation, funding acquisition and manuscript finalization. D.G. Tang serves as the lead corresponding author.

**Supplementary information** accompanies the manuscript on the Signal Transduction and Targeted Therapy website http://www.nature.com/sigtrans

---

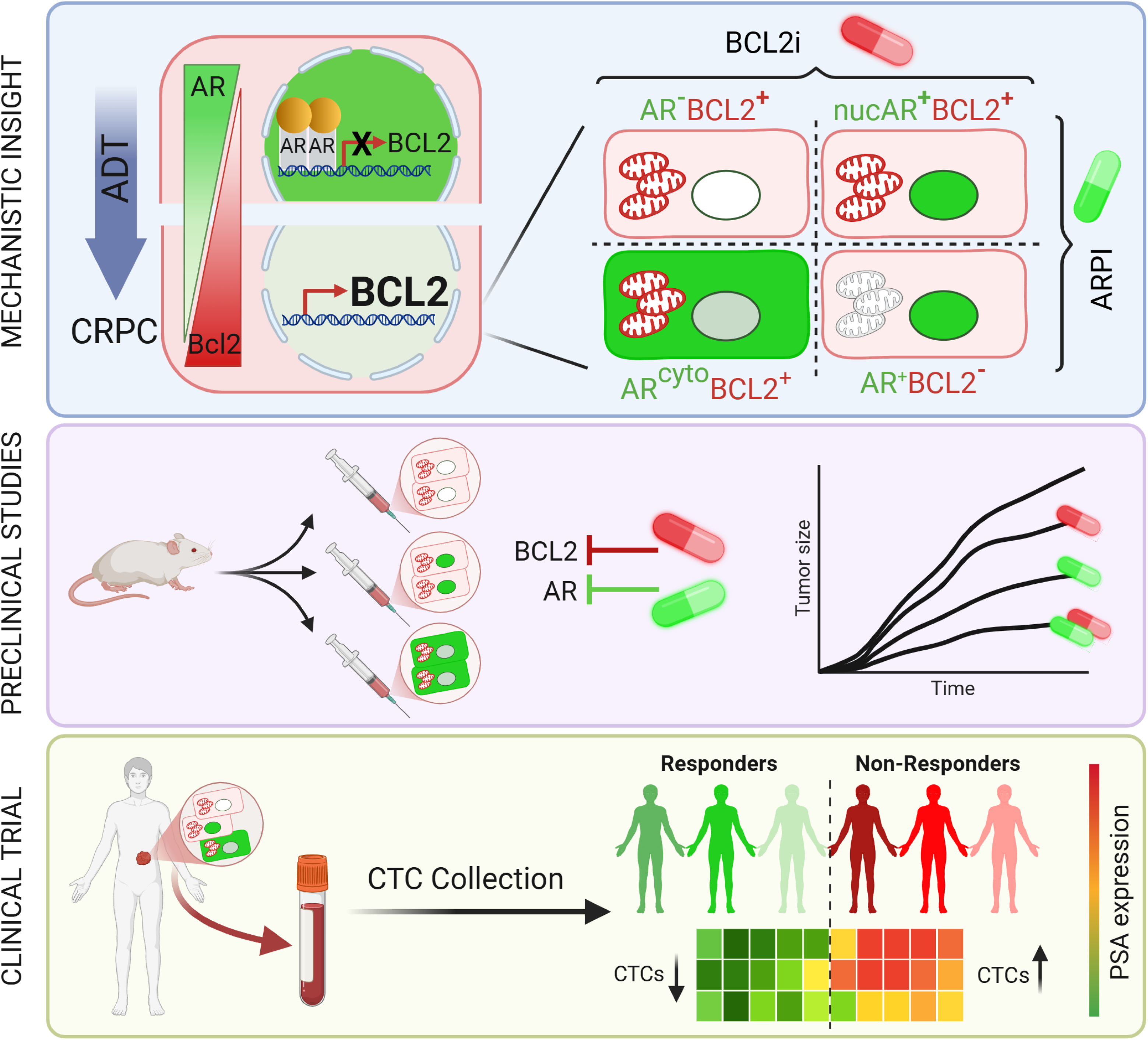

