## Supplementary material for "Cell-resolved high-dimensional imaging analysis, therapeutic modeling, and a Phase Ib clinical study establish BCL-2 as a target across heterogeneous CRPC subtypes": Table S1 - S5; Supplementary Figure S1 - S15

**Supplementary Table S1.** List of antibodies
**Supplementary Table S2.** Clinical specimens analyzed in Vectra-based quantitative multiplex immuno-fluorescence (qmIF) and imaging mass cytometry (IMC)
**Supplementary Table S3.** Chemicals and reagents employed throughout experimental procedures
**Supplementary Table S4.** Summary of ChIP-seq, RNA-seq, and ATAC-seq datasets used for profiling AR binding, transcriptional changes, and chromatin accessibility across prostate cancer models and treatment conditions
**Supplementary Table S5.** Fold changes and 95% CI for BCL-2 family genes.

**Related to Fig. 2**
**Supplementary Fig. S1.** Experimental scheme and increased $AR^{+}BCL\text{-}2^{-}$ cells in treatment-naïve primary PCa.
**Supplementary Fig. S2.** qmIF analysis of dynamic changes in $AR^{+/-}$ and/or $BCL\text{-}2^{+/-}$ cells in TMA-1.
**Supplementary Fig. S3.** Increased diversity in $AR^{+/-}BCL\text{-}2^{+/-}$ cells and increased $BCL\text{-}2^{+}$ PCa cells in CRPC.
**Supplementary Fig. S4.** Increased diversity in $AR^{+/-}BCL\text{-}2^{+/-}$ and increased $BCL\text{-}2^{+}$ PCa cells in CRPC.
**Supplementary Fig. S5.** Quantitative summary of $AR^{+/-}BCL\text{-}2^{+/-}$ cell subtypes in primary PCa and CRPC.

**Related to Fig. 3**
**Supplementary Fig. S6.** AR heterogeneity in primary CRPC linked to distinct Enza response.
**Supplementary Fig. S7.** Dynamic changes in $AR^{+/-}BCL\text{-}2^{+/-}$ PCa cell types across the LNCaP-AD, and 1° and 2° LNCaP-CRPC models.
**Supplementary Fig. S8.** Dynamic changes in $AR^{+/-}BCL\text{-}2^{+/-}$ PCa cell types in LAPC9-AD/AI xenograft models.
**Supplementary Fig. S9.** Dynamic changes in $AR^{+/-}BCL\text{-}2^{+/-}$ PCa cell types in VCaP-AD/AI xenograft models.

**Related to Fig. 4**
**Supplementary Fig. S10.** Generation and phenotypic characterization of castration-resistant LAPC4 sublines.

**Related to Fig. 5**
**Supplementary Fig. S11.** Androgen-dependent AR occupancy at the *BCL-2* Locus Is lost in CRPC.
**Supplementary Fig. S12.** The $AR^{-/lo}$ CRPC have increased chromatin accessibility surrounding the *BCL-2* genomic region.

**Related to Fig. 6**
**Supplementary Fig. S13.** The $AR^{cyto}BCL\text{-}2^{+}$ LAPC4-AI organoids are sensitive to BCL-2i ABT-199.

**Related to Fig. 7**
**Supplementary Fig. S14**. Therapeutic studies with the BCL-2i ABT-199 in 3 CRPC models.

**Related to Fig. 8**
**Supplementary Fig. S15**. CTC gene expression dynamics in responders and non-responders.

.

**Table S1. Detailed information on antibodies used**

*Abbreviations: mAb, monoclonal antibody; pAb, polyclonal antibody.

| Antibody | Source | Reactivity | Company | Catalog # | Clone | Usage | Dilution |
|---|---|---|---|---|---|---|---|
| AR | Rabbit mAb | Human | CST | 8938S | D6F11 | Vectra | 1/200 |
| BCL-2 | Mouse mAb | Human | abcam | ab692 | 100/D5 | Vectra | 1/100 |
| Cytokeratin | Mouse mAb | All species | Dako | #M3515 | AE1/AE3 | Vectra | 1/100 |
| AR-$^{156}$Gd | Rabbit | Human | Cell Signaling | 5153 | D6F11 | IMC | 1/100 |
| BCL2-$^{146}$Nd | Mouse | Human | Biolegend | 658701 | 100 | IMC | 1/100 |
| Pan-CK-$^{141}$Pr | Mouse | Human, Rat | Biolegend | 914204 | AE1/AE3 | IMC | 1/100 |
| A1/Bfl-1 | Rabbit mAb | All species | Cell signaling | #14093 | D1A1C | WB | 1/1000 |
| Apoptosis cocktail | Rabbit | Human | Abcam | ab136812 | - | WB | 1/1000 |
| AR | Rabbit mAb | Human | CST | 8938S | D6F11 | WB | 1/1000 |
| BCL-2 | Mouse mAb | Human | Cell signaling | #15071 | 124 | WB | 1/1000 |
| BCL-w | Rabbit pAb | H, M, R | LS bio | LS-C382259 | | WB | 1/1000 |
| BCL-xL | Rabbit mAb | H, M, R, Mk | Cell signaling | #2764 | 54H6 | WB | 1/1000 |
| Cleaved Caspase-3 | Rabbit | H M R Mk | Cell signaling | #9661 | - | WB | 1/1000 |
| Caspase-3 | Rabbit pAb | H M R Mk | Cell signaling | #9662 | Asp175 | WB | 1/1000 |
| β-Actin | Rabbit mAb | H,M,R,Mk,Pg | CST | 5125 | 13E5 | WB | 1/1000 |
| Mcl-1 | Rabbit mAb | Human | CST | 5453 | D35A5 | WB | 1/1000 |
| BCL2 | Mouse | Human | Biolegend | 658701 | 100 | IF | 1/150 |
| AR Alexa Fluor® 647 Conjugate | Rabbit mAb | Human | CST | #7397 | D6F11 | IF | 1/100 |
| AR | Rabbit mAb | Human | CST | 8938S | D6F11 | IHC | 1/500 |

**Supplementary table 2: Information available on patient samples in the TMAs and whole-mount sections.**

| | Source | Subject ID/# of Patients | Tissue type | Treatment |
|---|---|---|---|---|
| **Whole Mount** | **UCLA** | U12-7606-CRPC | Prostate | LHRH agonist and bicalutamide |
| | | U13-01316-CRPC | Prostate | LHRH agonist |
| | | U13-6707-CRPC | Prostate | LHRH agonist |
| | | S13-13553-CRPC | Prostate | Abiraterone and enzalutimide |
| | | S13-19900-CRPC | Prostate | LHRH agonist |
| | | HPCa14N | Prostate | No Treatment |
| | | HPCa18N | Prostate | No Treatment |
| | | HPCa21N | Prostate | No Treatment |
| | | HPCa28N | Prostate | No Treatment |
| | | HPCa27T | Prostate | No Treatment |
| | | HPCa31T | Prostate | No Treatment |
| | | HPCa33T | Prostate | No Treatment |
| **CRPC TMA** | **UCLA CRPC TMA (20 CRPC samples /40 cores)** | CRPC-1 | Prostate | LHRH agonist |
| | | CRPC-2 | Prostate | LHRH agonist |
| | | CRPC-3 | Prostate | LHRH agonist |
| | | CRPC-4 | Prostate | LHRH agonist |
| | | CRPC-5 | Prostate | LHRH agonist |
| | | CRPC-6 | Prostate | LHRH agonist |
| | | CRPC-7 | Prostate | LHRH agonist |
| | | CRPC-8 | Prostate | LHRH agonist |
| | | CRPC-9 | Prostate | LHRH agonist |
| | | CRPC-10 | Prostate | LHRH agonist |
| | | CRPC-11 | Prostate | LHRH agonist |
| | | CRPC-12 | Prostate | LHRH agonist |
| | | CRPC-13 | Prostate | LHRH agonist |
| | | CRPC-14 | Prostate | Radiation, LHRH agonist, and bicalutamide |
| | | CRPC-15 | Prostate | Radiation, LHRH agonist, and bicalutamide |
| | | CRPC-16 | Prostate | Radiation, LHRH agonist, and bicalutamide |
| | | CRPC-17 | Prostate | Radiation, LHRH agonist, and bicalutamide |
| | | CRPC-18 | Prostate | Radiation and cryotherapy |
| | | CRPC-19 | Prostate | LHRH agonist (2 weeks) |
| | | CRPC-20 | Prostate | LHRH agonist (4 mo) + bicalutamide (2 mo) |
| **TMA- GL-115** | **Jiaoti Huang** | 115 (N+T) | Prostate | No Treatment |
| **Xenograft TMA 22** | **Tang Lab** | 4 AD-AI pairs | Xenografts | Intact+Castartion |
| **Xenograft TMA 23** | **Tang Lab** | 4 AD-AI pairs | Xenografts | Intact+Castartion |

*Presented is available patient/sample information for 2 TMA sets, and 12 whole mount, N/Tand CRPC samples from UCLA. The 2 TMA sets contained a total of 125 cores derived from duplicate sections of patient samples. UCLA whole-mount sections were from the prostate (radical prostatectomy or TURP).Note that virtually all samples were collected from patients treated decades ago (mostly with LHRH agonists) before the introduction of new generation antiandrogens; only one patient (whole mount S13-13553) was treated with abiraterone and enzalutamide. De-identified patient samples were used and detailed treatment information for most patients is unavailable. *N, normal/benign glandular regions analyzed; T, tumor regions analyzed. HPCa, treatment-naïve human PCa. (Li et al., Nat Commun. 2018; Li et al., Nat Commun. 2019).

**Table S3. Comprehensive list of all reagents utilized throughout the study**

| Chremical/kit Name | Company | Catalog # |
|---|---|---|
| Enzalutamide | Selleck Chemicals | S1250 |
| Enzalutamide | Apex Bio | A3003 |
| ABT-199 | Apex Bio | A8194 |
| Sotrastaurin | Apex Bio | A8525 |
| Trypan Blue Solution | Thermo | 15250061 |
| Cultrex Poly-L-Lysin | R&D | 3438-200-01 |
| IMDM media | Gibco™ | 12440053 |
| DMEM Media | Gibco™ | 11965092 |
| IMDM no phenol red | Gibco™ | 21056023 |
| 191/193Ir DNA Intercalator | Fluidigm | 201192B |
| Paraformaldehyde | VWR | PI28908 |
| Ethylenediaminetetraacetic acid (EDTA) | Fisher Scientific | BP118500 |
| Maxpar 10x Barcode Perm Buffer | Fluidigm | 201057 |
| Puromycin | Sigma-Aldrich | P7255 |
| Mito view Green | Biotium | 70054 |
| Prolong Gold antifade with DAPI | Thermo | P36941 |
| Western Lightning Plus-ECL | Perkin Elmer | NEL104001 |
| Wes EZ standard pack 12-230 kDa | Protien Simple | PS-ST01EZ-8 |
| Wes EZ standard pack 2-40 kDa | Protien Simple | PS-ST05EZ-8 |
| Wes antibody diluent 2 | Protien Simple | 042-205 |
| Wes Anti-mouse secondary | Protien Simple | 042-205 |
| Wes Anti-rabbit secondary | Protien Simple | 042-206 |
| Wes Anti-goat secondary HRP | Protien Simple | 043-522 |
| Wes peroxide | Protien Simple | 043-379 |
| Wes streptavidin HRP | Protein Simple | 042-414 |
| Wes Luminol-S | Protien Simple | 043-311 |
| Western Lightning ECL Pro | Perkin Elmer | NEL12001 |
| Trizol reagent | Thermo | 15596018 |
| BSA | Cell Signaling | 9998 |
| BD Pharmingen™ FITC Annexin V Apoptosis Detection Kit II | BD Pharmingen | 556570 |
| LIVE/DEAD Cell Imaging Kit (488/570) | Thermo Fisher Scientific | R37601 |
| Firefly & Renilla Luciferase Single Tube Assay Kit, 150 assays | OriGene | PR300008 |
| Resazurin Assay Kit (Cell Viability) | Abcam | ab129732 |
| ChIP-IT® qPCR Analysis Kit | Active Motif | 5302 |
| SsoAdvanced Universal Probes Supermix | Bio-Rad | 1725280 |
| SuperScript™ III Reverse Transcriptase | Thermo Fisher Scientific | 18080093 |
| ChIP-IT High Sensitivity® Kit | Active Motif | 53040 |
| High Sensitivity Chromatin Preparation | Active Motif | 53046 |
| Arcturus RiboAmp HS PLUS RNA Amplification Kit (24 samples) | Thermo Fisher Scientific | KIT0505 |
| PicoPure RNA Isolation Kit (40 samples) | Thermo Fisher Scientific | KIT0204 |

**Sequences and specifications of qPCR and ddPCR primers and probes used**

| Probe ID | Catalog number | Assay ID |
|---|---|---|
| PrimePCR™ Probe Assay: MCL1, Human | 12001950 | qHsaCEP0052441 |
| PrimePCR™ Probe Assay: BCL2L1, Human | 12001950 | qHsaCEP0039517 |
| PrimePCR™ Probe Assay: BCL2, Human | 12001950 | qHsaCEP0058350 |
| PrimePCR™ Probe Assay: TBP, Human | 10031228 | qHsaCIP0036255 |
| PrimePCR™ Probe Assay: BCL2L2, Human | 12001950 | qHsaCEP0025467 |
| PrimePCR™ Probe Assay: HPRT1, Human | 10031228 | qHsaCIP0030549 |
| PrimePCR™ Probe Assay: AR human | 10025536 | qHsaCIP0026366 |
| PrimePCR™ Probe Assay: KLK3 human | 10025543 | qHsaCEP0051088 |
| PrimePCR™ Probe Assay:TMPRSS2 human | 10025544 | qHsaCEP0051087 |
| PrimePCR™ Probe Assay:BCL2L1, human | 10025545 | qHsaCEP0039517 |
| PrimePCR™ Probe Assay:NR3C1, human | 10025546 | qHsaCEP0050768 |
| PrimePCR™ Probe Assay:AMACR, human | 10025547 | qHsaCIP0029215 |
| PrimePCR™ Probe Assay:EPCAM, human | 10025548 | qHsaCEP0051089 |
| PrimePCR™ Probe Assay:TBP, human | 10025549 | qHsaCIP0036255 |
| PrimePCR™ Probe Assay:HPRT1, human | 10025550 | qHsaCIP0030549 |
| PrimePCR™ Probe Assay:PTPRC, human | 10025551 | qHsaCEP0041630 |

**Chip qPCR primers/probe sequences**

| Binding site# | Part # | Sequences |
|---|---|---|
| ARBS1 | 10031276 | Forward primer: ACAAGTTGCACGTGTGTATT |
| | | Reverse primer: CCCAATAATCCAGTGTCCCT |
| | | Probe sequence: AGTAAGCCGCTGTGCTTCTAGAAG |
| ARBS2 | 10031276 | Forward primer: TGGACAAGACGGTTTGTAAGA |
| | | Reverse primer: AACAGAACGAGGTACAGATCA |
| | | Probe sequence: TGTCTCTGTGGCATCTAACAGCGTF |
| ARBS3 | 10031276 | Forward primer: CACACACACACACGAAGGAT |
| | | Reverse primer: CTAAGGTTACGAGCTGAGCC |
| | | Probe sequence: AGTTCATGAGAGACTGGCTTGCTTGAF |
| Neg control seq | 10031279 | Forward primer: GTGGCTACCTAGGACTGG |
| | | Reverse primer: TGACATCTGTTTCCAGAACTT |
| | | Probe sequence: TTCATCATTCCAAATGGAAACTCTACCCAF |

**Table S4: Summary of ChIP-seq, RNA-seq, and ATAC-seq datasets used to profile AR binding, transcriptional changes, and chromatin accessibility across prostate cancer models and treatment conditions.**

| # | Figures | Item | Histology (Sample Type) | Data Type/Data Source | PMID | Publication |
|---|---|---|---|---|---|---|
| 1 | Fig.1a | BCL2 mRNA in normal prostate | Normal Prostate | RNA-seq/PMID: 27926864 (supplementary) | 27926864 | Liu X et al., Cell Rep 2016 |
| 2 | Fig.1a | BCL2 mRNA in normal prostate | Normal Prostate | Microarray/GSE89050 | 27926864 | Liu X et al., Cell Rep 2016 |
| 3 | Fig.1b | BCL2 mRNA in TCGA PRAD | Primary PCa (Pri-PCa) | RNA-seq/Xena portal | 26544944 | Abeshouse A et al., Cell 2015;163(4):1011-1025 |
| 4 | Fig.1c | BCL2 family mRNAs in 7 matched pre-/post-nADT prostate cancer pairs from Rajan et al., 2014 | PCa (nADT) | RNA-seq/GSE48403 | 24054872 | Rajan P et al., Eur Urol. 2014;66(1):32-39 |
| 5 | Fig.1c | BCL2 family mRNAs in 20 matched pre-/post-nADT prostate cancer pairs from Sharma et al., 2018 | PCa (nADT) | RNA-seq/GSE111177 | 30314329 | Sharma NV et al., Cancers (Basel). 2018;10(10). |
| 6 | Fig.1c | BCL2 family mRNAs in post-nADT vs. matched untreated PCa (n=43 each), Roswell cohort (Nastiuk & Chatta) | PCa (nADT) | RNA-seq/Nastiuk | unpublished | Jamroze A et al. (Chatta G., Nastiuk KL) 2024 (Submitted) |
| 7 | Fig.1c | BCL2 family mRNAs in CRPC tumors post- vs. pre-enzalutamide treatment (n=21 pairs) from Alumkal et al., 2022 | mCRPC | RNA-seq/PMID: 36109521 (supplementary) | 36109521 | Westbrook TC et al., Nat commun. 2022,13(1): 5345. |
| 8 | Fig.1e | BCL2 family mRNAs in PCa cell lines LNCaP-ARKO (6) vs. AR (4) (in vitro) | PCa cell lines | RNA-seq/Tang | unpublished | unpublished |
| 9 | Fig.1e | BCL2 family mRNAs in PCa cell lines LNCaP-ARKO (7) vs. AR (7) (in vivo, castr) | PCa cell lines | RNA-seq/Tang | unpublished | unpublished |
| 10 | Fig.1e | BCL2 family mRNAs in PCa Xenografts LNCaP Pri CRPC (4) vs. AD (4) | Xenografts | RNA-seq/GSE88752 | 30190514 | Li Q et al., Nat Commun. 2018;9(1):3600 |
| 11 | Fig.1e | BCL2 family mRNAs in PCa Xenografts LNCaP Sec CRPC (4) vs. AD (4) | Xenografts | RNA-seq/GSE88752 | 30190514 | Li Q et al., Nat Commun. 2018;9(1):3600 |
| 12 | Fig.1e | BCL2 family mRNAs in PCa Xenografts LNCaP Sec CRPC (4) vs. Pri CRPC (4) | Xenografts | RNA-seq/GSE88752 | 30190514 | Li Q et al., Nat Commun. 2018;9(1):3600 |
| 13 | Fig.1e | BCL2 family mRNAs in PCa Xenografts LAPC9 CRPC (5) vs. AD (5) | Xenografts | RNA-seq/GSE88752 | 30190514 | Li Q et al., Nat Commun. 2018;9(1):3600 |
| 14 | Fig. 5a | AR activity, BCL2 mRNA and their correlation in 7 matched pre-/post-nADT prostate cancer pairs from Rajan et al., 2014 | PCa (nADT) | RNA-seq/GSE48403 | 24054872 | Rajan P et al., Eur Urol. 2014;66(1):32-39 |
| 15 | Fig. 5a | AR activity, BCL2 mRNA levels, and their correlation in 7 matched pre-/post-nADT prostate cancer samples from the responder subgroup (referred to as the 'Low Impact Group') in Sharma et al., 2018 | PCa (nADT) | RNA-seq/GSE111177 | 30314329 | Sharma NV et al., Cancers (Basel). 2018;10(10). |
| 16 | Fig. 5a | AR activity, BCL2 mRNA, and their correlation in 6 matched pre-/post-nADT prostate cancer pairs from Long et al., 2020 | PCa (nADT) | RNA-seq/GSE150368 | 32951005 | Long X et al., Cell Death Dis. 2020;11(9):779. |
| 17 | Fig. 5b | AR activity, BCL2 mRNA and their correlation in post-nADT vs. matched untreated PCa (n=43 each), Roswell cohort (Nastiuk & Chatta) | PCa (nADT) | RNA-seq/Nastiuk | unpublished | Jamroze A et al. (Chatta G., Nastiuk KL) 2024 (Submitted) |
| 18 | Fig. 5c | AR and BCL2 mRNA and their correlation in Patient CRPC (n=40) from Tang et al., 2022 | mCRPC | RNA-seq/GSE199190 | 35617398 | Tang F et al. Science, 2022, 376(6596): eabe1505. |
| 19 | Fig. 5g-h | AR ChIP-seq in PCa cells LNCaP-AD (GSM699631) and LNCaP-AI (GSM699630) | PCa cell lines | AR ChIP-seq/GSE28264 | 22083957 | Tan PY et al., Mol Cell Biol. 2012;32(2):399-414 |
| 20 | Fig. S5a | BCL2 mRNA in Pri-PCa and tumor adjacent benign prostate (n=52) from TCGA_PRAD | Pri-PCa and adjacent benign prostate | RNA-seq/Xena portal | 26544944 | Abeshouse A et al., Cell 2015;163(4):1011-1025 |
| 21 | Fig. S5b | BCL2 mRNA in Pri-PCa (n=131) and tumor adjacent benign prostate (n=29) from Taylor et al., 2010 | Pri-PCa and adjacent benign prostate | Microarray/GSE21034; cBioPortal | 20579941 | Taylor BS et al., Cancer Cell 2010;18(1):11-22. |
| 22 | Fig. S12a | AR activity, BCL2 mRNA and their correlation in PCa Xenografts LNCaP Sec CRPC AI (4) vs. AD (4) | Xenografts | RNA-seq/GSE88752 | 30190514 | Li Q et al., Nat Commun 2018; 9(1):3600 |
| 23 | Fig. S12b | AR ChIP-seq in BCL2 region from PCa cell line LNCaP | PCa cell line | AR ChIP-seq/GSE85558 | 29153843 | Shukla S et al. Cancer Cell 2017; 32(6):792-806.e7. |
| 24 | Fig. S12b | AR ChIP-seq in BCL2 region from PCa cell line LNCaP-abl | PCa cell line | AR ChIP-seq/GSE80238 | 35031563 | Liao et al., PNAS 2022 |
| 25 | Fig. S12b | AR ChIP-seq in BCL2 region from PCa cell line LNCaP-BicR | PCa cell line | AR ChIP-seq/GSE66037 | 26404510 | Takayama K et al. Nat Commun 2015 |
| 26 | Fig. S12b | AR ChIP-seq in BCL2 region from PCa cell line C4-2 | PCa cell line | AR ChIP-seq/GSE65066 | 27068475 | Zhao Y et al. Cell Rep 2016 |
| 27 | Fig. S12b | AR ChIP-seq in BCL2 region from PCa cell line C4-2B | PCa cell line | AR ChIP-seq/GSE72714 | 27019329 | Wang J et al. Nat Med 2016; 22(5):488-96. |
| 28 | Fig. S12c-h | AR ChIP-seq from primary PCa tissue or PDX derived from CRPC Patients from Baca et al. 2022, with LNCaP-AD/AI data (adapted from Figure 5g) shown above for alignment. | Pri-PCa and CRPC | AR ChIP-seq/GSE130408 | 36071171 | Baca SC et al. Nat Genet 2022; 54(9):1364-1375. |
| 29 | Fig. S13a-d | BCL2 ATAC-seq, BCL2 RNA-seq and their correlation in Patient CRPC (n=40: ATAC_CRPC_AR (4); ATAC_CRPC_SCL (2); ATAC_CRPC_Wnt (4); ATAC_CRPC_NE (2)) from Tang et al., 2022 | mCRPC | ATAC-seq, RNA-seq/GSE199190 | 35617398 | Tang F et al. Science, 2022, 376(6596): eabe1505. |

**Table S5. Fold Change and 95% Confidence Intervals for BCL-2 Family Genes in Clinical and Preclinical Datasets (Figure 1 Panels)**

| Associated Figure | Dataset | Gene | Aliases | Ensembl ID | FC (95% CI) | p-value | False Dicovery Rate (FDR) | Note |
|---|---|---|---|---|---|---|---|---|
| Figure 1c | Clinical Dataset #1 | *BCL2* | BCL-2 | ENSG00000171791 | 2.40 (1.61, 3.58) | 1.65E-05 | 7.87E-04 | post vs pre-nADT (n=7)/Rajan |
| Figure 1c | Clinical Dataset #1 | *BCL2L1* | BCL-XL | ENSG00000171552 | 0.68 (0.54, 0.86) | 1.02E-03 | 1.53E-02 | post vs pre-nADT (n=7)/Rajan |
| Figure 1c | Clinical Dataset #1 | *MCL1* | MCL-1 | ENSG00000143384 | 1.18 (0.86, 1.61) | 3.13E-01 | 6.04E-01 | post vs pre-nADT (n=7)/Rajan |
| Figure 1c | Clinical Dataset #1 | *BCL2L2* | BCL-W | ENSG00000129473 | 0.72 (0.56, 0.92) | 9.88E-03 | 7.28E-02 | post vs pre-nADT (n=7)/Rajan |
| Figure 1c | Clinical Dataset #1 | *BCL2A1* | A1/BFL1 | ENSG00000140379 | 1.26 (0.73, 2.20) | 4.08E-01 | 6.86E-01 | post vs pre-nADT (n=7)/Rajan |
| Figure 1c | Clinical Dataset #2 | *BCL2* | BCL-2 | ENSG00000171791 | 5.13 (12.71, 2.07) | 4.17E-04 | 2.26E-03 | post vs pre-nADT (n=20)/Sharma |
| Figure 1c | Clinical Dataset #2 | *BCL2L1* | BCL-XL | ENSG00000171552 | 1.56 (3.31, 0.73) | 2.51E-01 | 3.56E-01 | post vs pre-nADT (n=20)/Sharma |
| Figure 1c | Clinical Dataset #2 | *MCL1* | MCL-1 | ENSG00000143384 | 2.72 (4.72, 1.57) | 3.65E-04 | 2.09E-03 | post vs pre-nADT (n=20)/Sharma |
| Figure 1c | Clinical Dataset #2 | *BCL2L2* | BCL-W | ENSG00000129473 | 1.15 (3.11, 0.42) | 7.84E-01 | 8.43E-01 | post vs pre-nADT (n=20)/Sharma |
| Figure 1c | Clinical Dataset #2 | *BCL2A1* | A1/BFL1 | ENSG00000140379 | 3.55 (11.98, 1.06) | 4.06E-02 | 9.24E-02 | post vs pre-nADT (n=20)/Sharma |
| Figure 1c | Clinical Dataset #3 | *BCL2* | BCL-2 | ENSG00000171791 | 1.32 (1.13, 1.55) | 7.72E-04 | 9.83E-03 | nADT vs control (n=43)/Roswell |
| Figure 1c | Clinical Dataset #3 | *BCL2L1* | BCL-XL | ENSG00000171552 | 0.81 (0.69, 0.95) | 8.95E-03 | 4.95E-02 | nADT vs control (n=43)/Roswell |
| Figure 1c | Clinical Dataset #3 | *MCL1* | MCL-1 | ENSG00000143384 | 0.80 (0.70, 0.92) | 8.77E-01 | 9.34E-01 | nADT vs control (n=43)/Roswell |
| Figure 1c | Clinical Dataset #3 | *BCL2L2* | BCL-W | ENSG00000129473 | 1.02 (0.81, 1.28) | 2.39E-03 | 2.07E-02 | nADT vs control (n=43)/Roswell |
| Figure 1c | Clinical Dataset #4 | *BCL2* | BCL-2 | ENSG00000171791 | 5.86 (-2.51, 14.23) | 3.82E-01 | NA | post vs pre-Enza (n=21)/Alumkal |
| Figure 1c | Clinical Dataset #4 | *BCL2L1* | BCL-XL | ENSG00000171552 | 1.16 (0.70, 1.62) | 2.70E-01 | NA | post vs pre-Enza (n=21)/Alumkal |
| Figure 1c | Clinical Dataset #4 | *MCL1* | MCL-1 | ENSG00000143384 | 2.02 (0.45, 3.59) | 1.00E-04 | NA | post vs pre-Enza (n=21)/Alumkal |
| Figure 1c | Clinical Dataset #4 | *BCL2L2* | BCL-W | ENSG00000129473 | 1.35 (0.72, 1.99) | 3.70E-01 | NA | post vs pre-Enza (n=21)/Alumkal |
| Figure 1e | Pre-clinical Dataset #1 | *BCL2* | BCL-2 | ENSG00000171791 | 2.05 (1.71, 2.45) | 2.71E-15 | 4.37E-14 | LNCaP-ARKO (6) vs. AR (4) (in vitro) |
| Figure 1e | Pre-clinical Dataset #1 | *BCL2L1* | BCL-XL | ENSG00000171552 | 0.70 (0.61, 0.80) | 1.21E-07 | 7.65E-07 | LNCaP-ARKO (6) vs. AR (4) (in vitro) |
| Figure 1e | Pre-clinical Dataset #1 | *MCL1* | MCL-1 | ENSG00000143384 | 0.70 (0.64, 0.77) | 3.88E-15 | 6.18E-14 | LNCaP-ARKO (6) vs. AR (4) (in vitro) |
| Figure 1e | Pre-clinical Dataset #1 | *BCL2L2* | BCL-W | ENSG00000129473 | 0.91 (0.78, 1.05) | 2.05E-01 | 3.02E-01 | LNCaP-ARKO (6) vs. AR (4) (in vitro) |
| Figure 1e | Pre-clinical Dataset #2 | *BCL2* | BCL-2 | ENSG00000171791 | 3.17 (2.21, 4.56) | 3.70E-10 | 4.93E-09 | LNCaP-ARKO (7) vs. AR (7) (in vivo, castration) |
| Figure 1e | Pre-clinical Dataset #2 | *BCL2L1* | BCL-XL | ENSG00000171552 | 0.90 (0.75, 1.08) | 2.67E-01 | 3.78E-01 | LNCaP-ARKO (7) vs. AR (7) (in vivo, castration) |
| Figure 1e | Pre-clinical Dataset #2 | *MCL1* | MCL-1 | ENSG00000143384 | 0.57 (0.49, 0.66) | 7.72E-14 | 1.78E-12 | LNCaP-ARKO (7) vs. AR (7) (in vivo, castration) |
| Figure 1e | Pre-clinical Dataset #2 | *BCL2L2* | BCL-W | ENSG00000129473 | 1.21 (1.05, 1.41) | 1.08E-02 | 2.66E-02 | LNCaP-ARKO (7) vs. AR (7) (in vivo, castration) |
| Figure 1e | Pre-clinical Dataset #3 | *BCL2* | BCL-2 | ENSG00000171791 | 1.78 (1.16, 2.74) | 8.83E-03 | 4.87E-02 | LNCaP Pri CRPC (4) vs. AD (4) |
| Figure 1e | Pre-clinical Dataset #3 | *BCL2L1* | BCL-XL | ENSG00000171552 | 0.82 (0.65, 1.03) | 9.24E-02 | 2.68E-01 | LNCaP Pri CRPC (4) vs. AD (4) |
| Figure 1e | Pre-clinical Dataset #3 | *MCL1* | MCL-1 | ENSG00000143384 | 0.87 (0.69, 1.10) | 2.41E-01 | 4.89E-01 | LNCaP Pri CRPC (4) vs. AD (4) |
| Figure 1e | Pre-clinical Dataset #3 | *BCL2L2* | BCL-W | ENSG00000129473 | 1.13 (0.88, 1.46) | 3.39E-01 | 5.94E-01 | LNCaP Pri CRPC (4) vs. AD (4) |
| Figure 1e | Pre-clinical Dataset #4 | *BCL2* | BCL-2 | ENSG00000171791 | 3.83 (2.42, 6.05) | 9.49E-09 | 1.99E-07 | LNCaP Sec CRPC (4) vs. AD (4) |
| Figure 1e | Pre-clinical Dataset #4 | *BCL2L1* | BCL-XL | ENSG00000171552 | 0.83 (0.66, 1.05) | 1.21E-01 | 2.92E-01 | LNCaP Sec CRPC (4) vs. AD (4) |
| Figure 1e | Pre-clinical Dataset #4 | *MCL1* | MCL-1 | ENSG00000143384 | 0.77 (0.60, 0.98) | 3.08E-02 | 1.07E-01 | LNCaP Sec CRPC (4) vs. AD (4) |
| Figure 1e | Pre-clinical Dataset #4 | *BCL2L2* | BCL-W | ENSG00000129473 | 0.99 (0.76, 1.28) | 9.20E-01 | 9.77E-01 | LNCaP Sec CRPC (4) vs. AD (4) |
| Figure 1e | Pre-clinical Dataset #5 | *BCL2* | BCL-2 | ENSG00000171791 | 2.15 (1.36, 3.39) | 1.02E-03 | 4.22E-02 | LNCaP Sec CRPC (4) vs. Pri CRPC (4) |
| Figure 1e | Pre-clinical Dataset #5 | *BCL2L1* | BCL-XL | ENSG00000171552 | 1.02 (0.80, 1.28) | 8.99E-01 | 1.00E+00 | LNCaP Sec CRPC (4) vs. Pri CRPC (4) |
| Figure 1e | Pre-clinical Dataset #5 | *MCL1* | MCL-1 | ENSG00000143384 | 0.88 (0.69, 1.13) | 3.19E-01 | 8.57E-01 | LNCaP Sec CRPC (4) vs. Pri CRPC (4) |
| Figure 1e | Pre-clinical Dataset #5 | *BCL2L2* | BCL-W | ENSG00000129473 | 0.87 (0.67, 1.13) | 2.91E-01 | 8.38E-01 | LNCaP Sec CRPC (4) vs. Pri CRPC (4) |
| Figure 1e | Pre-clinical Dataset #6 | *BCL2* | BCL-2 | ENSG00000171791 | 6.38 (2.38, 17.08) | 2.27E-04 | 9.19E-04 | LAPC9 CRPC (5) vs. AD (5) |
| Figure 1e | Pre-clinical Dataset #6 | *BCL2L1* | BCL-XL | ENSG00000171552 | 0.80 (0.69, 0.93) | 4.34E-03 | 1.28E-02 | LAPC9 CRPC (5) vs. AD (5) |
| Figure 1e | Pre-clinical Dataset #6 | *MCL1* | MCL-1 | ENSG00000143384 | 0.88 (0.76, 1.03) | 1.16E-01 | 2.09E-01 | LAPC9 CRPC (5) vs. AD (5) |
| Figure 1e | Pre-clinical Dataset #6 | *BCL2L2* | BCL-W | ENSG00000129473 | 1.34 (1.15, 1.57) | 2.71E-04 | 1.07E-03 | LAPC9 CRPC (5) vs. AD (5) |

***Note:** RNA-seq expression values (TPM, FPKM, or DESeq2-normalized counts) were used as provided in the original sources.
Where applicable, fold-change (FC), p-values, and false discovery rates (FDR) were extracted from the original datasets.
No additional normalization or batch correction was performed unless otherwise specified.
FC and 95% confidence intervals (CI) were calculated using DESeq2 when raw counts were available.
For the Alumkal cohort, where raw counts were unavailable, FC and statistical comparisons were calculated from TPM values using paired one-side Wilcoxon signed-rank tests.
NA: DESeq2 analysis not performed due to lack of raw counts in the original study.

**Supplementary Figure legend:**

**Figure S1. Experimental scheme and increased $AR^+BCL\text{-}2^-$ cells in treatment-naïve primary PCa.**

**a.** Patient samples and experimental models used in qmIF and IMC analyses.

**b.** Tabulated summary of Specimen types and numbers used in the current study.

**c.** Representative qmIF images from whole-mount (WM) benign prostate (HPCa14N and HPCa21N) sections stained for AR, BCL-2, and cytokeratin (CK) using the Vectra Polaris platform. Top panels show low-magnification overviews (1.5x) for full tissue context. Bottom panels show 40x magnification, including individual AR and BCL-2 images and composite views illustrating their spatial relationship. Dotted squares in 40x overview indicate regions further magnified at 80x. Note the reciprocal expression patterns of AR (localized to luminal epithelial cells) and BCL-2 (localized to basal cells) proteins.

**d.** Representative qmIF images from WM primary PCa (HPCa18 and HPCa33) sections stained for AR, BCL-2, and CK using the Vectra Polaris platform. Top panels show low-magnification (1.5x) overviews of the entire tissue sections. Bottom panels display higher magnification views, including individual and composite images of AR and BCL-2. Note that primary tumors exhibit a marked increase in $AR^+BCL\text{-}2^-$ cells when compared to benign prostate tissue,.

**Figure S2. qmIF analysis of dynamic changes in $AR^{+/-}$ and/or $BCL\text{-}2^{+/-}$ cells in TMA-1.**

**a.** Benign prostatic glands have $AR^+$ and $BCL\text{-}2^+$ cells in the luminal and basal layers, respectively. Shown on top are low-magnification (5.9x) WM images of 3 benign tissues and below are zoom-in (40x) images of 2 representative areas stained for individual markers. Dotted boxes indicate areas further magnified in the in-sets (80x) demonstrating $AR^+$ luminal and $BCL\text{-}2^+$ basal cell localization.

**b.** Primary tumors are characterized by a dramatically expanded $AR^+BCL\text{-}2^-$ PCa cell population. Shown on top are WM images of 3 tumors and at the bottom are zoom-in (40X) images of 2 representative areas stained for individual markers. Dotted boxes denote areas further magnified (80x). Note most PCa cells are $AR^+BCL\text{-}2^-$.

**c-d.** CRPC are characterized by significantly increased $(AR^{+/-})BCL\text{-}2^+$ PCa cells. Shown on top are WM images of 2 CRPC (5.9x) each and at the bottom are zoom-in (40x) images of representative areas stained for individual markers. Dotted boxes indicate areas shown in 80x in-sets. Note markedly increased $BCL\text{-}2^+$ PCa cells.

**Figure S3. Increased diversity in $AR^{+/-}BCL\text{-}2^{+/-}$ cells and increased $BCL\text{-}2^+$ PCa cells in CRPC.**

**a.** Representative qmIF images from a WM CRPC specimen (CRPC-7606), Shown are low-magnification (0.4x) composite images of the entire tissue section, displaying AR/BCL-2 (left) and AR/BCL-2/CK (right).

**b.** Two distinct regions of interest (ROIs) imaged at 40x, highlighting the double-positive $AR^{+}BCL\text{-}2^{+}$ CRPC cells. Dotted boxes indicate areas selected for further high-resolution imaging at 160x, shown in the insets.

c. Three spatially distinct ROIs from CRPC-7606 imaged at 40x, each highlighting heterogeneous AR and BCL-2 expression. Insets show further magnified views at 160x, demonstrating distinct cell subtypes including $AR^{+}BCL\text{-}2^{-}$, $AR^{-}BCL\text{-}2^{+}$, $AR^{+}BCL\text{-}2^{+}$ (double-positive), and $AR^{lo}BCL\text{-}2^{+}$ cells.

**Figure S4. Increased diversity in $AR^{+/-}BCL\text{-}2^{+/-}$ and increased $BCL\text{-}2^{+}$ PCa cells in CRPC.**

Analysis of $AR^{+/-}BCL\text{-}2^{+/-}$ PCa cell subtypes in CRPC WM-6707 (**a-b**) and CRPC WM-1316 (**c-d**). For each CRPC WM, shown on top are AR/BCL-2 (left) and AR/BCL-2/CK (right) WM images and below are high-magnification images of 3 ROIs illustrating the indicated subtypes of CRPC cells.

**Figure S5. Quantitative summary of $AR^{+/-}BCL\text{-}2^{+/-}$ cell subtypes in primary PCa and CRPC.**

**a.** *BCL-2* mRNA levels were reduced in primary PCa (Pri-PCa) in the TCGA_PRAD database. Patient numbers (n) are indicated in parentheses. Three comparisons are shown: matched pairs of prostate tumor (T) and normal (N; tumor-adjacent benign) tissues (n = 52 pairs; ****$p$<0.0001, Paired Student's *t*-test); N vs. Pri-PCa (treatment-naïve tumors in TCGA-PRAD, excluding samples from patients who received adjuvant [post-surgery] hormone therapy, n = 52 vs. 422; ****$p$<0.0001, Student's *t*-test); and N vs. all PRAD tumors (n = 52 vs. 495; ****$p$<0.0001, Student's *t*-test). Expression is shown as $\log_2$ (FPKM + 1).

**b.** *BCL-2* mRNA levels were reduced in Pri-PCa in the Taylor dataset. Patient numbers (n) are indicated in parentheses. *** $p$<0.001 (Student's *t*-test).

**c.** Quantification of the percentage of $AR^{+}$ (left) and $BCL\text{-}2^{+}$ (right) cells among epithelial cells ($CK^{+}$) in benign prostate tissue, untreated primary tumors, and CRPC specimens. Each dot represents an individual region of interest (ROI). Bars represent mean ± SD. *P* values were determined using the Wilcoxon rank-sum test and are shown above each comparison.

**d.** Proportion of PCa cell subtypes ($AR^{+}BCL\text{-}2^{-}$, $AR^{-}BCL\text{-}2^{+}$, $AR^{+}BCL\text{-}2^{+}$, $AR^{-}BCL\text{-}2^{-}$) within $CK^{+}$ regions across benign, primary tumor, and WM CRPC tissues. Group means were compared using one-way ANOVA with Bonferroni correction.

**e.** Mean cell density (cells/mm²) of each AR/BCL-2-defined cell subtype within $CK^{+}$ compartments from whole-mount tissues. Statistical comparisons were performed using one-way ANOVA with Bonferroni adjustment. Each point represents a single ROI.

f. Cell density analysis of $AR^{+/-}BCL\text{-}2^{+/-}$ PCa cell subtypes pooled across all $CK^{+}$ specimens, including both whole-mount and TMA samples. Data are shown as box plots with means compared by one-way ANOVA and Bonferroni post hoc testing. *p* values reflect significant pairwise differences.

For d-f, each dot in the box plots represents one $CK^{+}$ ROI, and *P* values for individual comparisons are indicated (repeated measures two-way ANOVA with Bonferroni multiple comparison test).

**Figure S6. AR heterogeneity in primary CRPC linked to distinct Enza response.**

**a.** Experimental scheme of generating, from androgen-dependent (AD) xenograft tumors, androgen-independent (AI), castration-resistant primary (1°) CRPC and castration/Enza-resistant secondary (2°) CRPC.

**b.** AR IHC images highlighting 3 AR expression/localization patterns in the AI xenograft models, i.e., nuclear $AR^{+/hi}$ (LNCaP-AI), $AR^{-/lo}$ (LAPC9-AI) and $AR^{cyto}$ (LAPC4-AI and VCaP-AI).

**c-f.** Distinct Enza responses in the four 1° CRPC (AI) models. Left panels: Shown are individual tumor volume measurement (red, control (CTL) mice; blue, Enza-treated mice. Animal numbers for each group are indicated in parentheses on the right), with downward arrows indicating the starting time of Enza treatment (i.e., at 1, 2.5, 4 and 5 weeks, respectively, for LNCaP, LAPC9, LAPC4 and VCaP 1° CRPC). Middle panels: Shown are the mean tumor volumes of the CTL (red) and Enza treatment (blue) groups. *$p<0.05$ at the time points compared between the two groups (paired Student's *t*-test). Right panels: Shown are differences in tumor growth kinetics between groups using log-transformed tumor volumes. Group differences were assessed by testing the main effect of treatment, and statistical significance was determined using two-sided *t*-tests with Satterthwaite's approximation for degrees of freedom (see Methods).

**Figure S7. Dynamic changes in $AR^{+/-}BCL\text{-}2^{+/-}$ PCa cell types across the LNCaP-AD, and 1° and 2° LNCaP-CRPC models.**

**a.** Schematic workflow for IMC analysis.

**b.** qmIF images of LNCaP-AD tumors stained for AR, BCL-2, and CK, with top panels showing full-tissue views (1.5x) and bottom panels showing enlarged (40x) images highlighting $AR^{+}BCL2^{-}$ cells.

**c.** Primary LNCaP-AI (1° CRPC) tumors showed markedly increased $AR^{+/hi}$ cells and slightly increased $BCL\text{-}2^{+}$ PCa cells. (top: 1.5x, bottom: 40x).

**d.** Secondary castration-/Enza-resistant LNCaP-AI (2° CRPC) tumors displayed an $AR^{+}BCL\text{-}2^{+}$ phenotype in most tumor cells. (top: 1.5x, bottom: 40x).

**e.** Representative IMC images of LNCaP-AD and LNCaP-AI (1° CRPC) tumors stained for AR, BCL-2, and DNA. Shown on the right are representative zoom-in images of AR and BCL-2 in the AD/AI tumors.

For b-d: the top panels show merged, zoomed-out views along with corresponding individual marker channels (AR, BCL-2, and DNA shown in distinct colors). Bottom panels display higher-resolution images highlighting spatial reorganization of marker expression patterns.

**Figure S8. Dynamic changes in $AR^{+/-}BCL\text{-}2^{+/-}$ PCa cell subtypes in LAPC9-AD/AI xenograft models.**

**a.** qmIF images of LAPC9-AD tumors stained for AR, BCL-2, and CK. Top panels show full-tissue views at 1.5x; bottom panels show 40x magnified ROIs highlighting $AR^{+}BCL\text{-}2^{-}$ cells.

**b.** LAPC9-AI (1° CRPC) tumors are populated by $AR^{-}BCL\text{-}2^{+}$ cells. Top panels show 1.5x full-tissue views; bottom panels present 40x magnified ROIs highlighting the $AR^{-}BCL\text{-}2^{+}$ cells.

**c.** Representative IMC images of LAPC9-AD and LAPC9-AI tumors stained for AR, BCL-2, and DNA. Shown below are images acquired at 100$\mu$m resolution (bottom) to validate AR/BCL-2 expression dynamics under androgen deprivation.

**Figure S9. Dynamic changes in $AR^{+/-}BCL\text{-}2^{+/-}$ PCa cell types in VCaP-AD/AI xenograft models.**

**a.** qmIF images of VCaP-AD tumors stained for AR, BCL-2, and CK. Shown on top are WM images (1.5x) of BCL-2 staining alone (left) and compound AR, BCL-2 and CK staining (right). Shown below are 40× magnified ROIs highlighting the $AR^{+}BCL\text{-}2^{+}$ VCaP-AD cells.

b. VCaP-AI (1° CRPC) tumors are populated mostly by $AR^{cyto}BCL\text{-}2^{-/lo}$ PCa cells. Shown on top are WM images of BCL-2 staining alone (left) and compound AR, BCL-2 and CK staining (right). Shown below are 40× magnified ROIs highlighting the $AR^{cyto}BCL\text{-}2^{-/lo}$ phenotype of VCaP-AI cells.

**c.** Representative IMC images of VCaP-AD and VCaP-AI tumors stained for AR, BCL-2, and DNA (top). Shown below are images acquired at 100$\mu$m resolution, illustrating changes in AR and BCL-2 protein expression and localization in response to androgen deprivation.

**Figure S10. Generation and phenotypic characterization of castration-resistant LAPC4 sublines.**

**a.** Schematic of the treatment protocol to generate resistant sublines. LAPC4-AD cells were first cultured in CDSS to create LAPC4-CR cells. Chronic treatment of LAPC4-CR cells with 20 µM or 100 µM enzalutamide (Enza) for 4 weeks produced LAPC4-Enza(20)-R and LAPC4-Enza(100)-R sublines, respectively.

**b.** Phase-contrast images (10x and 20x magnification) showing the morphology of LAPC4-AD, LAPC4-CR, LAPC4-Enza(20)-R, and LAPC4-Enza(100)-R cells. All castration-resistant LAPC4 cell sublines exhibited altered morphology compared to parental (LAPC4-AD) cells.

**c.** Cell proliferation assays showing relative cell numbers of each subline over 5 days. All castration-resistant LAPC4 sublines showed reduced growth kinetics compared to the parental LAPC4-AD cells.

**d.** Multiplex immunofluorescence images (40x) of LAPC4-AD, LAPC4-CR, and Enza-resistant sublines [LAPC4-Enza (20)-R and LAPC4-Enza(100)-R] stained for AR (Cy5, red), BCL-2 (TRITC, yellow), mitochondria (MitoTracker, green), and nuclei (DAPI, blue). Isotype (IgG) controls are shown for each condition. Merged panels display combined marker expression across cell lines.

**Figure S11. Androgen-dependent AR occupancy at the *BCL-2* Locus Is lost in CRPC.**

**a.** RNA-seq analysis (GSE88752) of LNCaP-AD/AI xenografts reveals reduced AR activity and upregulated *BCL-2* mRNA levels in LNCaP-AI tumors (top panels; *$p < 0.05$, **$p < 0.01$, Student's *t*-test). Shown below are scatter plot presenting an inverse correlation between AR activity and *BCL-2* expression (left panel; Pearson's r = –0.78, $p$ =0.023) and the AR activity vs. *BCL-2* mRNA levels in individual AD/AI tumors (right panel).

**b.** AR ChIP-seq profiles from Cistrome datasets demonstrate AR binding at the *BCL-2* locus in the indicated PCa cell lines (datasets indicated in the parentheses below).

**c–h.** Loss of AR Binding at the *BCL-2* locus in CRPC as compared to primary PCa (AR ChIP-seq data source: GSE130408). **c.** Genome browser view of AR ChIP-seq at the *BCL-2* locus in primary PCa (Pri-PCa; total n=18 with 10 representative samples shown) and mCRPC (total n=15 with 10 representative samples shown). The LNCaP-AD/AI data (adapted from Figure 5g) was shown on top for alignment. Note that strong AR binding peaks detected in LNCaP-AD (ARBS1–7, highlighted in grey, see Figure 5g) are evident in Pri-PCa but largely absent in mCRPC, mirroring the pattern in LNCaP-AI. (**d**) Visualization and (**g**) quantification of ARBS peaks at the *BCL-2* locus confirm 15 ARBSs in Pri-PCa, of which 13 exhibit significant AR binding loss in CRPC ($p$<0.05), while 2 show no significant difference (*$p < 0.05$, **$p < 0.01$, ***$p < 0.001$, **** $p < 0.0001$, Student's *t*-test.). **e**. MA plot illustrates the global loss of AR peaks in CRPC relative to Pri-PCa, including those at the *BCL-2* locus, supporting a model of AR-mediated BCL2 repression. **f.** Venn diagram showing overlap of 10 ARBSs between Pri-PCa (GSE130408) and LNCaP-AD (GSE28264). **h.** Averaged AR ChIP-seq signal across the 15 ARBSs reveals a >2-fold reduction in CRPC (paired Wilcoxon test, ****$p$<0.0001).

Data sources are indicated in the figure panels (see also Supplementary Table S4 for detailed dataset information).

**Figure S12. The AR$^{-/lo}$ CRPC have increased chromatin accessibility surrounding the *BCL-2* genomic region.**

**a.** Genome browser view of ATAC-seq tracks (GSE199190) at the *BCL-2* locus across CRPC subtypes including CRPC_AR (n=4), CRPC_SCL (n=2), CRPC_Wnt (n=4), and CRPC_NE (n=2). Note that the AR$^{-/lo}$ CRPC (CRPC_NE and CRPC_Wnt) show higher chromatin accessibility at the *BCL-2* genomic region compared to CRPC_AR and CRPC_SCL tumors.

**b.** RNA-seq tracks (GSE199190) from the same representative tumors as in (a) illustrate corresponding *BCL-2* expression levels, with higher expression in CRPC_NE and CRPC_Wnt.

**c.** Bar-coded correlation plot showing a positive relationship between *BCL-2* mRNA expression and average ATAC-seq peak intensity at the *BCL-2* locus (Pearson r = 0.66, $p$<0.0001). Color-coded distributions highlight that CRPC_NE tumors exhibit both highest chromatin accessibility and *BCL-2* expression, while

CRPC_AR tumors show the lowest in both dimensions. Color and shape keys indicating CRPC subtype and AR expression status are shown adjacent to the correlation plots.

**d**. Bar-coded correlation plot demonstrating an inverse relationship between AR activity scores and *BCL-2* mRNA levels across CRPC tumors (Pearson r = –0.28, $p$ = 0.086). CRPC_AR tumors exhibit the highest AR activity and lowest *BCL-2* expression, whereas CRPC_NE tumors display the lowest AR activity and highest *BCL-2* expression.

**Figure S13. The $AR^{cyto}BCL\text{-}2^{+}$ LAPC4-AI organoids are sensitive to BCL-2i ABT-199.**

**a.** Schematic workflow for generation of organoids from LAPC4-AD and LAPC4-AI tumors.

**b.** Representative brightfield images showing the LAPC4-AI organoid morphology at Days 3, 6, and 9 following plating at three different densities.

**c.** Bar graphs presenting the cell growth (i.e., relative fluorescence units (RFU) from resazurin assays) for LAPC4-AI organoids seeded at different densities (1,000, 2,500, and 5,000 cells/well).

**d.** Heatmap of combination index for Enza and ABT-199 treatments in LAPC4-AD organoids.

**e.** Synergy scores for Enza/RU486 combination treatments in LAPC4-AI organoids. The line graph shows a positive synergy score (CI <1) indicating synergistic interactions.

**f.** Combination index (CI) plot from combination treatment of Enza and ABT-199 in LAPC4-AD organoids. CI values plotted against the fraction affected (Fa). CI > 1 suggests antagonism.

**g.** 2D dose-response matrix depicting percentage inhibition across Enza and ABT-199 concentration gradients in LAPC4-AD organoids.

**h.** Dose-response curves of RU486 treatment in LAPC4-AD and LAPC4-AI organoids, showing relative viability as a function of RU486 concentration. $IC_{50}$ values are indicated.

**i.** Bar graphs show normalized viable cell numbers after treatment with increasing concentrations of RU486 in LAPC4-AD and LAPC4-AI organoids. Statistical significance determined by unpaired two-tailed Student's *t*-test; $^{*}p< 0.05$, $^{**}p< 0.01$.

**j.** Synergy score plots depicting the interaction between Enza and RU486 across different dose combinations in LAPC4-AI organoids. Positive synergy scores indicate synergistic effects.

**Figure S14. Therapeutic studies with the BCL-2i ABT-199 in 3 CRPC models.**

**a.** Schematic of phenotypes of LAPC4-AD and LAPC4-AI CRPC models.

**b.** Treatment schema for LAPC4-AI xenografts.

**c.** Tumor control index (TCI) score showing superior inhibition of LAPC4-AI tumors by in Enza/ABT-199 combination.

**d.** Tumor regression score showing superior response in the combination group.

**e-i**. In vitro studies in the progressive LNCaP-AD/AI models. (e). Schematic of generating LNCaP-AI progression models. (f). Immunoblot of AR and BCL-2 in parental LNCaP, LNCaP-CR, and LNCaP-C/Enza-R cells. GAPDH serves as loading control. (g). Densitometric quantification of AR and BCL-2 levels from (f), normalized to GAPDH. LNCaP-CR and LNCaP-C/Enza-R cells show reduced AR but increased BCL-2 (*$p < 0.05$). (h). Schematic of cell viability assays in LNCaP-AD/AI cell line models after ABT-199 treatment. (i). Cell viability assays showing LNCaP-AI cells were more sensitive to ABT-199 than LNCaP-AD cells (*$p < 0.05$).

**j-l**. Experiments in $AR^{-/lo}$ LAPC9-AI model. (j). Workflow for treatment with BCL-2i in LAPC9-AI CRPC xenograft model. (k). Tumor volumes at the time of randomization across treatment arms. (l). Animal body weight over time showing minimal toxicity with ABT-199 but significant toxicity with AT-101.

**Figure S15: CTC gene expression dynamics in responders and non-responders.**

**a.** One-dimensional ddPCR fluorescence plots showing longitudinal expression patterns of target genes across sequential treatment timepoints in non-responder patients, highlighting minimal baseline BCL-2 expression and compensatory NR3C1 activation at the EOT samples.

**b.** One-dimensional ddPCR fluorescence plots showing longitudinal expression patterns of target genes across sequential treatment timepoints in the 2 responder patients, illustrating BCL-2 suppression and decreased CTC burden (i.e., TMPRSS2 and TMPRSS2-ERG fusion type III).

**c.** PSA responses in responders (left) and non-responders (right). PSA levels were measured over time and plotted as percentage change from baseline (baseline set to 0%) during treatment with enzalutamide plus venetoclax. Each panel represents an individual patient, grouped by clinical response. Black dots indicate PSA measurements at serial time points. Vertical green dashed lines mark the start of combination therapy and red dashed lines indicate treatment discontinuation. The yellow-shaded region represents the off-treatment phase.

**a**

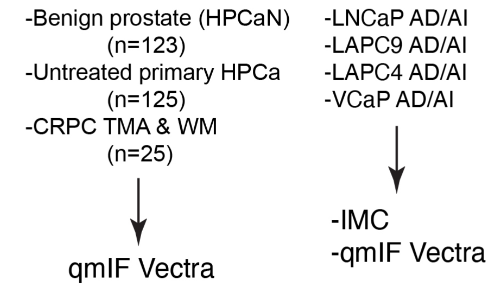


**b**

| Specimen* | Benign | Tumor | CRPC |
|---|---|---|---|
| TMA-1 | 5 | 6 | 20 |
| TMA-2 | 115 | 115 | 0 |
| WM-HPCa or WM-HPCaN | 3 benign tissues (HPCa14N,21N,28N) | 4 primary tumors (HPCa18,27,31,33) | N/A |
| WM-CRPC | N/A | N/A | 5 (#1316, 6707, 7606, 3553, 19900) |
| **Total** | **123** | **125** | **25** |

*TMA, tissue microarray (note that the TMA-2 consists of 115 paired benign and tumor pieces cut in triplicate); WM, whole-mount slices; HPCa, (primary) human PCa (HPCa); HPCaN, benign prostate tissues obtained from distant and uninvolved regions of HPCa; CRPC, castration-resistant prostate cancer.

**c**

**d**

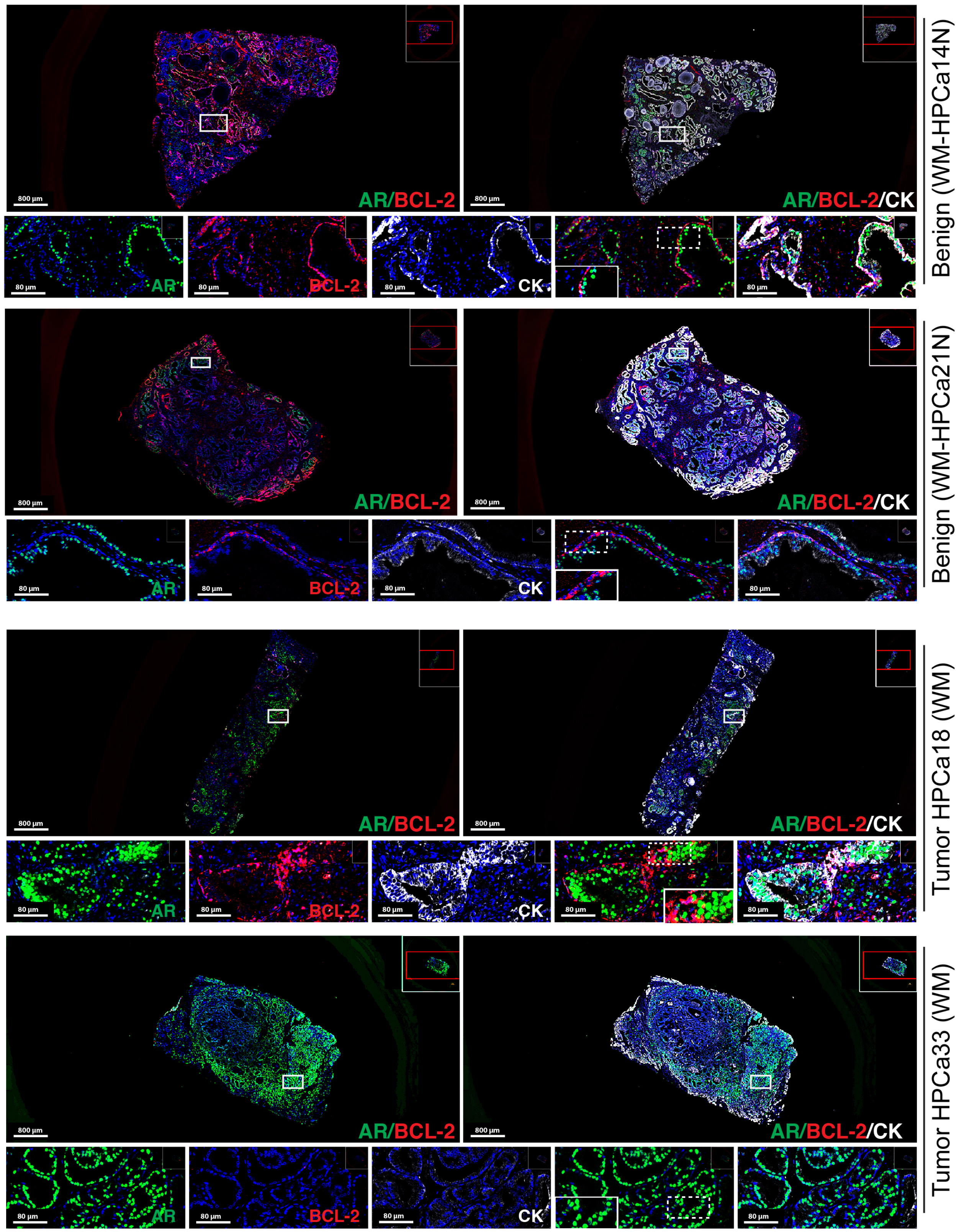


Figure S1

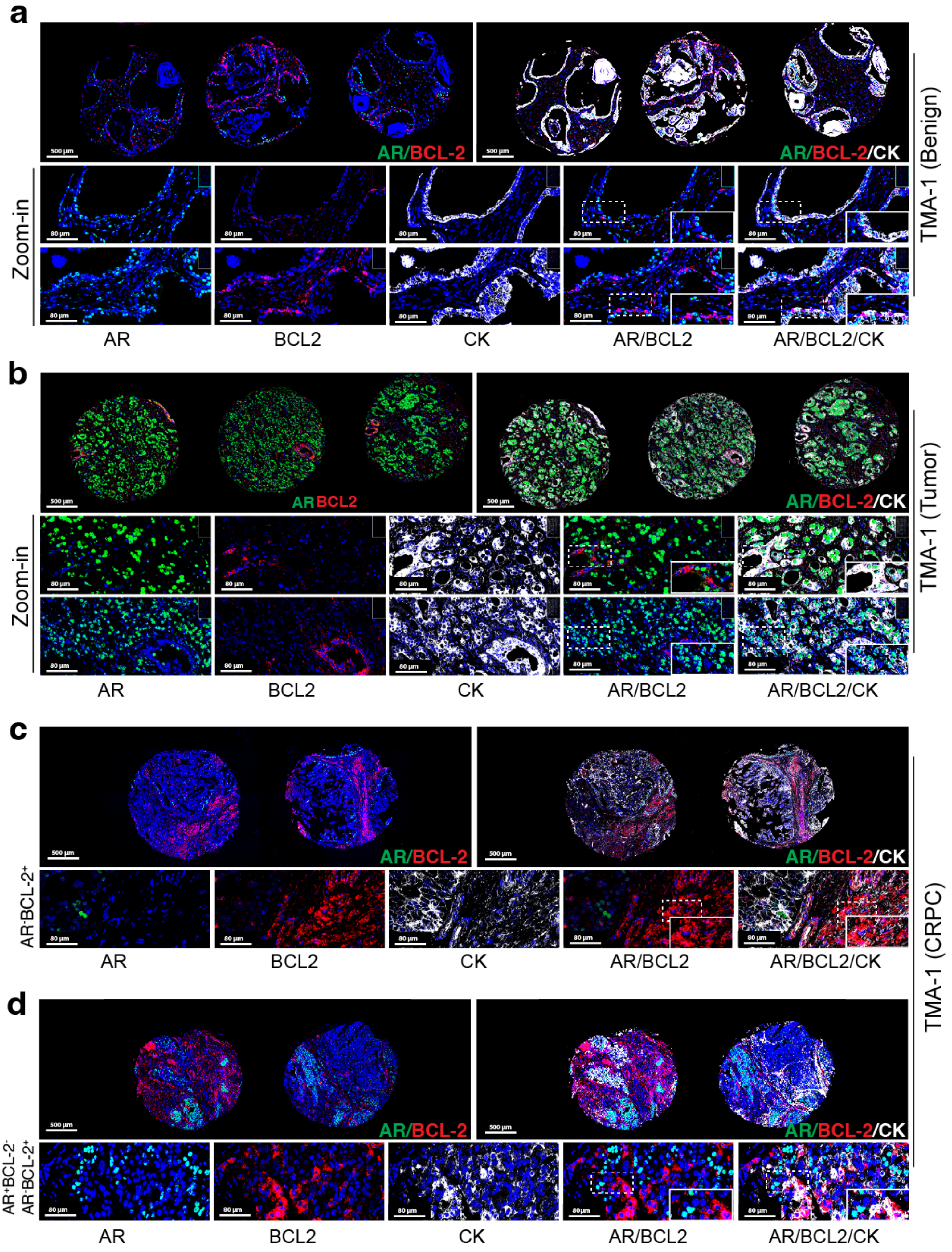
a
500 µm
AR/BCL-2
AR/BCL-2/CK
TMA-1 (Benign)
Zoom-in
80 µm
AR
BCL2
CK
AR/BCL2
AR/BCL2/CK
b
AR BCL2
AR/BCL-2/CK
TMA-1 (Tumor)
Zoom-in
AR
BCL2
CK
AR/BCL2
AR/BCL2/CK
c
AR/BCL-2
AR/BCL-2/CK
AR-BCL-2+
AR
BCL2
CK
AR/BCL2
AR/BCL2/CK
TMA-1 (CRPC)
d
AR/BCL-2
AR/BCL-2/CK
AR+BCL-2-
AR-BCL-2+
AR
BCL2
CK
AR/BCL2
AR/BCL2/CK

Figure S2

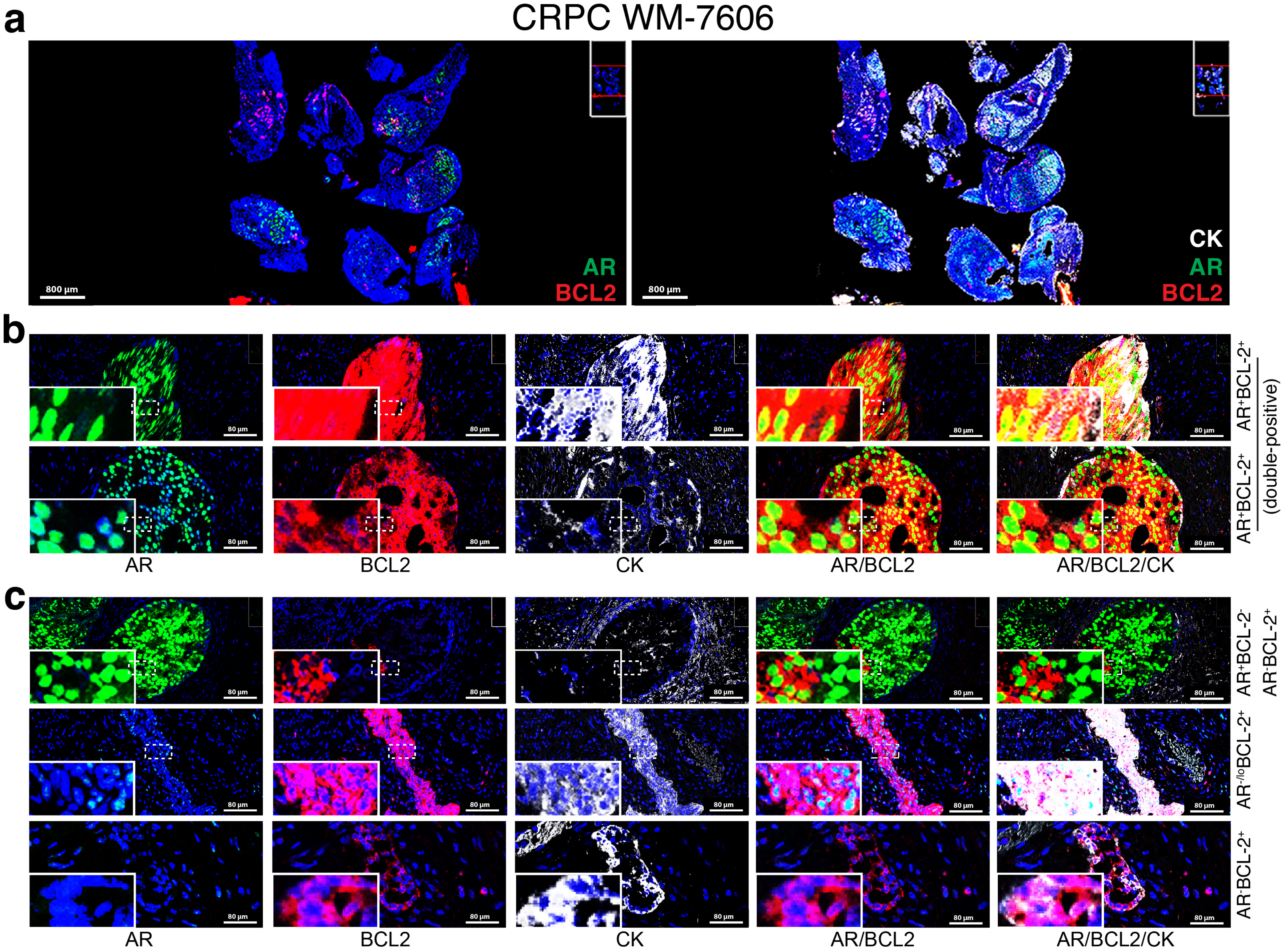
a
CRPC WM-7606
800 μm
AR
BCL2
CK
b
AR+BCL-2+
AR+BCL-2+
(double-positive)
80 μm
AR
BCL2
CK
AR/BCL2
AR/BCL2/CK
c
AR+BCL-2-
AR-BCL-2+
AR-/loBCL-2+
AR-BCL-2+

Figure S3

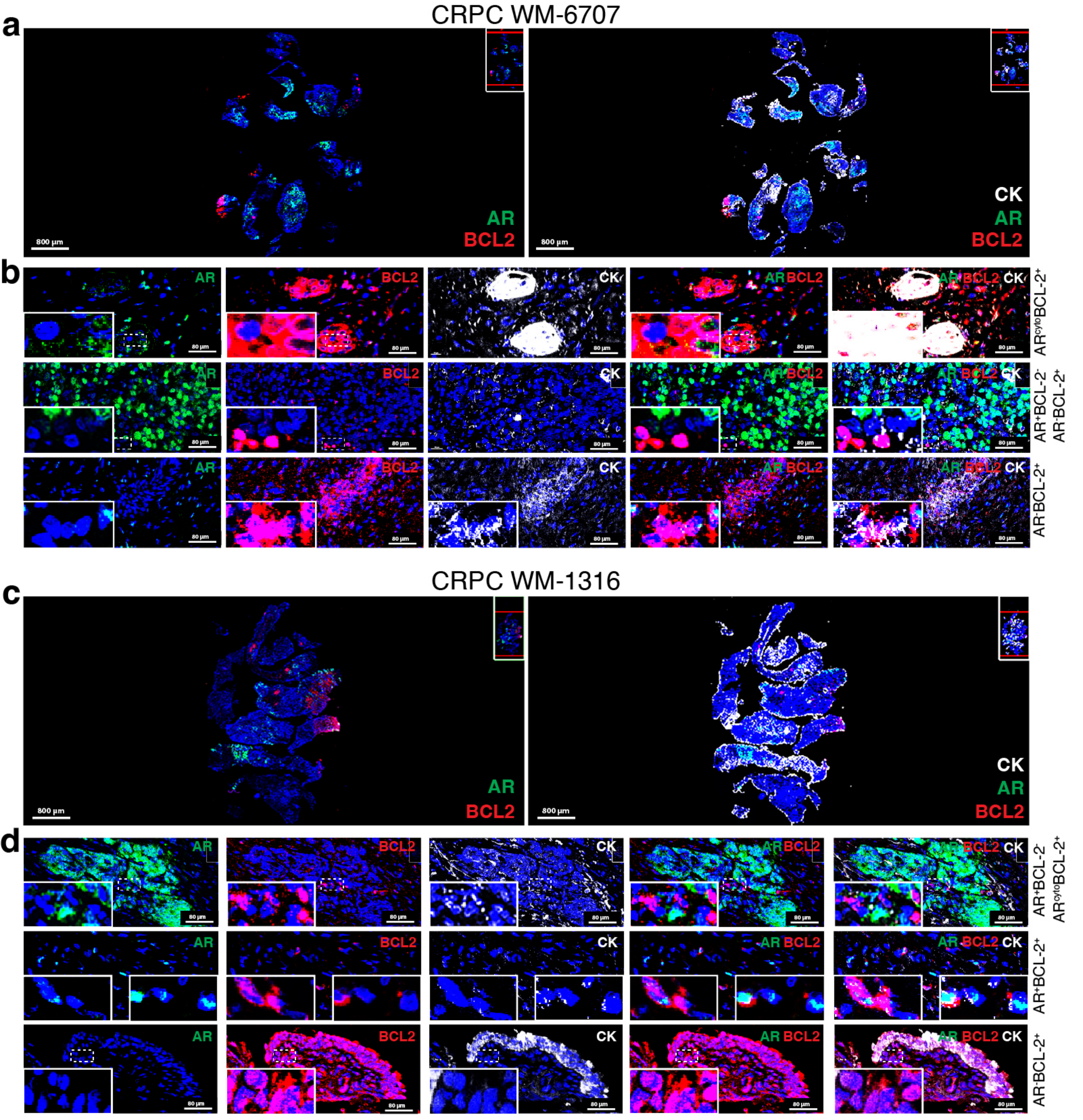

a
CRPC WM-6707
800 µm
AR
BCL2
CK
b
AR
BCL2
CK
80 µm
AR^cytoBCL-2^+
AR^+BCL-2^-
AR^-BCL-2^+
c
CRPC WM-1316
d
AR^+BCL-2^-
AR^cytoBCL-2^+
AR^+BCL-2^+
AR^-BCL-2^+


Figure S4

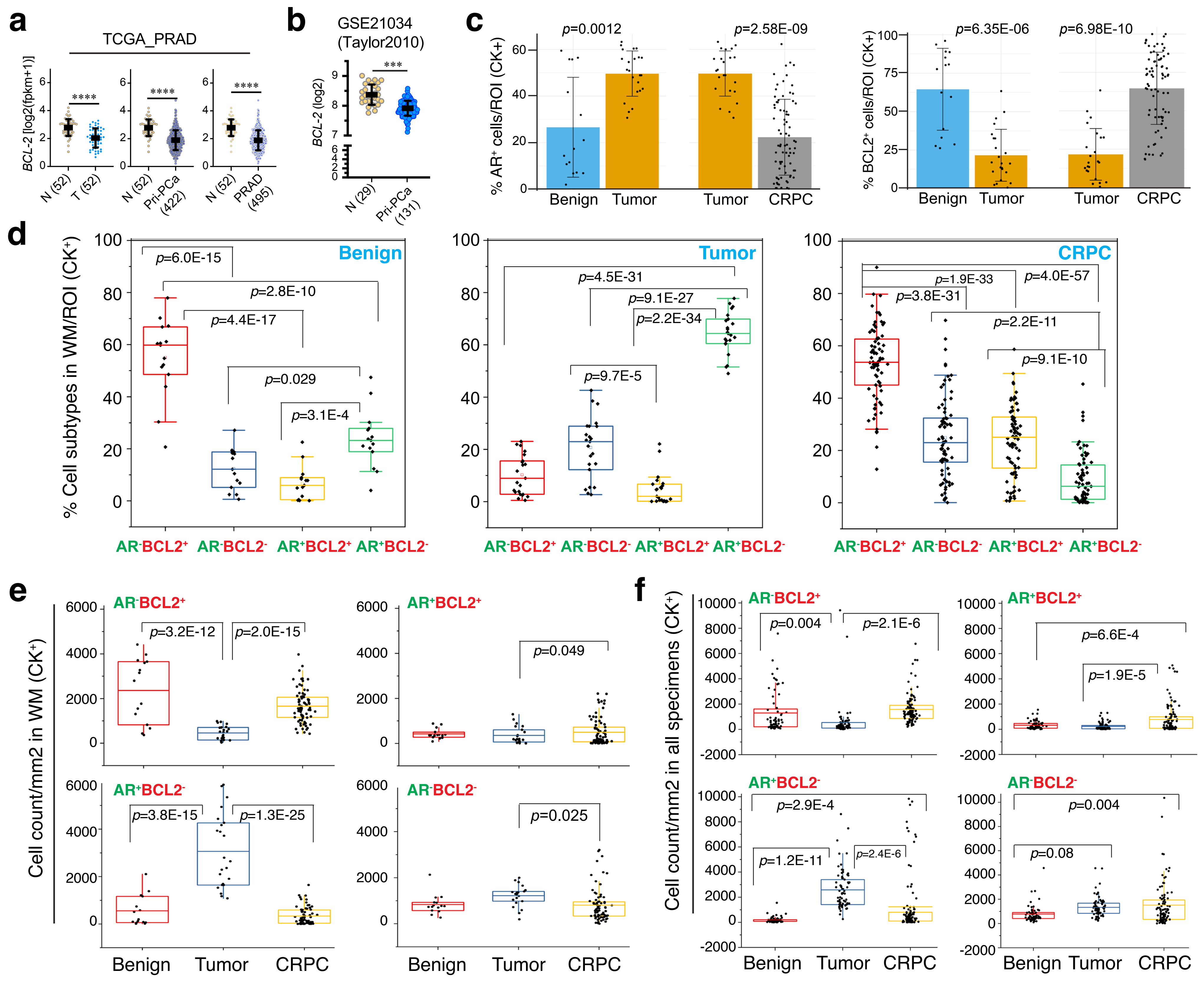

a
TCGA_PRAD
BCL-2 [log2(fpkm+1)]
N (52)
T (52)
Pri-PCa (422)
PRAD (495)
b
GSE21034 (Taylor2010)
BCL-2 (log2)
N (29)
Pri-PCa (131)
c
% AR+ cells/ROI (CK+)
p=0.0012
p=2.58E-09
Benign
Tumor
CRPC
% BCL2+ cells/ROI (CK+)
p=6.35E-06
p=6.98E-10
d
% Cell subtypes in WM/ROI (CK+)
Benign
p=6.0E-15
p=2.8E-10
p=4.4E-17
p=0.029
p=3.1E-4
Tumor
p=4.5E-31
p=9.1E-27
p=2.2E-34
p=9.7E-5
CRPC
p=1.9E-33
p=4.0E-57
p=3.8E-31
p=2.2E-11
p=9.1E-10
AR-BCL2+
AR-BCL2-
AR+BCL2+
AR+BCL2-
e
Cell count/mm2 in WM (CK+)
AR-BCL2+
p=3.2E-12
p=2.0E-15
AR+BCL2+
p=0.049
AR+BCL2-
p=3.8E-15
p=1.3E-25
AR-BCL2-
p=0.025
Benign
Tumor
CRPC
f
Cell count/mm2 in all specimens (CK+)
AR-BCL2+
p=0.004
p=2.1E-6
AR+BCL2+
p=6.6E-4
p=1.9E-5
AR+BCL2-
p=2.9E-4
p=1.2E-11
p=2.4E-6
AR-BCL2-
p=0.004
p=0.08
Benign
Tumor
CRPC


Figure S5

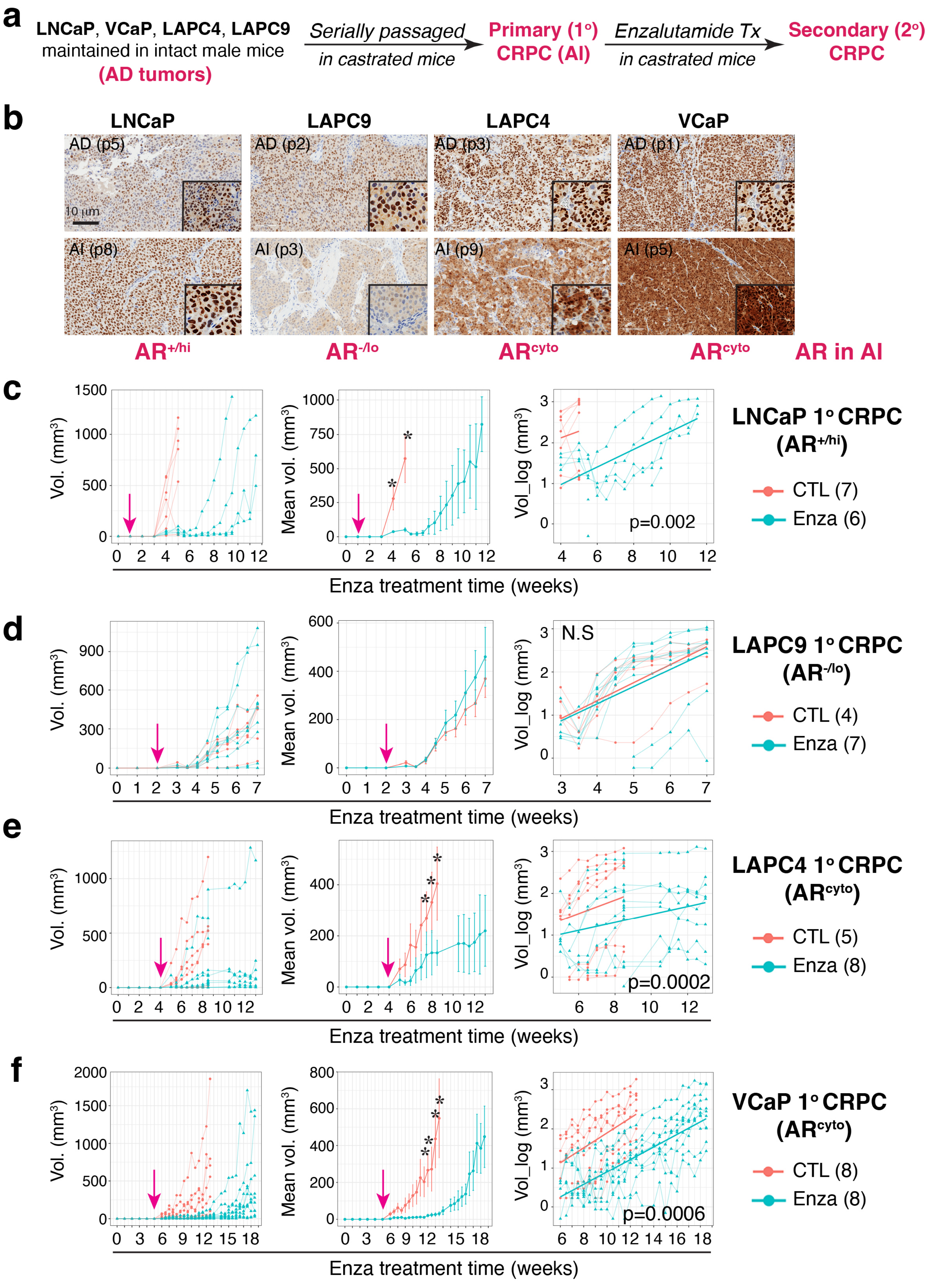
a
LNCaP, VCaP, LAPC4, LAPC9
maintained in intact male mice
(AD tumors)
Serially passaged
in castrated mice
Primary (1°)
CRPC (AI)
Enzalutamide Tx
in castrated mice
Secondary (2°)
CRPC
b
LNCaP
LAPC9
LAPC4
VCaP
AD (p5)
AD (p2)
AD (p3)
AD (p1)
10 µm
AI (p8)
AI (p3)
AI (p9)
AI (p5)
AR+/hi
AR-/lo
ARcyto
ARcyto
AR in AI
c
LNCaP 1° CRPC
(AR+/hi)
CTL (7)
Enza (6)
p=0.002
d
LAPC9 1° CRPC
(AR-/lo)
CTL (4)
Enza (7)
N.S
e
LAPC4 1° CRPC
(ARcyto)
CTL (5)
Enza (8)
p=0.0002
f
VCaP 1° CRPC
(ARcyto)
CTL (8)
Enza (8)
p=0.0006
Vol. (mm³)
Mean vol. (mm³)
Vol_log (mm³)
Enza treatment time (weeks)

Figure S6

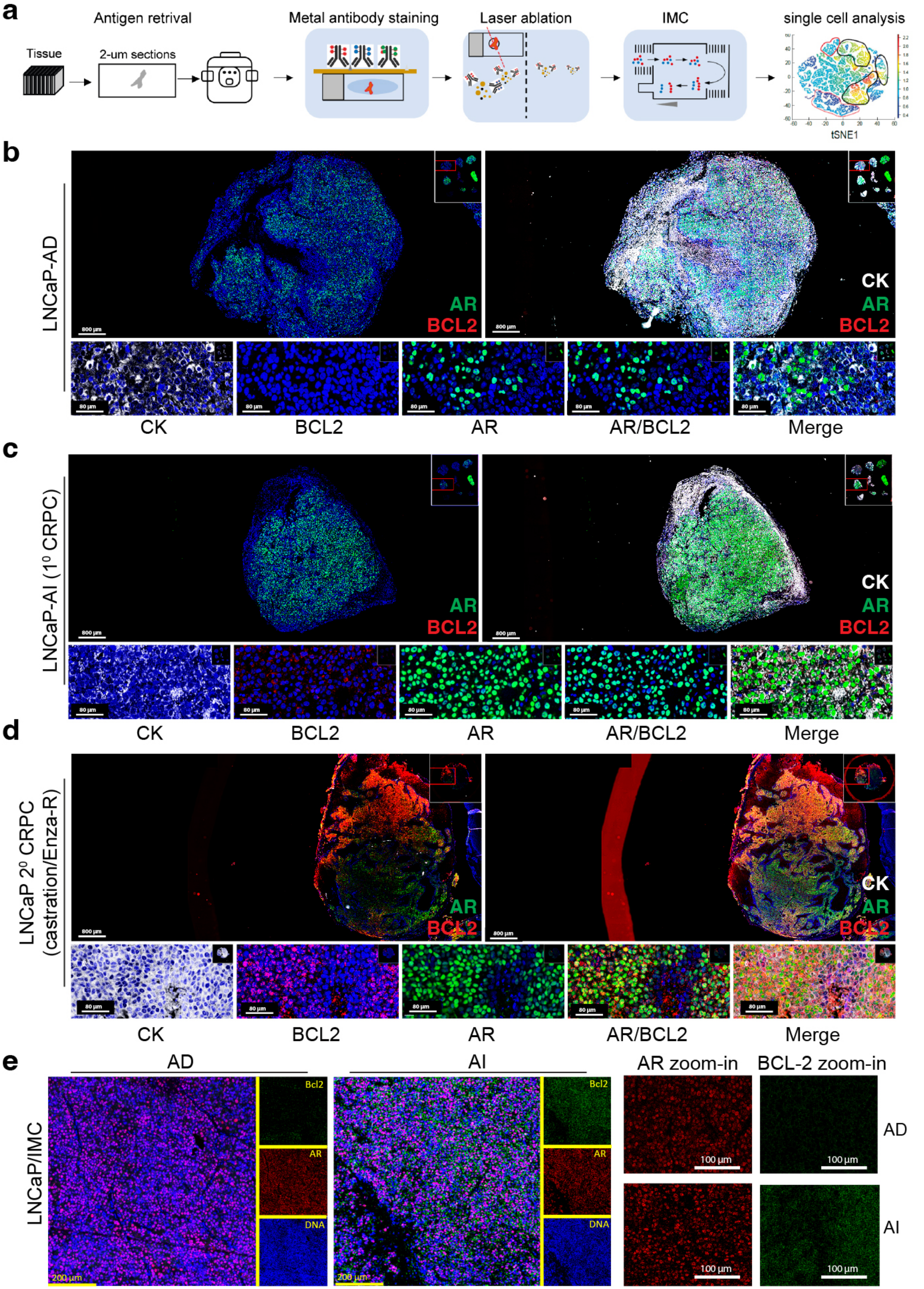
a
Antigen retrival
Metal antibody staining
Laser ablation
IMC
single cell analysis
Tissue
2-um sections
tSNE1
b
LNCaP-AD
AR
BCL2
CK
AR
BCL2
800 µm
80 µm
CK
BCL2
AR
AR/BCL2
Merge
c
LNCaP-AI (1⁰ CRPC)
AR
BCL2
CK
AR
BCL2
CK
BCL2
AR
AR/BCL2
Merge
d
LNCaP 2⁰ CRPC (castration/Enza-R)
AR
BCL2
CK
AR
BCL2
CK
BCL2
AR
AR/BCL2
Merge
e
AD
AI
AR zoom-in
BCL-2 zoom-in
LNCaP/IMC
Bcl2
AR
DNA
200 µm
100 µm
AD
AI

Figure S7

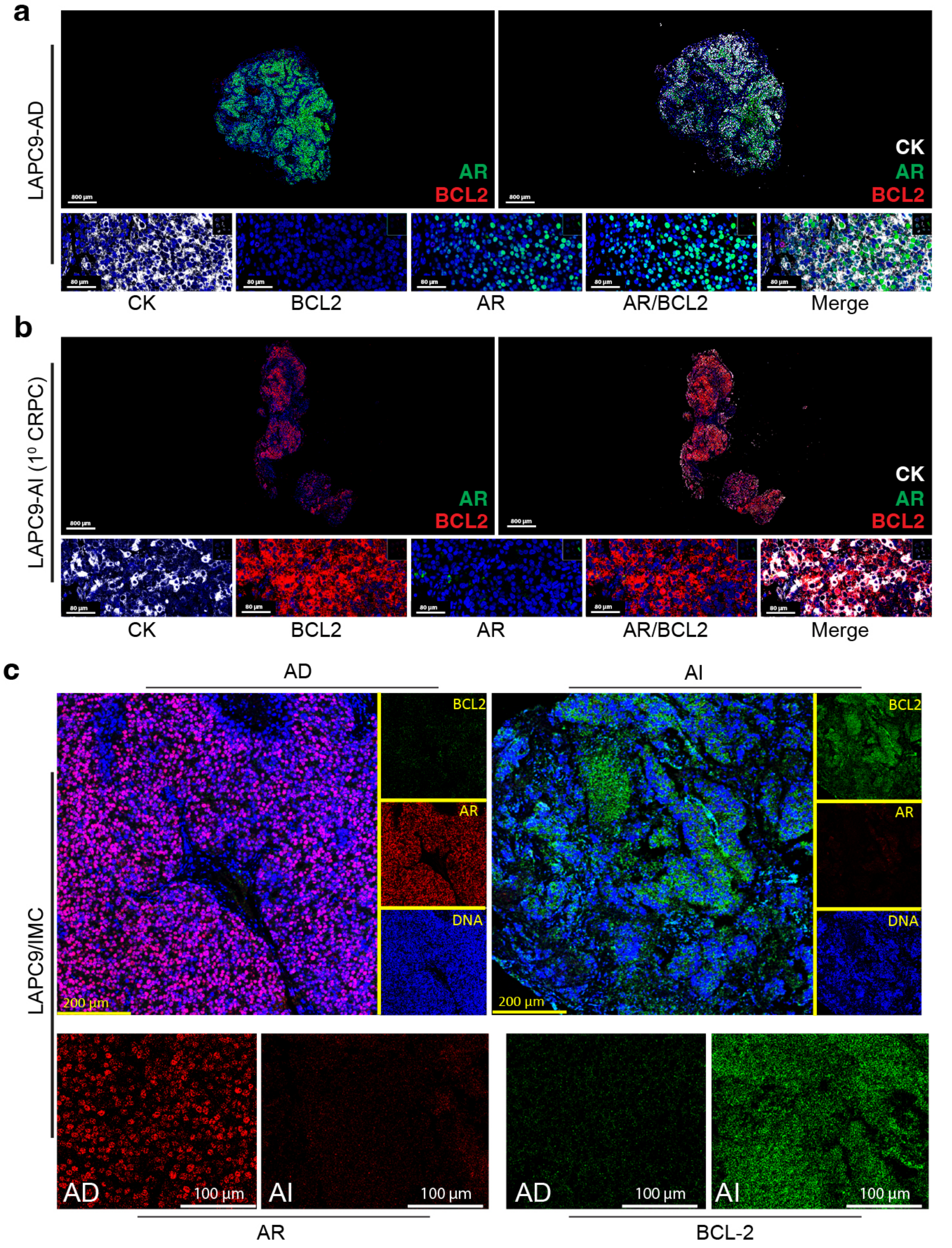
a
LAPC9-AD
AR
BCL2
CK
AR
BCL2
800 µm
80 µm
CK
BCL2
AR
AR/BCL2
Merge
b
LAPC9-AI (1° CRPC)
AR
BCL2
CK
AR
BCL2
800 µm
80 µm
CK
BCL2
AR
AR/BCL2
Merge
c
AD
AI
LAPC9/IMC
BCL2
AR
DNA
200 µm
AD
AI
100 µm
AR
BCL-2

Figure S8

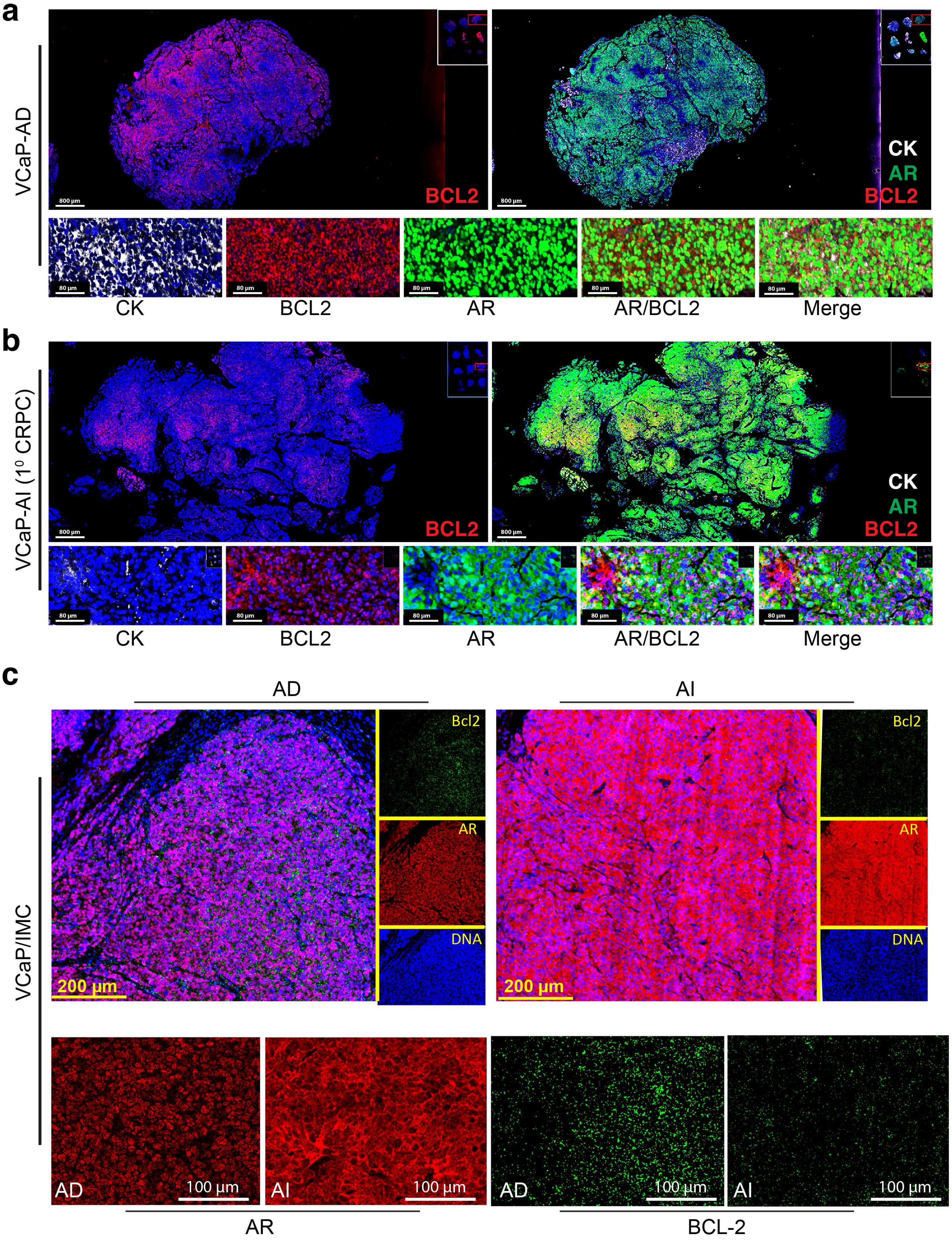
a
VCaP-AD
800 µm
BCL2
CK
AR
BCL2
80 µm
CK
BCL2
AR
AR/BCL2
Merge
b
VCaP-AI (1⁰ CRPC)
800 µm
BCL2
CK
AR
BCL2
80 µm
CK
BCL2
AR
AR/BCL2
Merge
c
AD
AI
VCaP/IMC
Bcl2
AR
DNA
200 µm
AD
AI
100 µm
AR
BCL-2

Figure S9

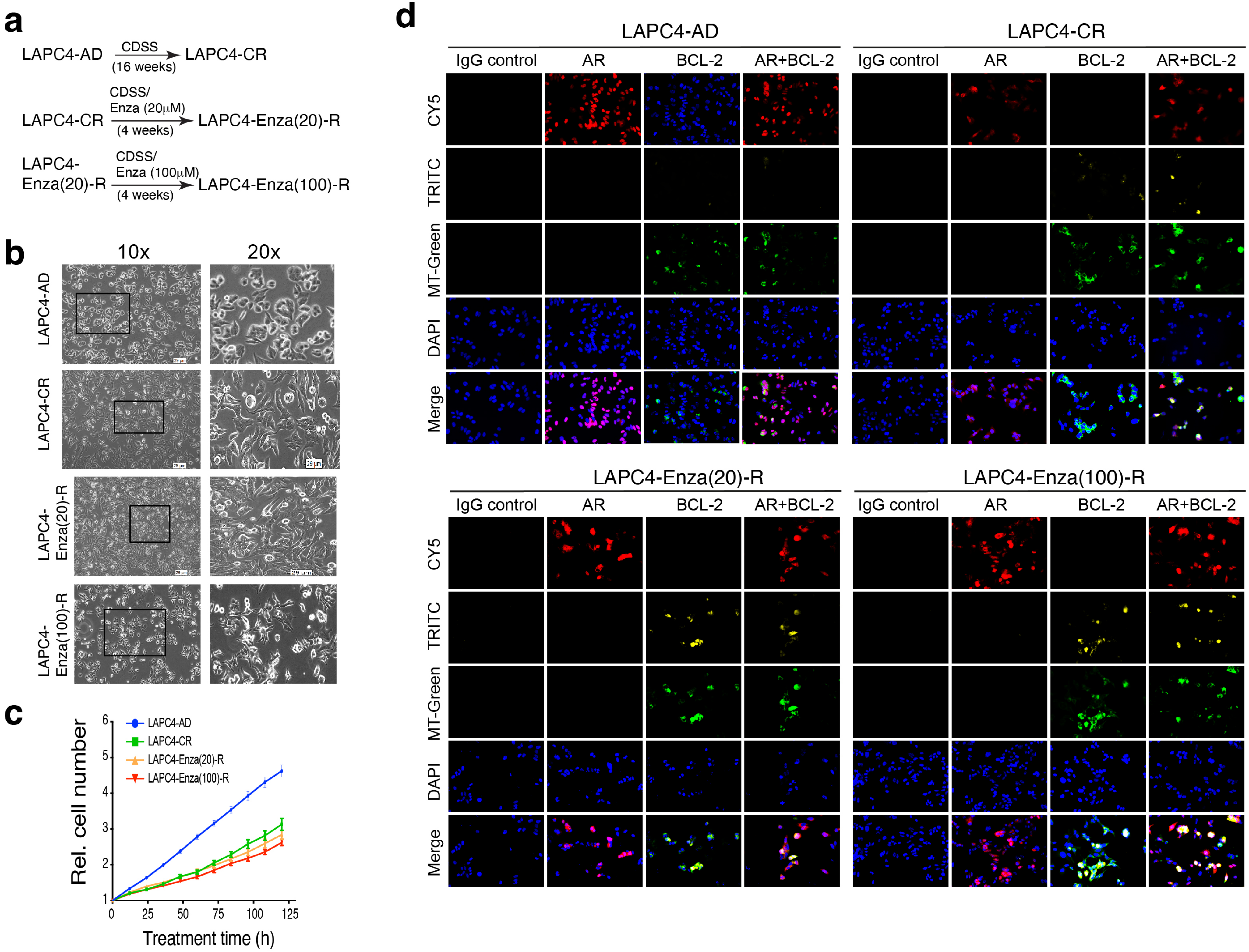
a
LAPC4-AD
CDSS
(16 weeks)
LAPC4-CR
LAPC4-CR
CDSS/
Enza (20μM)
(4 weeks)
LAPC4-Enza(20)-R
LAPC4-Enza(20)-R
CDSS/
Enza (100μM)
(4 weeks)
LAPC4-Enza(100)-R
b
10x
20x
LAPC4-AD
LAPC4-CR
LAPC4-Enza(20)-R
LAPC4-Enza(100)-R
c
Rel. cell number
Treatment time (h)
LAPC4-AD
LAPC4-CR
LAPC4-Enza(20)-R
LAPC4-Enza(100)-R
d
LAPC4-AD
LAPC4-CR
LAPC4-Enza(20)-R
LAPC4-Enza(100)-R
IgG control
AR
BCL-2
AR+BCL-2
CY5
TRITC
MT-Green
DAPI
Merge

Figure S10

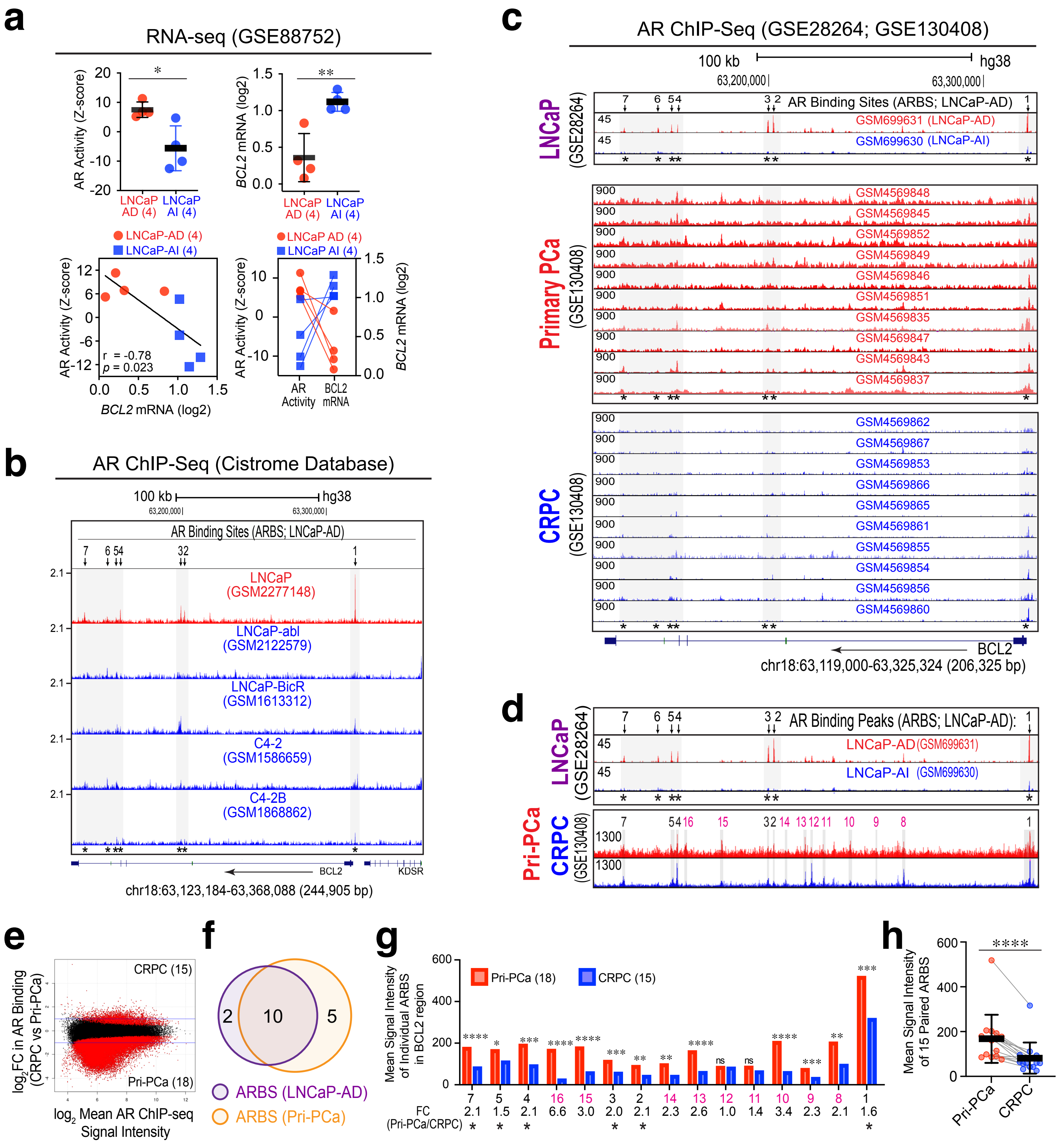

a
RNA-seq (GSE88752)
AR Activity (Z-score)
BCL2 mRNA (log2)
LNCaP AD (4)
LNCaP AI (4)
LNCaP-AD (4)
LNCaP-AI (4)
r = -0.78
p = 0.023
AR Activity
BCL2 mRNA
b
AR ChIP-Seq (Cistrome Database)
100 kb
hg38
AR Binding Sites (ARBS; LNCaP-AD)
LNCaP (GSM2277148)
LNCaP-abl (GSM2122579)
LNCaP-BicR (GSM1613312)
C4-2 (GSM1586659)
C4-2B (GSM1868862)
BCL2
KDSR
chr18:63,123,184-63,368,088 (244,905 bp)
c
AR ChIP-Seq (GSE28264; GSE130408)
LNCaP (GSE28264)
GSM699631 (LNCaP-AD)
GSM699630 (LNCaP-AI)
Primary PCa (GSE130408)
CRPC (GSE130408)
chr18:63,119,000-63,325,324 (206,325 bp)
d
AR Binding Peaks (ARBS; LNCaP-AD):
LNCaP-AD (GSM699631)
LNCaP-AI (GSM699630)
Pri-PCa
CRPC
e
log2FC in AR Binding (CRPC vs Pri-PCa)
CRPC (15)
Pri-PCa (18)
log2 Mean AR ChIP-seq Signal Intensity
f
ARBS (LNCaP-AD)
ARBS (Pri-PCa)
g
Mean Signal Intensity of Individual ARBS in BCL2 region
Pri-PCa (18)
CRPC (15)
FC (Pri-PCa/CRPC)
h
Mean Signal Intensity of 15 Paired ARBS
Pri-PCa
CRPC


Figure S11

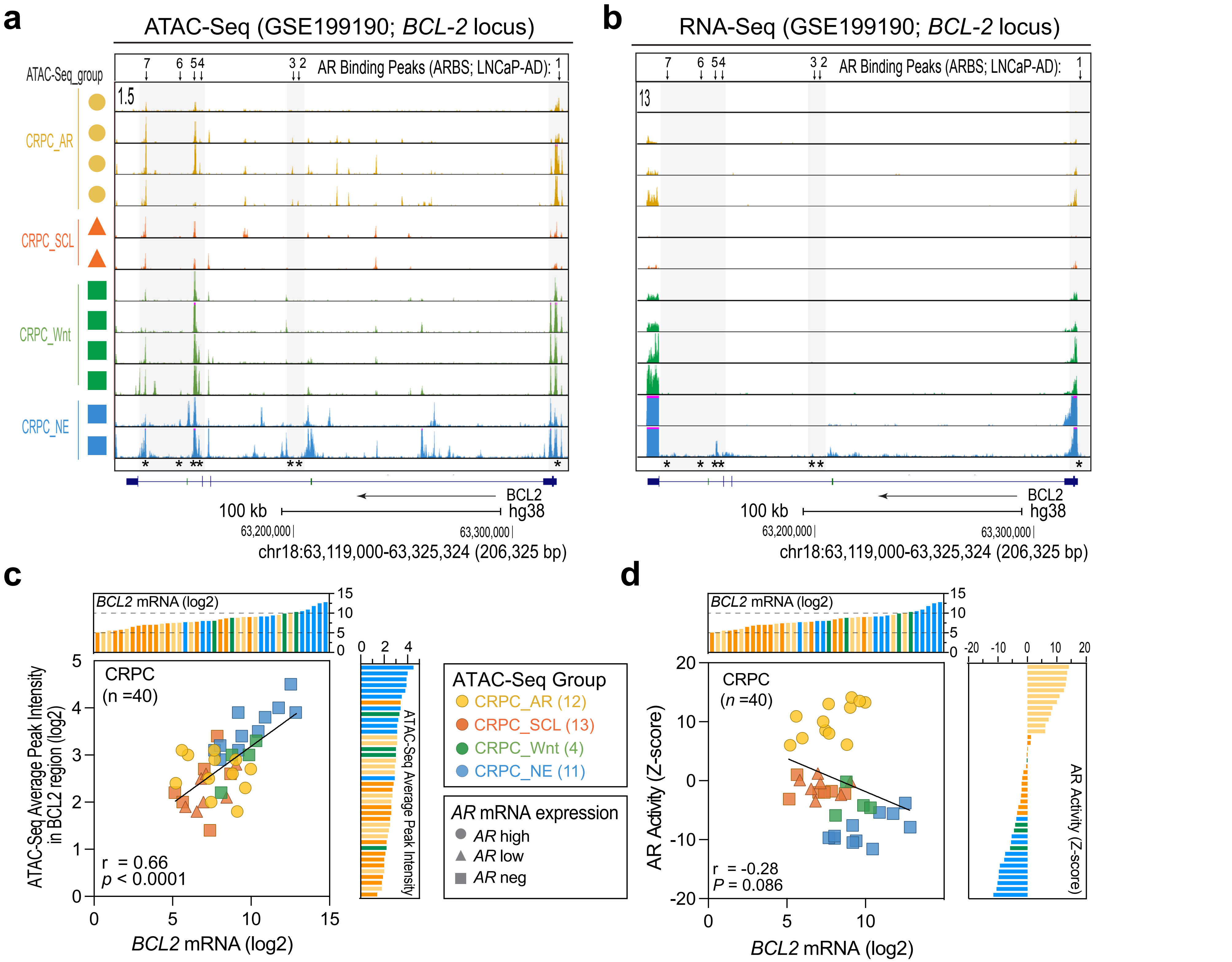
a
ATAC-Seq (GSE199190; BCL-2 locus)
AR Binding Peaks (ARBS; LNCaP-AD):
ATAC-Seq_group
CRPC_AR
CRPC_SCL
CRPC_Wnt
CRPC_NE
1.5
BCL2
100 kb
hg38
63,200,000
63,300,000
chr18:63,119,000-63,325,324 (206,325 bp)
b
RNA-Seq (GSE199190; BCL-2 locus)
AR Binding Peaks (ARBS; LNCaP-AD):
13
BCL2
100 kb
hg38
63,200,000
63,300,000
chr18:63,119,000-63,325,324 (206,325 bp)
c
BCL2 mRNA (log2)
CRPC
(n =40)
r = 0.66
p < 0.0001
ATAC-Seq Average Peak Intensity in BCL2 region (log2)
BCL2 mRNA (log2)
ATAC-Seq Average Peak Intensity
ATAC-Seq Group
CRPC_AR (12)
CRPC_SCL (13)
CRPC_Wnt (4)
CRPC_NE (11)
AR mRNA expression
AR high
AR low
AR neg
d
BCL2 mRNA (log2)
CRPC
(n =40)
r = -0.28
P = 0.086
AR Activity (Z-score)
BCL2 mRNA (log2)
AR Activity (Z-score)

Figure S12

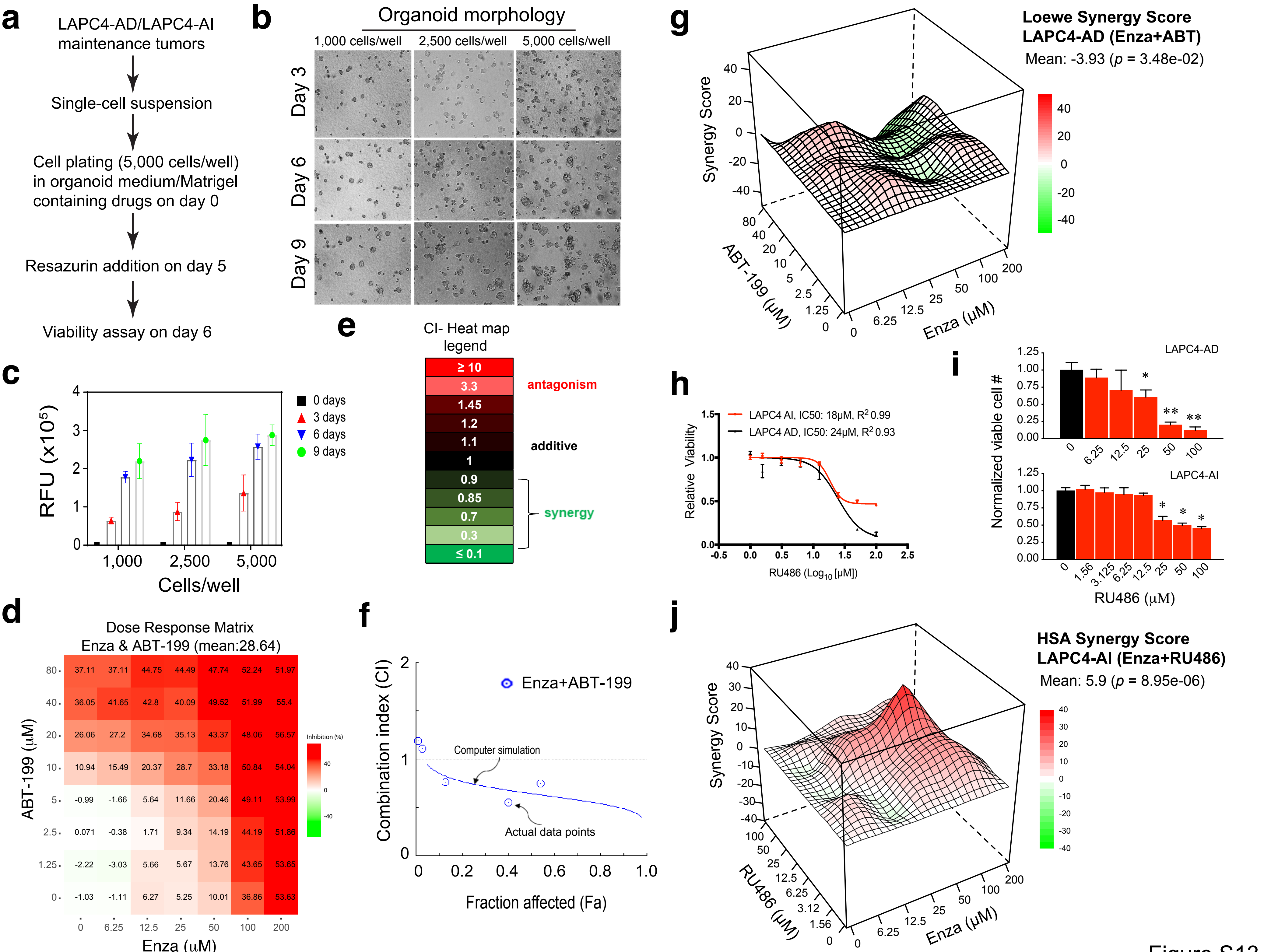
a
LAPC4-AD/LAPC4-AI maintenance tumors
Single-cell suspension
Cell plating (5,000 cells/well) in organoid medium/Matrigel containing drugs on day 0
Resazurin addition on day 5
Viability assay on day 6
b
Organoid morphology
1,000 cells/well
2,500 cells/well
5,000 cells/well
Day 3
Day 6
Day 9
c
RFU (x10$^5$)
Cells/well
1,000
2,500
5,000
0 days
3 days
6 days
9 days
d
Dose Response Matrix
Enza & ABT-199 (mean:28.64)
ABT-199 (μM)
80: 37.11 37.11 44.75 44.49 47.74 52.24 51.97
40: 36.05 41.65 42.8 40.09 49.52 51.99 55.4
20: 26.06 27.2 34.68 35.13 43.37 48.06 56.57
10: 10.94 15.49 20.37 28.7 33.18 50.84 54.04
5: -0.99 -1.66 5.64 11.66 20.46 49.11 53.99
2.5: 0.071 -0.38 1.71 9.34 14.19 44.19 51.86
1.25: -2.22 -3.03 5.66 5.67 13.76 43.65 53.65
0: -1.03 -1.11 6.27 5.25 10.01 36.86 53.63
Enza (μM): 0 6.25 12.5 25 50 100 200
Inhibition (%)
e
CI- Heat map legend
≥ 10
3.3
1.45
1.2
1.1
1
0.9
0.85
0.7
0.3
≤ 0.1
antagonism
additive
synergy
f
Combination index (CI)
Fraction affected (Fa)
Enza+ABT-199
Computer simulation
Actual data points
g
Loewe Synergy Score
LAPC4-AD (Enza+ABT)
Mean: -3.93 (p = 3.48e-02)
Synergy Score
ABT-199 (μM)
Enza (μM)
h
LAPC4 AI, IC50: 18μM, R$^2$ 0.99
LAPC4 AD, IC50: 24μM, R$^2$ 0.93
Relative Viability
RU486 (Log$_{10}$ [μM])
i
LAPC4-AD
LAPC4-AI
Normalized viable cell #
RU486 (μM)
j
HSA Synergy Score
LAPC4-AI (Enza+RU486)
Mean: 5.9 (p = 8.95e-06)
Synergy Score
RU486 (μM)
Enza (μM)

Figure S13

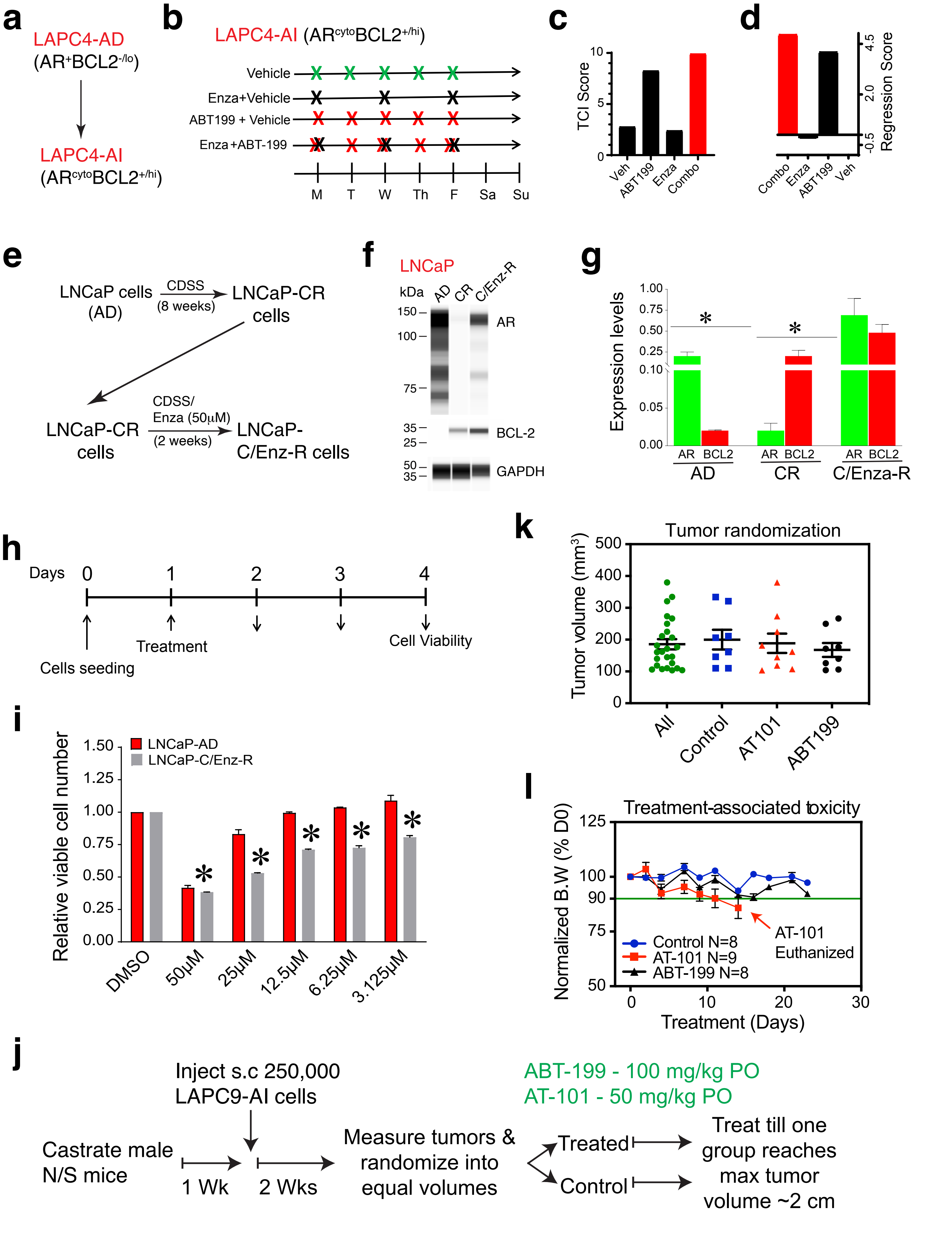

a
LAPC4-AD
(AR+BCL2-/lo)
LAPC4-AI
(ARcytoBCL2+/hi)
b
LAPC4-AI (ARcytoBCL2+/hi)
Vehicle
Enza+Vehicle
ABT199 + Vehicle
Enza +ABT-199
M T W Th F Sa Su
c
TCI Score
10
5
0
Veh
ABT199
Enza
Combo
d
Regression Score
4.5
2.0
-0.5
Combo
Enza
ABT199
Veh
e
LNCaP cells (AD)
CDSS (8 weeks)
LNCaP-CR cells
CDSS/ Enza (50µM) (2 weeks)
LNCaP-C/Enz-R cells
f
LNCaP
kDa
AD
CR
C/Enz-R
150
100
75
35
25
50
35
AR
BCL-2
GAPDH
g
Expression levels
1.00
0.75
0.50
0.25
0.10
0.05
0.00
*
AR BCL2
AD
CR
C/Enza-R
h
Days
0 1 2 3 4
Cells seeding
Treatment
Cell Viability
i
Relative viable cell number
1.50
1.25
1.00
0.75
0.50
0.25
0.00
LNCaP-AD
LNCaP-C/Enz-R
DMSO
50µM
25µM
12.5µM
6.25µM
3.125µM
k
Tumor randomization
Tumor volume (mm3)
500
400
300
200
100
0
All
Control
AT101
ABT199
l
Treatment-associated toxicity
Normalized B.W (% D0)
125
100
90
75
50
Control N=8
AT-101 N=9
ABT-199 N=8
AT-101 Euthanized
0 10 20 30
Treatment (Days)
j
Inject s.c 250,000 LAPC9-AI cells
ABT-199 - 100 mg/kg PO
AT-101 - 50 mg/kg PO
Castrate male N/S mice
1 Wk
2 Wks
Measure tumors & randomize into equal volumes
Treated
Control
Treat till one group reaches max tumor volume ~2 cm


Figure S14

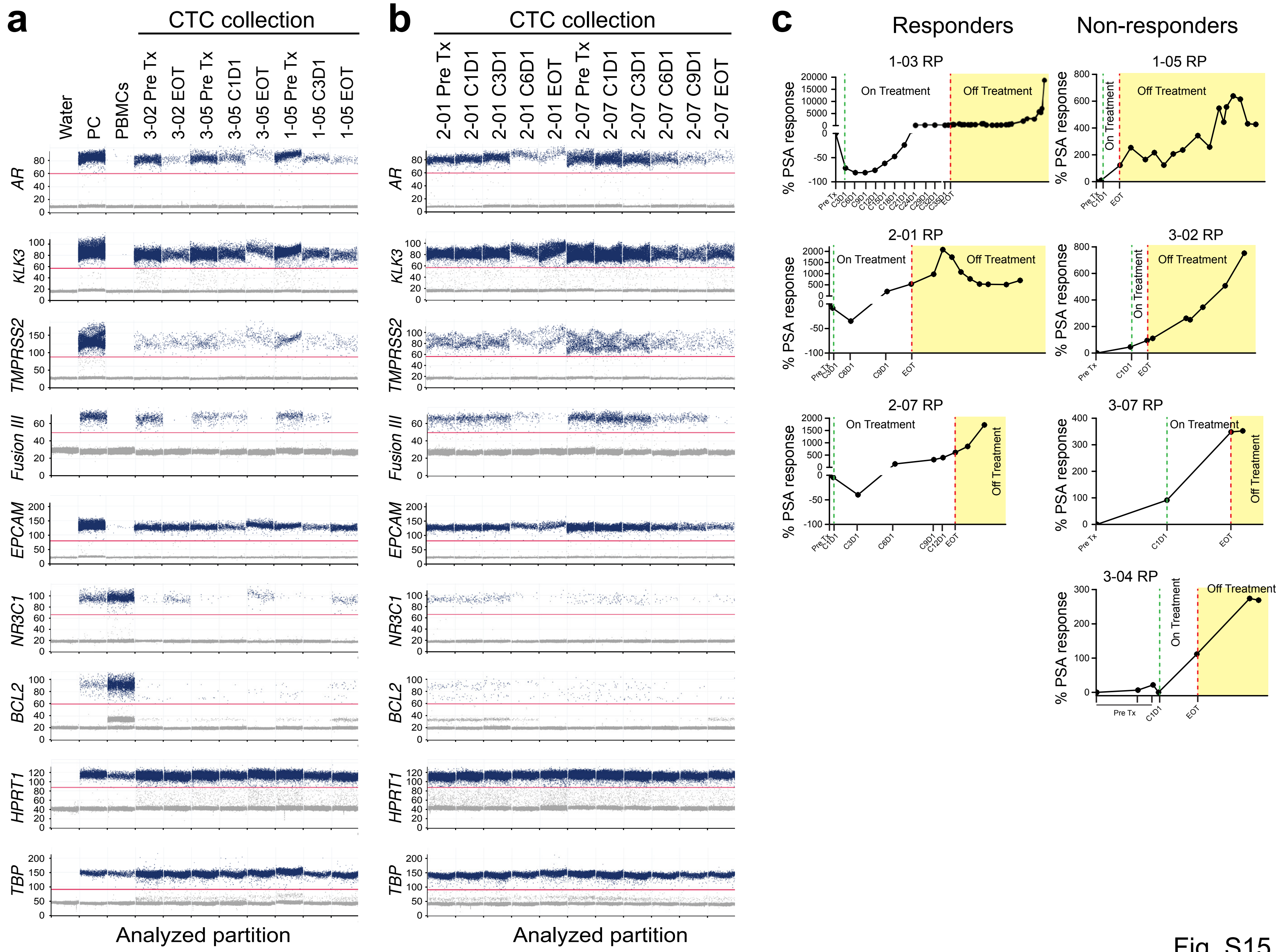
a
CTC collection
Water
PC
PBMCs
3-02 Pre Tx
3-02 EOT
3-05 Pre Tx
3-05 C1D1
3-05 EOT
1-05 Pre Tx
1-05 C3D1
1-05 EOT
AR
KLK3
TMPRSS2
Fusion III
EPCAM
NR3C1
BCL2
HPRT1
TBP
Analyzed partition
b
CTC collection
2-01 Pre Tx
2-01 C1D1
2-01 C3D1
2-01 C6D1
2-01 EOT
2-07 Pre Tx
2-07 C1D1
2-07 C3D1
2-07 C6D1
2-07 C9D1
2-07 EOT
Analyzed partition
c
Responders
Non-responders
1-03 RP
1-05 RP
2-01 RP
3-02 RP
2-07 RP
3-07 RP
3-04 RP
On Treatment
Off Treatment
% PSA response

Fig. S15
